# A genomic diversity atlas of North American frogs reveals geographic hotspots and coldspots of intraspecific variation

**DOI:** 10.64898/2026.03.13.711368

**Authors:** Lisa N. Barrow, Chris X. McDaniels, Anusha P. Bishop, Luis Amador, Daniele L. F. Wiley, Celina M. Eberle, Noelle M. Mason, Esteban O. Rosario Sanchez, J. Tomasz Giermakowski, Eric A. Hoffman, Gregory F. M. Jongsma, Moses J. Michelsohn, Maria Nava Martinez, Kadie N. Omlor, Sophia R. Rivera, Ariadna S. Torres López, Alexa R. Warwick, Kathleen C. Webster

**Affiliations:** Museum of Southwestern Biology and Department of Biology, University of New Mexico; Albuquerque, New Mexico, 87131, United States; Department of Environmental Science, Policy, and Management, University of California, Berkeley; Berkeley, California, 94720, United States; Department of Biology, University of Central Florida; Orlando, Florida, 32816, United States; Department of Natural History, New Brunswick Museum; New Brunswick, E2K 1E5, Canada; Department of Fisheries and Wildlife, Michigan State University; East Lansing, Michigan, 48824, United States

**Keywords:** extended specimens, museum collections, latitudinal diversity gradient, biodiversity hotspots, amphibians, ddRAD sequencing

## Abstract

Genetic variation is a key component of population resilience and thus an essential tool for wildlife management. For most species, however, we lack information about their current range-wide genomic diversity and how historical and contemporary factors have shaped those patterns. To address these gaps, we built new georeferenced genomic datasets for 2,481 individuals from 46 frog species to develop a genomic diversity atlas representing 43% of species native to the United States and Canada. We identified geographic “hotspots” of high genomic diversity and “coldspots” of low genomic diversity across species and analyzed potential explanations for these patterns. Our results show that several species have their lowest diversity at northern latitudes, at range edges, or within fragmented populations, suggesting areas in need of careful monitoring. More than half (56.8%) of species exhibited a latitudinal gradient in genomic diversity, but we found little evidence for an association between genomic diversity and human disturbance. Furthermore, discordant latitudinal patterns among species suggest that a combination of factors, including range center-edge dynamics and range fragmentation, interact to produce variable diversity patterns. Our study provides a general framework for mapping range-wide genomic diversity of multiple species to support genetically informed biodiversity conservation.

---

Genetic diversity within species is essential to adaptation and survival in changing environments^1,2^, and it is a key target under the Convention on Biological Diversity’s Kunming-Montreal Global Biodiversity Framework^3^. The logistical challenges of documenting and explaining genetic diversity patterns within and among species, however, remain daunting. High-quality tissues and genome-scale data do not exist for most populations and taxonomic groups^4,5^, and methods to interpret population genomic data for many species in a comparative framework are still emerging^6,7^. Natural history tissue repositories, high-throughput genotyping, and novel approaches for analyzing spatial genomic patterns provide the samples, data, and framework needed to understand this critical component of biodiversity.

Biodiversity hotspots are commonly used to prioritize areas for conservation based on criteria such as species richness, endemism, and rarity^8^. More recently, multispecies genetic diversity hotspots have been considered in some regions^9–12^, but incorporating genetic information into conservation planning at broad scales remains challenging. Synthesizing existing data is more cost-effective than generating new sequences, but this approach requires substantial data curation because most genetic sequences in public repositories are not linked to geographic coordinates^13–15^. Furthermore, much of the available genetic information from natural populations consists of mitochondrial or chloroplast DNA, which represents a single history and may not reflect the intraspecific variation relevant for biodiversity conservation^16,17^. Previous efforts to uncover the spatial distribution of genetic variation at a global scale have also highlighted important taxonomic and regional gaps in existing data^18^. Additional georeferenced, genome-wide datasets for underrepresented taxa are therefore needed to characterize the spatial distribution of genomic diversity and incorporate genetic information into conservation planning.

In addition to determining where genomic variation occurs, understanding how historical and contemporary factors influence that variation, and why species exhibit discordant patterns, are important goals in evolutionary biology. Decades of phylogeographic studies focused on one or few species at a time have provided valuable insights into regional biogeographic patterns, but sampling differences often make comparisons in a common framework difficult^19,20^. Macrogenetic studies reanalyzing repurposed data from thousands of species have shown that mitochondrial variation is associated with geographic attributes such as latitude, range size, and climate across eukaryotes^21,22^, insects^23^, mammals^18,24^, and amphibians^18,24,25^. The availability and representation of single-locus data have often required analyses at coarse scales that could not account for species identity or regional differences^24^. New reduced-representation genomic sequencing datasets^26^ make it increasingly feasible to analyze genomic diversity within and across species, providing opportunities to assess the consistency of range-wide patterns and identify species-specific discrepancies that may be important for managing vulnerable taxa.

Amphibians are the most threatened vertebrate class^27^ and would benefit from further genomic study, but progress has been hindered by their large, complex genomes^28^. Here, we present 46 georeferenced, genome-scale datasets for frogs sampled across the eastern two-thirds of North America. We harness these data to build a genomic diversity atlas and address consequential goals in both conservation and evolutionary biology. First, we generated continuous maps of genomic diversity for each species and combined these to identify “hotspots” of high diversity and “coldspots” of low diversity across multiple species that can be used to inform conservation priorities. Second, we test whether latitude or human disturbance, two major factors previously associated with genomic diversity^18^, explain the spatial distribution of intraspecific variation while considering differences among species and regions. Finally, we investigate the predictability of these range-wide diversity patterns based on geographic range and biological characteristics. Overall, our study provides a general framework for mapping and interpreting spatial genomic diversity within and across species that is extendable to other taxa and regions.

## Results and Discussion

### New Population Genomic Datasets for 46 Frog Taxa

We sampled 46 species or species complexes from >690 localities through field collections and loan requests from natural history museums and research labs (Supplementary Table 1). Our field efforts involved coordination with 30 state wildlife resource agencies and more than 90 landowners to obtain permits and land access. Fieldwork conducted from 2022 to 2023 yielded >2,100 multi-part, high-quality specimens archived at the Museum of Southwestern Biology, University of New Mexico (UNM) that are searchable via Arctos^29^. Tissue loans from 25 natural history collections and genomic DNA extracts from two research labs provided an additional 1,113 samples. Our dataset represents five of 10 frog families and 43% of extant species native to the United States and Canada, focused on species with broad geographic distributions. These species are currently ranked “Least Concern” by the International Union for Conservation of Nature^30^, but establishing baseline levels of genomic variation in wild populations is a necessary step for monitoring and preventing biodiversity loss^3^.

We generated genome-wide, population-level data within species by sequencing 2,688 samples with double-digest restriction site-associated DNA sequencing^26^ at the University of Wisconsin-Madison Biotechnology Center DNA Sequencing Core Facility following established protocols. Raw reads were demultiplexed, filtered, and assembled using resources at the UNM Center for Advanced Research Computing and the ipyrad v. 0.9.102 toolkit^31^. Since most species in the dataset lack reference genomes^28^, we conducted a series of *de novo* assemblies and quality control steps to identify and remove outliers, misidentified samples, potential hybrids, and samples with poor sequencing performance. For each species, we then evaluated assemblies and generated datasets using different sequence clustering and missing data thresholds to assess the robustness of downstream results to assembly parameter choice. Full methodological details are provided in Supplementary Information.

Our final datasets included 2,481 samples and an average of 54 (6 to 132) samples per species, with more samples and localities for species with larger ranges. We assembled an average of 5.27 x 10^6^ (4.21 x 10^5^ to 2.91 x 10^7^) filtered reads per sample. Datasets requiring a minimum of 50% of samples to retain a locus included an average of 17,514 (4,742 to 30,640) loci per species, and those requiring a minimum of 80% of samples to retain a locus included an average of 8,840 (1,794 to 17,300) loci per species. Species with the fewest loci retained were the Spring Peeper *(Pseudacris crucifer*), which includes multiple deeply divergent lineages^32^, and spadefoot toads in the family Scaphiopodidae, which are estimated to have the smallest genome sizes among the species included in our study^28^. The large number of loci and inclusion of two samples for most localities provide sufficient variation to describe population-level diversity patterns within each species^33^.

### Maps of Frog Genomic Diversity on a Continental Scale

We leveraged these new data and a recently developed method for calculating continuous maps of genomic diversity, *WINGEN*^34^, to visualize spatial patterns of genomic variation within each species. This method uses a moving window to calculate genetic diversity for each cell on a raster, followed by kriging to generate a smooth map of diversity, and finally masking to the area of interest, such as the sampled range for each species. Observed heterozygosity (H_O_) values ranged from 0.006 to 0.22, but these are not directly comparable across species since each dataset includes a different set and number of loci^35^. Instead, we compared the variation in H_O_ and the spatial patterns that emerged across species.

Species in eastern North America exhibited several consistent spatial patterns of genomic diversity (Fig. 1). First, many species had higher diversity in eastern localities compared to western localities, often with a more than two-fold decrease in H_O_ from east to west (e.g., *Rana grylio*, *Hyla femoralis*, and *Gastrophryne carolinensis*). Second, several species had lower diversity in northern localities compared to central or southern localities (e.g., *Acris crepitans*, *Pseudacris ocularis*, and *Anaxyrus quercicus*), demonstrating patterns consistent with both the central-marginal range hypothesis and post-glacial range expansion^36–38^. Third, species with disjunct ranges (e.g., *Scaphiopus holbrookii*, *Hyla andersonii*, and *Hyla gratiosa*) had reduced diversity in smaller range fragments compared to larger, more continuous areas. These results were robust to different assembly parameters except for species with more homogeneous H_O_ values, such as *Anaxyrus terrestris* (Supplementary Figs. 1–46).

**Fig. 1.**
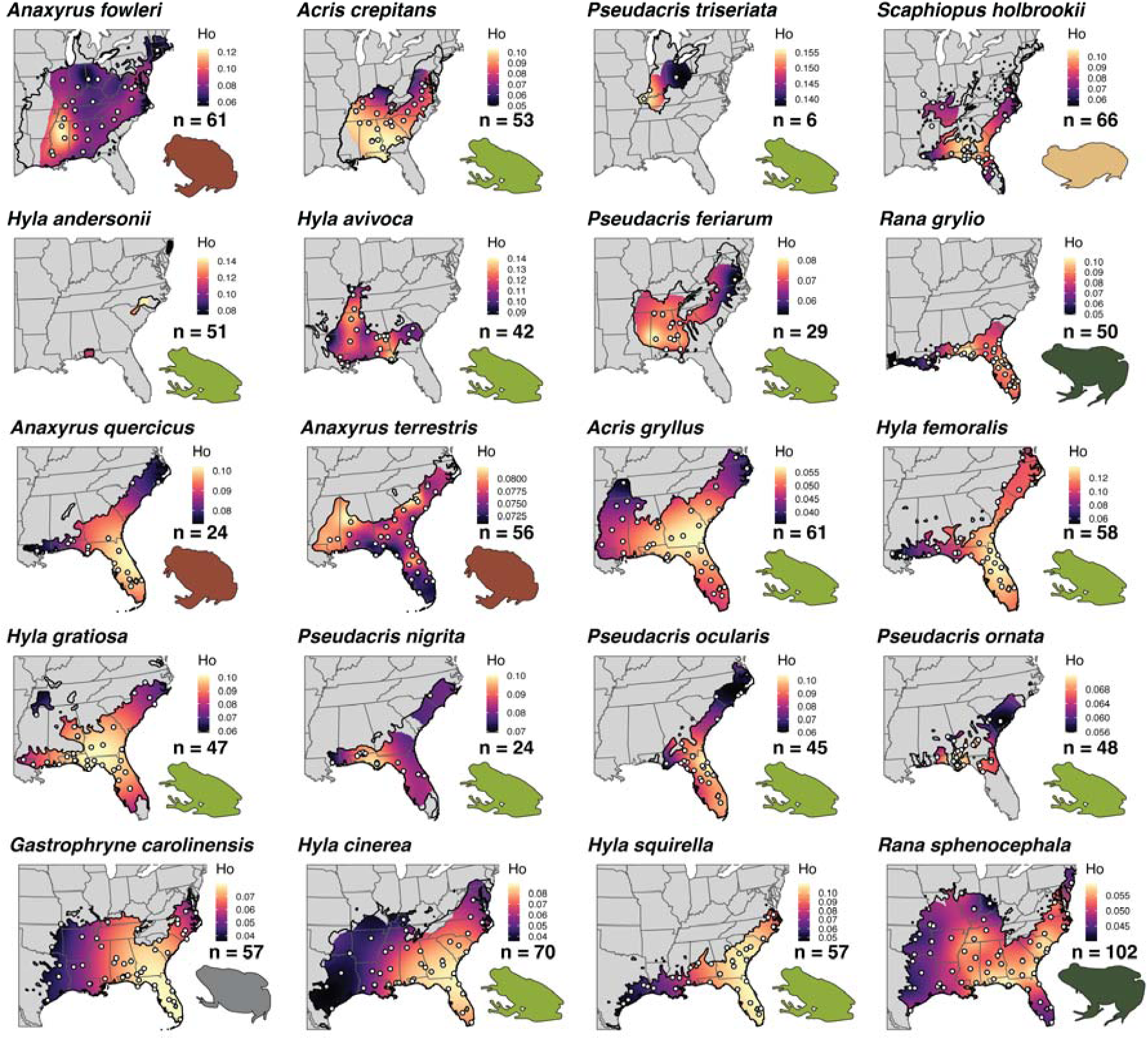
Maps of genomic diversity for 20 frog species in eastern North America. Continuous maps of observed heterozygosity (H_O_) are shown from lower values (cooler colors) to higher values (warmer colors) and were masked to the sampled portion of the species’ range. Maps were generated from assemblies using the 50% minimum samples per locus threshold and the optimal clustering threshold for each species. Sample localities are shown as white dots except for *H. andersonii*, where they would obscure the map. The total number of samples (n) is reported for each species. Species binomials follow AmphibiaWeb taxonomy^79^ and range maps (bold black outlines) are based on those maintained by Travis W. Taggart (cnah.org). Frog icons were generated by the authors and are colored by taxonomic family: Bufonidae (brown), Hylidae (light green), Microhylidae (gray), Ranidae (dark green), and Scaphiopodidae (tan). **Alt text**: Maps depicting sample localities and levels of genomic diversity within each of 20 frog species sampled across eastern North America. Each map includes the sample size, range map, and a color scale that represents observed heterozygosity interpolated across the study area.

Species with northern distributions displayed high variation in H_O_ and several of the same spatial patterns (Figs. 2 and 3B). The two species we evaluated with the widest distributions, *Rana pipiens* and *Rana sylvatica*, displayed stark declines in diversity from east to west, which has been reported in previous phylogeographic studies using traditional genetic markers^39,40^. Multiple species had lower diversity in northeastern localities and at range edges compared to southern and central localities (e.g. *Anaxyrus americanus*, *Pseudacris crucifer*, and *Rana catesbeiana*). The disjunct southwestern range of *Rana palustris* had much lower diversity compared to the eastern part of the range, again highlighting the potential consequences of fragmentation for the maintenance of genomic diversity^2,38^.

**Fig. 2.**
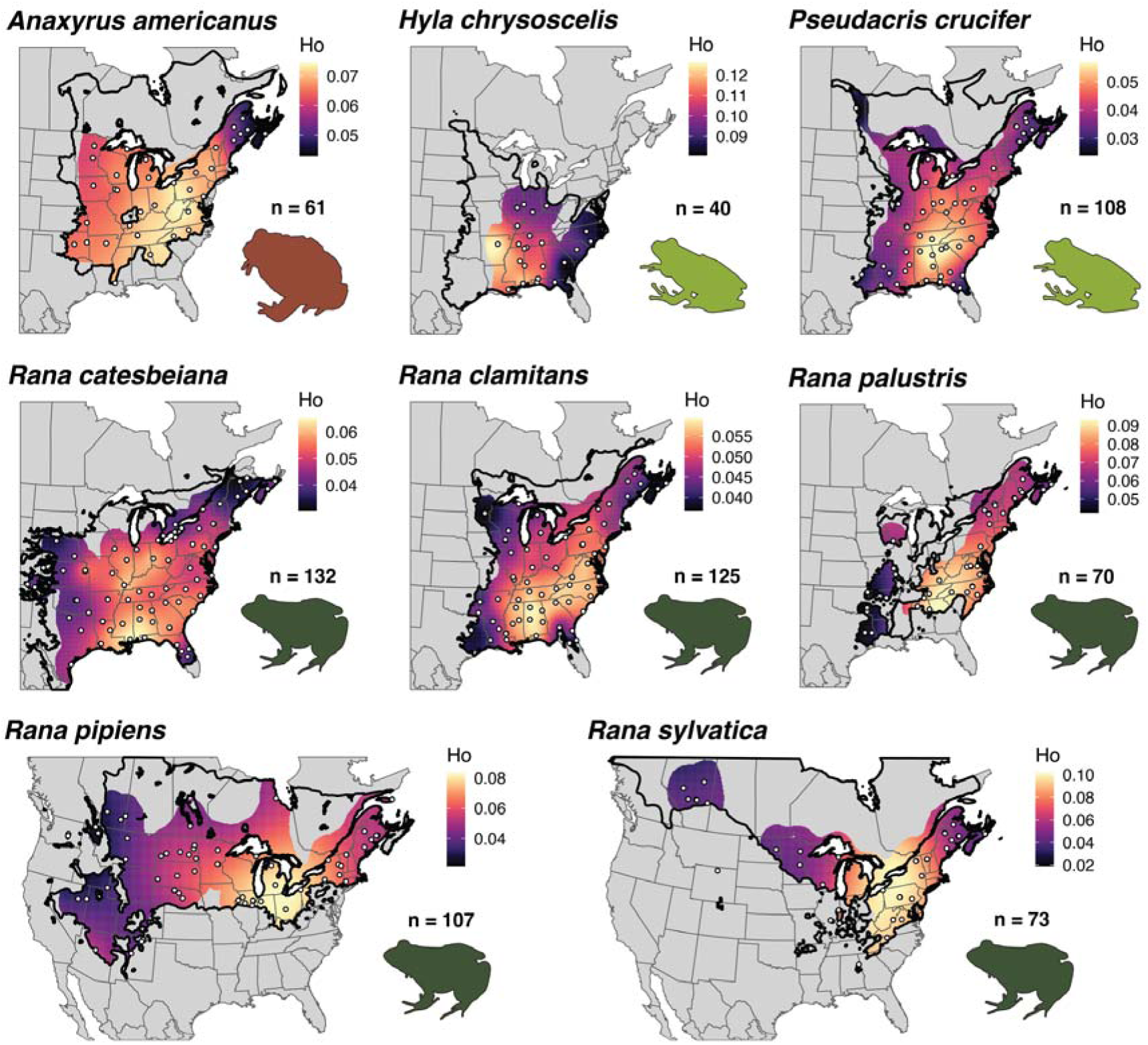
Maps of genomic diversity for eight frog species with northern distributions in North America. Continuous maps of observed heterozygosity (H_O_) are shown from lower values (cooler colors) to higher values (warmer colors) and were masked to the sampled portion of the species’ range. Details as described in Fig. 1. **Alt text**: Maps depicting sample localities and levels of genomic diversity within each of eight frog species sampled across eastern North America. Each map includes the sample size, range map, and a color scale that represents observed heterozygosity interpolated across the study area.

**Fig. 3.**
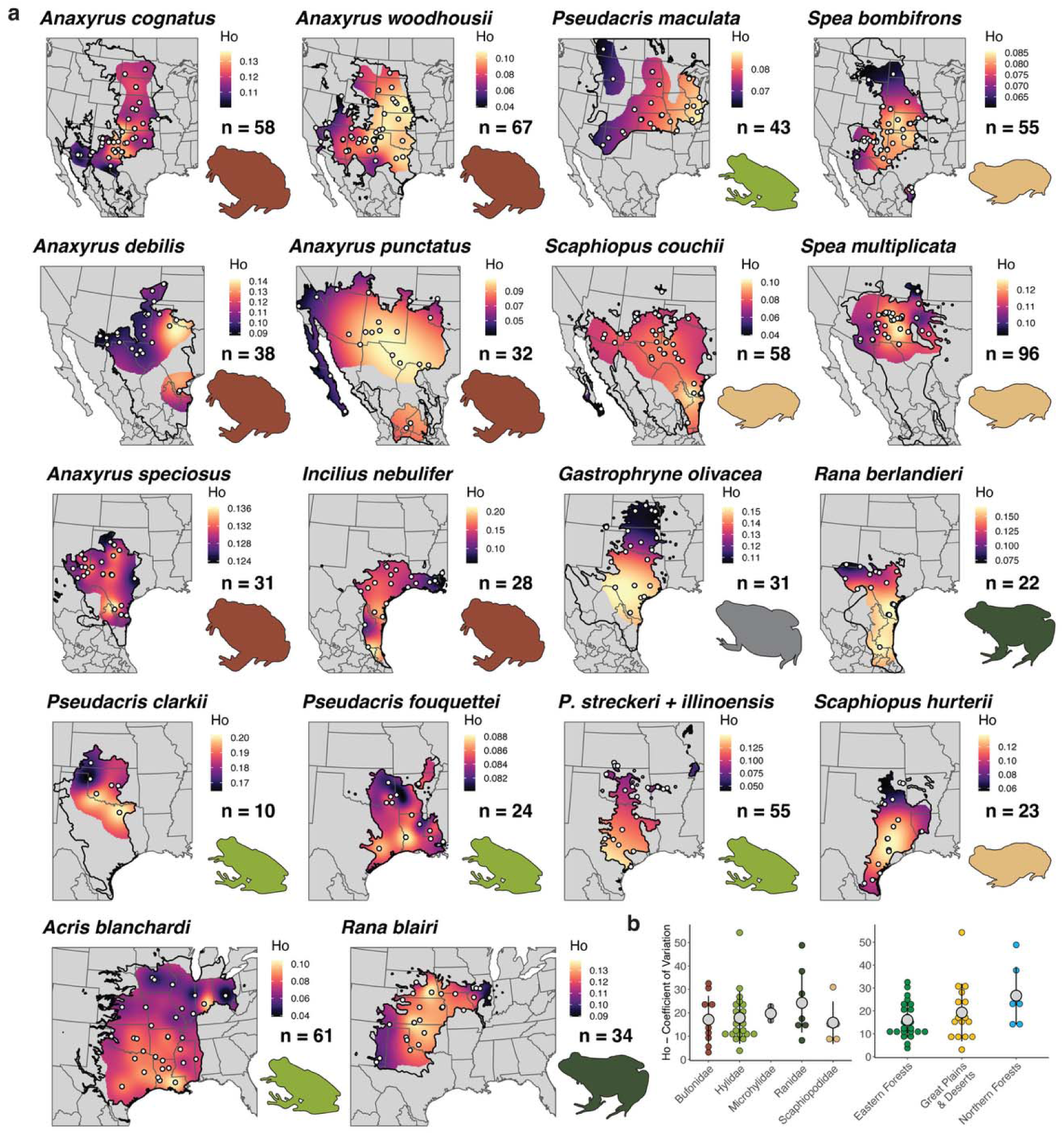
Maps of genomic diversity for 18 species in central North America and summary of heterozygosity across 46 species. (**a**) Continuous maps of observed heterozygosity (H_O_) for 18 frog species with distributions in the Great Plains and North American Deserts. Sample localities are shown as white dots except for *P. illinoensis*, where they would obscure the map. Other details as described in Fig. 1. (**b**) Coefficient of variation in H_O_ by taxonomic family and by ecoregion. Large gray circles and bars represent means and two standard deviations. **Alt text**: Panel (**a**) shows maps depicting sample localities and levels of genomic diversity within each of 18 frog species sampled across central North America. Each map includes the sample size, range map, and a color scale that represents observed heterozygosity interpolated across the study area. Panel (**b**) shows graphs of the variation in observed heterozygosity for each species either grouped by taxonomic family or by ecoregion.

Species in central and western North America exhibited less consistent spatial patterns overall, perhaps because few species in this region share similar distributions (Fig. 3). Some species had patterns consistent with a latitudinal gradient of genomic diversity, with lower diversity in northern localities compared to southern localities (e.g., *Acris blanchardi*, *Gastrophryne olivacea*, and *Rana berlandieri*). Two species with ranges extending to the Baja California Peninsula, *Scaphiopus couchii* and *Anaxyrus punctatus*, exhibited their lowest diversity in this region. The disjunct distribution of *Pseudacris illinoensis* had a more than two-fold reduction in diversity compared to the broader range of *Pseudacris streckeri*. These maps provide a valuable resource for understanding the current distribution of genomic diversity within species, highlight key patterns and the variability of H_O_ across geographic distributions, and lay the foundation for testing hypotheses about predictors of genomic diversity. Importantly, they can continue to be improved with finer-scale sampling and exploration of additional diversity metrics in the future.

### Building Composite Maps Across Species

Resources for conserving biodiversity are limited, thus finding opportunities that maximize conservation benefits across multiple habitats and species are important to consider^9,41,42^. We sought to identify shared regions of the highest and lowest genomic diversity across species to highlight areas that could be prioritized for preserving this important component of biodiversity. We used the maps of genomic diversity generated above and investigated different thresholds for identifying areas with the highest and lowest diversity within each species. We created binary maps with more restrictive (smaller area) to less restrictive (larger area) thresholds based on 5%, 10%, and 20% quantile values for the highest and lowest diversity areas for each species (Supplementary Figs. 47–92). For each threshold, we combined maps across species to visualize areas where multiple species have their highest or lowest diversity. We accounted for differences in species richness across the study region by generating maps based on the proportion of species that have their highest or lowest diversity in each area. We then applied global and local Getis-Ord *G* statistics to identify areas with significant clustering of high and low genomic diversity across species^43,44^.

### Hotspots of High Genomic Diversity

We identified consistent hotspots of high genomic diversity across multiple species in the southeastern U.S., particularly the Gulf Coast region of northern Florida, southwestern Georgia, and southeastern Alabama (Fig. 4 and Supplementary Figs. 93–94). At the least restrictive 20% threshold, 13 species had their highest diversity in that region (Fig. 4A), and even at the most restrictive 5% threshold, seven species shared their highest diversity there (Fig. S93A). Clustering analyses also identified western Virginia and surrounding areas (central Appalachian Mountains) and southern Texas with significant clustering of high diversity across multiple species (Fig. 4C). The southeastern U.S. remained significant when we examined the proportion of species with their highest diversity to account for its high species richness (Fig. 4, E and G and Supplementary Fig. S93, E and G). This region is known for its high concentration of contact zones and phylogeographic breaks for amphibians and other taxa, a pattern likely explained by maintenance of genetic diversity in glacial refugia^37,45,46^.

**Fig. 4.**
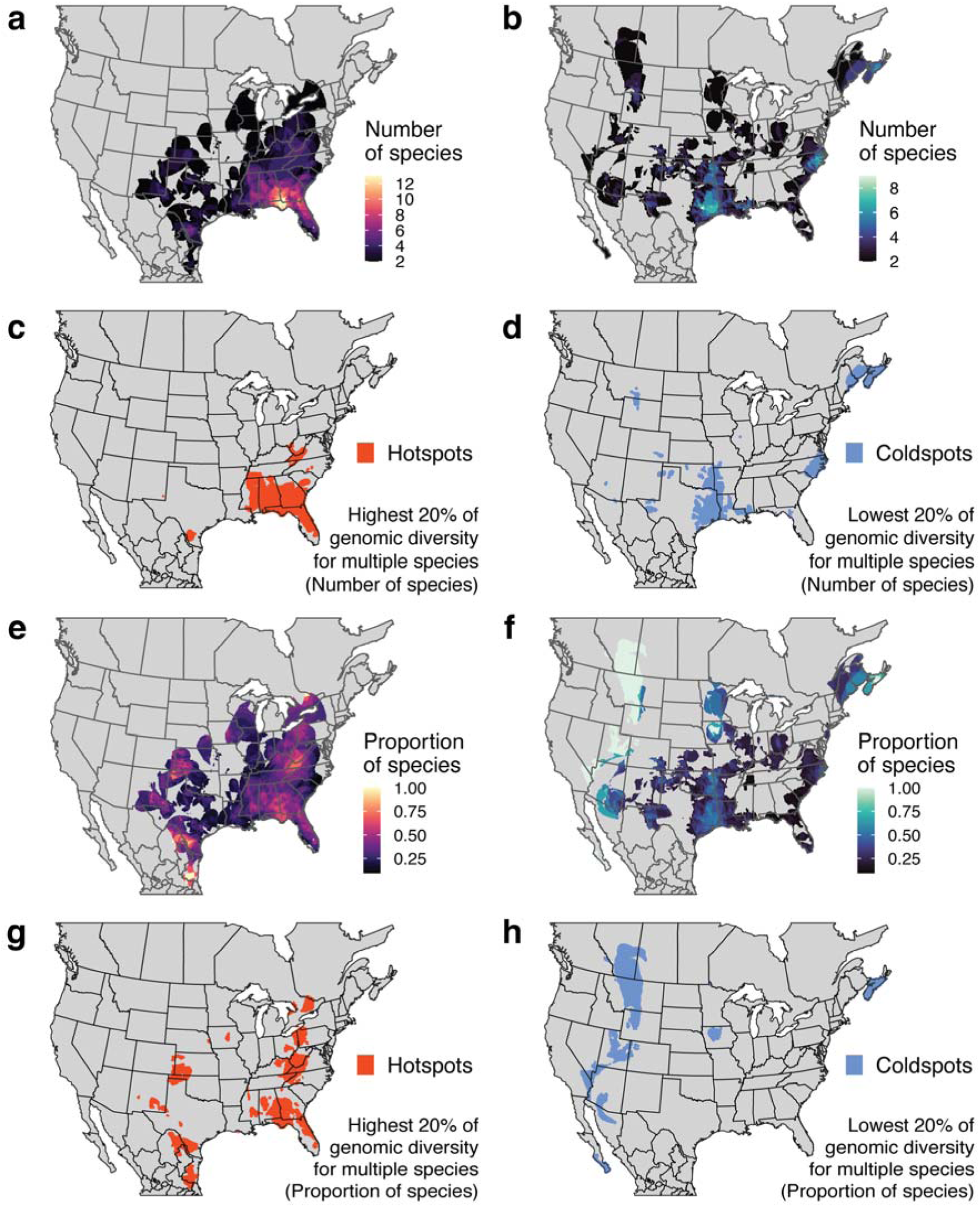
Composite maps of high (“hotspots”) and low (“coldspots”) genomic diversity across 46 species. (**a**, **b**) The color scale indicates the number of species in each grid cell with their highest (**a**) or lowest (**b**) 20% of genomic diversity. (**c**, **d**) Areas of significant clustering based on Getis-Ord Gi statistics where multiple species have their highest (**c**, orange) or lowest (**d**, blue) 20% of genomic diversity. (**e**, **f**) The color scale indicates the proportion of species in each grid cell with their highest (**e**) or lowest (**f**) 20% of genomic diversity. (**g**, **h**) Areas of significant clustering based on Getis-Ord Gi statistics where multiple species have their highest (**g**, orange) or lowest (**h**, blue) 20% of genomic diversity. **Alt text**: Composite maps displaying areas with either the highest (left column) or lowest (right column) 20% of genomic diversity across multiple species. The color scales depict either the number of species (**a**, **b**) or the proportion of species (**e**, **f**) with their highest or lowest diversity in each area. Areas of significant clustering based on Getis-Ord Gi statistics are shown in orange for hotspots (**c**, **g**) or blue for coldspots (**d**, **h**).

The greatest proportion of species exhibited their highest genomic diversity in northeastern Mexico (Fig. 4 and Supplementary Figs. 93–94). Three sampled species, *Incilius nebulifer*, *Rana berlandieri*, and *Scaphiopus couchii*, shared their highest diversity in this southern extent of their ranges (Supplementary Figs. 68, 79, and 88). Other regions with a large proportion of species with high diversity included southeastern New Mexico, western Kansas and Oklahoma, eastern Ohio and western Pennsylvania, and southern Ontario in Canada (Fig. 4G). Protecting such hotspots has the potential to preserve the most genomically diverse populations of multiple species. This approach would take an agnostic view of which variation is being preserved and how it relates to current and future environmental conditions, but it could maximize the variation preserved when resources are limited and assessments of adaptive potential for each species are not feasible.

### Coldspots of Low Genomic Diversity

We identified less consistent coldspots of low genomic diversity depending on whether we accounted for species richness or not (Fig. 4 and Supplementary Figs. 93 and 94). Up to nine species had their lowest diversity in eastern Texas (Fig. 4B), and this area had significant clustering for all thresholds examined (Fig. 4D and Supplementary Figs. 93D and 94D). When we examined maps based on the proportion of species with low diversity, however, eastern Texas was no longer identified as a coldspot, suggesting the high species richness in the southeastern U.S. and the pattern of decreasing diversity from east to west in several of those species explained this result. Instead, the northeastern U.S. and adjacent regions in New Brunswick and Nova Scotia, Canada emerged as a consistent region with low diversity across species for all thresholds except the most restrictive 5% threshold (Supplementary Fig. 94H).

The other coldspots we identified were consistent among thresholds but differed based on whether we accounted for species richness. The Atlantic Coast region of North Carolina and Virginia included multiple species with low diversity (Fig. 4B), but this region was no longer identified as a coldspot after accounting for species richness (Fig. 4F). Areas with a large proportion of species with low diversity included Baja California Sur, northern Utah, central Montana, and southern Alberta, Canada (Fig. 4F). These patterns of low diversity could result from a combination of historical processes, such as post-glacial range expansion leading to decreased diversity in recently colonized areas^37,39^, and contemporary factors, such as smaller populations being supported at less suitable range edges^38^. In species such as *Rana pipiens*, these regions are associated with a major range contraction and documented population declines^47,48^, although some evidence suggests these peripheral populations had depauperate diversity before range contraction occurred^49^. These regions of low diversity suggest populations in need of close monitoring to preserve the genomic variation that remains, regardless of the cause.

### Latitudinal Gradients of Intraspecific Genomic Diversity

The Latitudinal Diversity Gradient is a well-described biogeographic pattern of increasing species richness from higher latitudes to the Equator^50^. More recently, macrogenetic studies have demonstrated similar patterns of higher genetic diversity at lower, more equatorial latitudes for several taxonomic groups, but the nature of these datasets has precluded assessments of within-species gradients and differences among species^18,22,24^. We leveraged our new, population-level genomic data to investigate the relationship between latitude and genomic diversity within each species and determine how universal the latitudinal gradient in genomic diversity is among frogs of eastern North America. Within species, we predicted a negative association between latitude and genomic diversity. We calculated observed heterozygosity (H_O_) for each individual and averaged H_O_ values among samples with the same coordinates.

We compared results from both Spearman correlation tests^51^ and spatial regression models^52^ to account for spatial nonindependence of the samples. Several species, 25 out of 44 (56.8%), had the expected negative correlations between latitude and H_O_, but the correlation strength varied substantially and was not always robust to corrections (Fig. 5A and Supplementary Table 2). After a Benjamini-Hochberg correction^53^, we found 19 of 44 (43.2%) species with significant Spearman correlations and 9 of 44 (20.5%) species with significant relationships between latitude and H_O_ based on spatial regressions (Fig. 5A). We found one species, *Anaxyrus terrestris*, with a significant positive correlation because southern populations in peninsular Florida had lower diversity than more northern populations (Fig. 1 and Supplementary Fig. 11). Our results demonstrate that the latitudinal gradient in genomic diversity is a common, but not universal pattern within North American frogs.

**Fig. 5.**
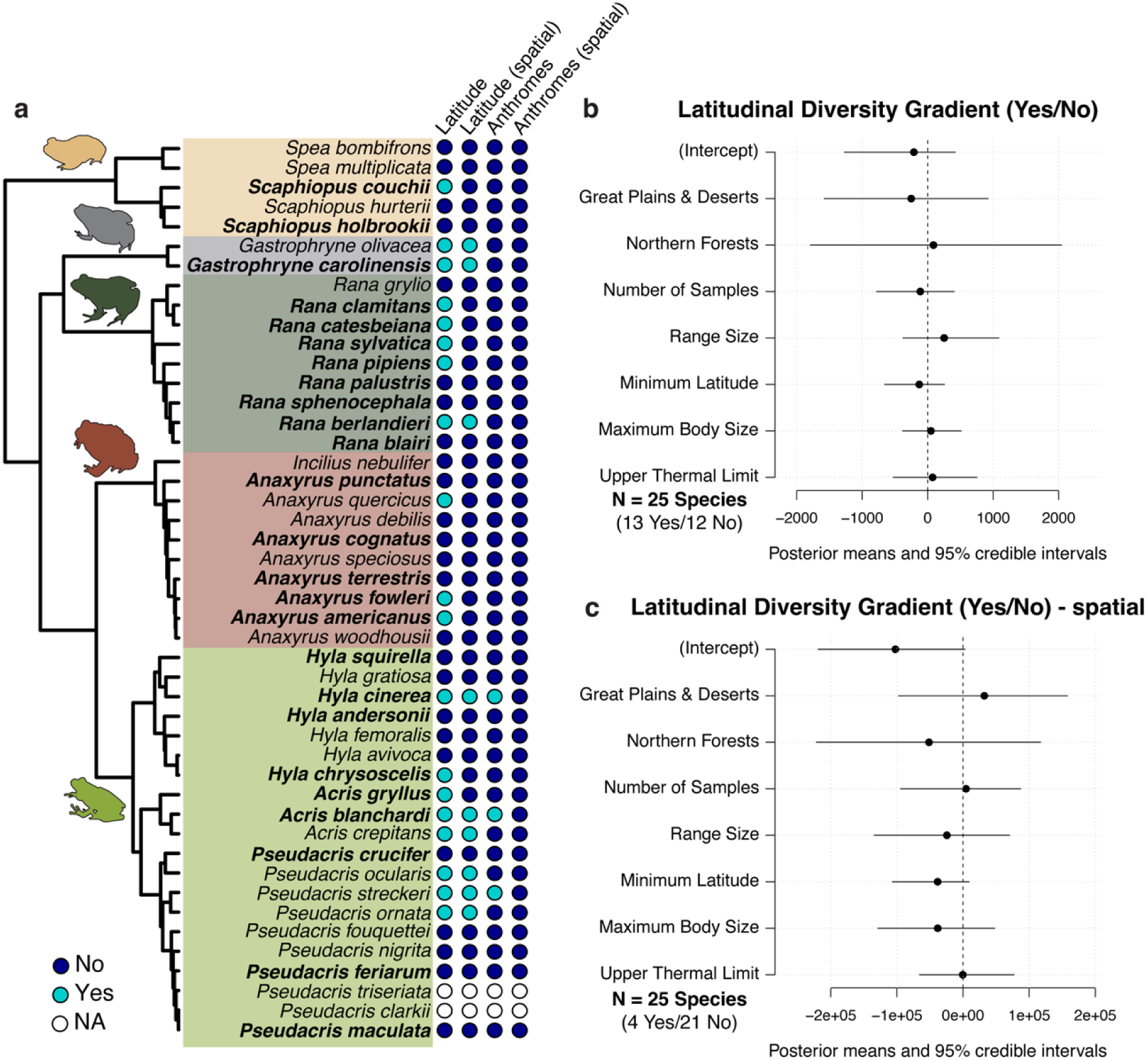
Associations between intraspecific diversity and latitude or human disturbance. (**a**) Species with (Yes, light blue) or without (No, dark blue) significant associations between genomic diversity and latitude or anthromes. Results from Spearman correlations or spatial regressions (spatial) are shown after correction for multiple tests. Phylogenetic relationships are based on Jetz and Pyron^78^ and species listed in bold have thermal tolerance information from Pottier et al.^59^. Tests were not conducted on *P. triseriata* or *P. clarkii* because of limited sample sizes. (**b**, **c**) Model results from *MCMCglmm* testing potential predictors of whether species show a latitudinal gradient in genomic diversity based on (**b**) Spearman correlations or (**c**) spatial regressions. For ecoregions, effects are relative to Eastern Temperate Forests. The 95% credible intervals all overlap with 0, indicating no predictors are significant. **Alt text**: Graphs showing the phylogenetic relationships of the study species, whether each species shows a significant association between intraspecific diversity and latitude or human disturbance, and model results testing predictors of whether species show a latitudinal gradient in genomic diversity.

### Human Disturbance and Intraspecific Genomic Diversity

Human activities in the last two centuries have had a profound influence on wildlife populations and the distribution of biological diversity^54^. Evidence for the expected relationship of lower genetic diversity with increasing human disturbance has been inconsistent^7,18,55^. We leveraged the History Database of the Global Environment database (HYDE v3.5^56^) to test for a positive association between H_O_ and ‘anthromes’ classified from most disturbed (e.g., 11 Urban) to least disturbed (e.g., 63 Wild, remote). Following a similar approach as above, we conducted these tests within each species to determine whether genomic diversity has been detectably influenced by human activities for any species in our study.

Few species, 8 of 44 (18.2%), showed correlations between H_O_ and human disturbance in the expected direction, but these were relatively weak and not robust to corrections (Supplementary Table 2 and Fig. 5A). After correction for multiple tests, only three taxa (*Acris blanchardi*, *Hyla cinerea*, and *Pseudacris streckeri-P. illinoensis* complex) had significant Spearman correlations and none were significant based on spatial regressions. Despite documented population declines in some of these species and predicted genomic diversity losses^7^, several plausible explanations could account for the lack of an association with human disturbance. One issue is that only populations persisting after substantial environmental change can be sampled, excluding those that have already been extirpated or reduced. Relatedly, the full range of anthromes is not represented within each species. The expected lag time between recent disturbance and detectable changes in genetic variation further complicates assessments of these patterns^57^.

Finally, the species included in this study may also have been relatively resilient to human-mediated environmental changes thus far, since all are currently ranked “Least Concern”^30^.

### Predictability of Intraspecific Genomic Diversity Patterns

The presence of a latitudinal gradient in genomic diversity in several, but not all species examined raises the question: Is this pattern predictable based on species characteristics or sampling biases? We used phylogenetic comparative models^58^ to test factors related to sampling effort (number of samples), geographic distribution (range size, minimum latitude, and primary ecoregion), and biological characteristics (maximum body size, thermal tolerance) as predictors of whether species showed a latitudinal gradient in genomic diversity (Yes/No) and as predictors of the correlation strength (Spearman’s rho) between latitude and H_O_. We obtained upper thermal limits from AmphiTherm, an amphibian thermal tolerance database^59^, prioritizing CTmax values from adults with the same experimental endpoint where possible. Thermal data were available for only 25 of the 44 species, thus we built full models including the first five predictors for all species and reduced models including 25 species with all six predictors.

None of the factors we included were significant predictors of the latitudinal gradient of genomic diversity within species (Fig. 5 and Supplementary Figs. S95 and S96), indicating that latitude is associated with genomic diversity for species with a variety of geographic distributions and biological characteristics. The lack of this simple pattern in many other species can likely be explained by a combination of factors, such as range center-edge dynamics or range fragmentation resulting in genomic diversity levels that reflect local population abundances regardless of latitude^2,36,38^. Our results contribute to growing efforts to understand range-wide genomic diversity patterns within species and, like other systems^36,60^, suggest these complex patterns may be largely unpredictable. If species’ genomic diversity patterns are idiosyncratic, then we must continue to survey genomic variation within any species we aim to understand and conserve. Our study takes an important step towards this goal by providing 46 new frog datasets and establishing a framework to evaluate spatial patterns of diversity within and across species.

## Methods

### Sampling

We obtained samples from 46 species or species complexes through fieldwork and museum loans, aiming to sample two individuals per site separated by 50–100 km across the distribution of each species. Frogs were captured under appropriate state and local permits and were processed following protocols approved by the University of New Mexico Institutional Animal Care and Use Committee (UNM IACUC 20-201006-MC and 23-201375-MC). Additional tissues and genomic DNA extracts were requested via loan from natural history collections and research laboratories. We included 2,688 samples from the United States, Canada, and Mexico, collected from the years 1975–2024, with most samples (∼91%) collected in the last two decades. Additional details are provided in Supplementary Information and Supplementary Table 1.

### Sequence Data Collection and Processing

We extracted genomic DNA from liver, toe, muscle, or blood samples using the E.Z.N.A.^®^ Tissue DNA Kit (Omega Bio-tek, Norcross, GA) following the manufacturer’s protocols. Sample concentrations were quantified using a Qubit™ Broad Range Quantification Assay Kit (Life Technologies Corporation, Eugene, OR) and quality was assessed via agarose gel electrophoresis. Samples were sent to the University of Wisconsin Biotechnology Center DNA Sequencing Core Facility (RRID:SCR_017759) for double-digest restriction-site associated DNA (ddRAD) library preparation and sequencing^26^.

We processed raw sequence data using the ipyrad v. 0.9.102 toolkit^31^ and resources from the UNM Center for Advanced Research Computing. Full details of our assembly process and downstream analyses are provided in Supplementary Information. Briefly, we demultiplexed and filtered raw reads, then conducted a series of *de novo* assemblies and quality-checking steps. Our workflow included initial assemblies with multiple species at the genus or subgenus level to perform checks for outliers, misidentified samples, potential hybrids, and samples with poor sequencing performance. For these multi-species assemblies, we set the clustering threshold to 0.85 and the minimum samples per locus to 4, and then conducted exploratory analyses—a Pearson principal component analysis (PCA), a sparse non-negative matrix factorization (sNMF) analysis^61^, and a distance-based split network in SplitsTree App v6.4.11^62^—to verify species identities and identify problematic samples to remove (details in Supplementary Table 1). After correcting or removing samples, we generated assemblies for each species dataset that tested clustering thresholds from 0.85–0.97 and evaluated several metrics to identify the “optimal” threshold, following a modified workflow from^63^. We repeated downstream analyses with assemblies using either the “optimal” clustering threshold or the same threshold for each species, and the minimum number of samples required to retain a locus of either 50% or 80% to determine consistency of results based on different assembly parameters and levels of missing data.

### Mapping Genomic Diversity Within Species

We generated continuous maps of genetic diversity for each species using the *WINGEN* R package v2.2.0^34^. We calculated observed heterozygosity (H_O_) from datasets with one random SNP per locus using the moving window function, in which each window encompassed samples within 150 km. Additional diversity metrics including nucleotide diversity and allelic richness were considered, but we opted to use H_O_ because it is less sensitive to low sample sizes and rarefaction^34^. To create a smooth diversity map, we used the kriging function to a final interpolated resolution of 10 km. We then masked the kriged layer by the sampled range for each species to create our final maps. Range maps were obtained from the Center for North American Herpetology (https://www.cnah.org/; accessed 24 September 2025).

### Identifying “Hotspots” and “Coldspots” of Genomic Diversity Across Species

Binary rasters for each species were generated using the *terra* R package^64^ by computing the upper quantile for each threshold (the highest or lowest 5%, 10%, and 20% of genomic diversity) across cells and setting cell values greater than or equal to the threshold to 1 and all other values to 0. We generated composite maps across all species by either summing the binary rasters together to visualize the number of species with their highest or lowest diversity in each cell, or by calculating the proportion of species within each cell that had their highest or lowest diversity to account for differences in species richness. For each composite raster, we calculated the global Getis-Ord *G* statistic to determine whether high or low diversity areas clustered significantly in geographic space using the *spdep* R package^44,65^. Finally, we used a local G test in the *sfdep* R package to identify and visualize the locations of “hotspots” of high diversity and “coldspots” of low diversity^43,66^.

### Testing for a Latitudinal Diversity Gradient Within Species

We calculated individual-based H_O_ within each species from datasets including one random SNP per locus using the *dartRverse* R package^67,68^. We again opted to use H_O_ (calculated as the proportion of heterozygous loci per individual) and then averaged across individuals because it accounts for missing data^67^ and is less sensitive to small sample sizes than other metrics such as allelic richness. We assessed and removed outliers, calculated average H_O_ per locality, and tested for correlations between H_O_ and latitude using Spearman correlations^51^. We also used spatial regression models to investigate the relationship between latitude and H_O_ while accounting for spatial autocorrelation. We fit generalized least squares (GLS) models with exponential, spherical, and gaussian correlation structure in the *nlme* R package^69,70^ and spatial error and lag models using functions in the *spatialreg* R package^71,72^. We compared spatial models and the original GLS model and selected the model with the lowest Akaike Information Criterion (AIC) value^73^. Finally, we plotted model residuals in space and computed Moran’s I using the *spdep* R package^74^ to confirm whether the final model reduced spatial autocorrelation relative to the original model.

### Testing the Influence of Human Disturbance on Genomic Diversity

We obtained human disturbance information from the History Database of the Global Environment (HYDE v3.5^56^) and extracted the associated ‘anthrome’ for each sample based on collection year (1975–2024) using the classification system of ^75–77^. Anthromes range from most disturbed (e.g., 11 Urban) to least disturbed (e.g., 61–63 Wild, remote). Following the same approach described for latitude above, within each species, we tested for a positive relationship between H_O_ and anthromes using Spearman correlations and spatial regression models. To account for multiple tests, we applied a Benjamini-Hochberg correction^53^ with a false discovery rate of 0.05 to adjust the p-values from each test.

### Phylogenetic Comparative Methods

We fit phylogenetic generalized linear mixed models using *MCMCglmm*^58^ to assess whether predictors related to sampling effort, geographic distribution, or biological characteristics explain why some species exhibit a latitudinal gradient in genomic diversity while others do not. We tested model sets with binary responses (Yes/No) based on results from Spearman correlations and spatial regressions or a continuous response (Spearman rho). We built full models with 44 species and five predictors (number of samples, sampled range size, minimum latitude, primary ecoregion, maximum body size) and models with a reduced dataset including six predictors (adding thermal tolerance) and the 25 species with data available in the AmphiTherm database^59^. Phylogenetic relationships were obtained from Jetz and Pyron^78^. We set priors of *V* = 1, nu = 0.02 for the phylogenetic random effect and ran each model across four chains of 5,000,000 iterations with a burn-in of 1,000,000 and thinning every 4,000 steps. We inspected trace plots, Gelman diagnostic statistics, and effective sample sizes to evaluate convergence.

## Data and code availability

All collected specimens and parts were archived at the Museum of Southwestern Biology, are available via loan request, and are searchable via the Arctos collection management system; Supplementary Table 1). All sequence data are available through the NCBI Sequence Read Archive (SRA BioProject Number PRJNA1423219). Additional data files, R scripts, and output files are available from Dryad (10.5061/dryad.x95x69q0k). All other data are available in the main text or Supplementary Information.

## Author contributions

L.N.B., C.X.M., and A.P.B designed research; L.N.B., C.X.M., L.A., D.L.F.W, C.M.E., N.M.M., and E.O.R.S analyzed data; L.N.B. supervised research and wrote the paper; and all authors curated data, performed research, and reviewed the paper.

## Acknowledgments

For tissues and genomic DNA samples, we thank D. Kizirian and S. Katanova (AMNH), L. Ammerman and K. Graves (ASNHC), D. Laurencio and D. Warner (AUM), L. Scheinberg (CAS), V. de Brito, C. Dillman, and C. Dardia (CUMV), J. Roberts, L. Wilson, W. Stark, and T. W. Taggart (FHSM), C. Phillips and E. Santoyo Brito (INHS), A. Motta (KU), G. Pauly and N. Camacho (LACM), D. Dittman, G. Thom, and C. Austin (LSUMZ), T. LaDuc and D. Cannatella (TNHC), M. Campbell (MSB), C. Spencer, J. McGuire, and R. Tarvin (MVZ), A. Bremner (NBM), B. Stuart and J. Beane (NCSM), L. Steger and K. Yule (NEON), J. Braun, R. Hawkins, C. Siler, and J. Watters (OCGR), N. Cairns (RAM), A. Lathrop, C. Dutton, and J. Basseches (ROM), L. Fitzgerald and T. Hibbitts (TCWC), P. Soltis, T. Lott, and C. Sheehy III (UF), G. Pandelis (UTA), V. Zhuang, C. Lieb, and J. Johnson (UTEP), S. Birks (UWBM), G. Watkins-Colwell and D. Skelly (YPM), E. M. Lemmon, and S. Lougheed. For permitting, land access, and field support, we thank B. Kappel, C. Threadgill, T. Carter, G. Scull, D. Beaty, A. Scott, A. Cochran, S. Baxter, K. Irwin, A. Williams, E. Shackleton, M. Kiser, M. Knight, C. Iversen, M. Klassen, R. Slyter, E. Smith, M. Larson, C. Werner, S. Stipkovits, O. Wetsch, J. Darr, J. Bishop, J. Cook, D. Young, M. Petoskey, C. Pope, E. Dytrych, M. Jenkins, J. Freese, D. Turner, D. Schutt, M. Massatt, E. McCook, C. Hughes, C. Barrow, A. Barrow, J. Hawkins, D. Rambo, L. Kring, J. Sparks, D. Mixon, B. Howze, L. Petercheff, R. Winstead, K. Klingenberg, M. Van Scoyoc, M. Huston, D. Beard, S. Klueh-Mundy, R. Smith, J. Duguay, D. Yorks, J. Swift, C. Reitz, H. Cyr, S. Peyton, C. Vaughn, J. Thomas, S. Whittington, K. Hanson-Dorr, W. Meriwether, D. Elsen, J. Moree, B. Holder, C. Masley, K. Watkins, J. Briggler, S. Dunn, L. Mee, P. Stringer, P. Isakson, R. Meissner, K. Wecker, K. Schultes, K. Omlor, M. Howery, D. Nichols, C. Deal, E. Kearse, D. J. Morris, D. Miller, B. Davis, C. McCorkle, B. Kendall, E. Dowd Stukel, R. Boles, B. Anderson, M. Pons, J. White, J. Gunnels, D. Gissel, D. Dittmer, A. Candelaria, J. Kart, S. Dressler, R. Francis, and J. Kitchell. For administrative and research support, we thank J. Kavka, T. Lam, K. Madrid, J. Knight, T. Clayshulte, J. Kuestner, P. Johnston, J. Wilson, J. L. Tyler, S. Siegrist, B. Yazzie, and E. Gyllenhaal. We thank the UNM Center for Advanced Research Computing for providing the high-performance computing and large-scale storage resources used in this work and T. W. Taggart for providing range maps. ChatGPT-4o was used to assist with writing some of the R scripts for this work.

