## Supplementary Information for "A genomic diversity atlas of North American frogs reveals geographic hotspots and coldspots of intraspecific variation"

**This file includes:**

Extended Methods

Supplementary Figs. 1 to 96

Supplementary References

**Other supporting materials for this manuscript include:**

Supplementary Tables 1 and 2 (Excel files)

Extended Methods

**Sample collection.** Samples from 46 frog species or species complexes were obtained through a combination of field collections and museum loans. We focused our sampling on all broadly distributed species in eastern North America, which represent five of the 10 frog families and 43% of the extant species native to the United States and Canada. We conducted fieldwork in the United States primarily from 2022-2023 with the goal of sampling two individuals per site from sites separated by approximately 50-100 km and distributed across the geographic range of each species. This sampling design was chosen to provide population-level estimates of genomic diversity across each species’ range, assuming localities at these distances would be unlikely to have frequent migration among them given the limited dispersal ability of frogs. We aimed to sample at least 20 localities per species (with more localities for species with larger ranges) and sequenced two individuals per locality to represent population history with thousands of loci across the genome^1–3^.

We obtained permits and landowner permission from 30 state wildlife resource agencies and more than 90 national forests, state forests, wildlife management areas, and private landowners. These include the **Alabama** Department of Conservation and Natural Resources, Tuskegee National Forest (NF), Talladega NF, Bankhead NF; **Arkansas** Game and Fish Commission; **Florida** Fish and Wildlife Conservation Commission, Ocala NF, Picayune Strand State Forest (SF), Blackwater River SF, Point Washington SF, Lake Talquin SF, Jennings SF, Goethe SF, Withlacoochee SF, Little Big Econ SF, Lake Wales Ridge SF, Matanzas SF, Seminole SF, Tate’s Hell SF, Pine Log SF, Watermelon Pond Wildlife and Environmental Area (WEA), Babcock Webb Wildlife Management (WMA), Lake Wales Ridge WMA, Hilochee WMA, Everglades WMA, Salt Lake WMA, Three Lakes WMA, Hungryland WEA, Suwannee River Water Management District (WMD), Southwest Florida WMD; **Georgia** Department of Natural Resources, Chattahoochee-Oconee NF, Tallulah Gorge WMA, Lake Russell WMA, Altamaha WMA, Altama Plantation WMA, Ceylon WMA, Richmond Hill WMA, Ohoopee Dunes WMA, Canoochee Sandhills WMA, Alapaha River WMA, Dawson Forest WMA, Alligator Creek WMA, Grand Bay WMA, Elmodel WMA, Chickasawhatchee WMA, Albany Nursery WMA, Sandhills WMA, Chattahoochee Fall Line WMA; **Indiana** Department of Natural Resources, Hoosier NF, Jasper-Pulaski Fish and Wildlife Area; **Iowa** Department of Natural Resources; **Kansas** Department of Wildlife and Parks, Wilson WMA; **Kentucky** Department of Fish and Wildlife Resources, Harris-Dickerson WMA; **Louisiana** Department of Wildlife and Fisheries, Kisatchie NF, Clear Creek WMA, West Bay WMA, Fort Polk WMA, J.C. Sonny Gilbert WMA, Boeuf WMA, Sandy Hollow WMA, Lake Ramsay WMA, Soda Lake WMA, Joyce WMA, Manchac WMA, Maurepas Swamp WMA; **Maine** Department of Inland Fisheries and Wildlife; **Maryland** Department of Natural Resources; **Michigan** Department of Natural Resources; **Minnesota** Department of Natural Resources; **Mississippi** Department of Wildlife, Fisheries and Parks, DeSoto NF, Tombigbee NF, Holly Springs NF, Homochitto NF, Bienville NF, Marion County WMA, Pearl River WMA, Caney Creek WMA, Tallahala WMA, O’Keefe WMA, Charles Ray Nix WMA, Upper Sardis WMA, Malmaison WMA, Nanih Waiya WMA, Wolf River WMA, Chickasawhay WMA, Mason Creek WMA; **Missouri** Department of Conservation; **Nebraska** Game and Parks Commission; **New Mexico** Department of Game and Fish; **New York** State Department of Environmental Conservation; **North Dakota** Game and Fish Department, Bureau of Land Management Schnell Recreation Area; **Ohio** Division of Natural Resources, Wayne NF; **Oklahoma** Department of Wildlife Conservation; **Pennsylvania** Fish and Boat Commission, State Game Lands 101, 314; **South Carolina** Department of Natural Resources, Francis Marion NF, Sumter NF, Sandhills SF, Poe Creek SF, Manchester SF, Wee Tee SF; **South Dakota** Department of Game, Fish, and Parks, Farm Island State Park; **Tennessee** Wildlife Resources Agency; **Texas** Parks and Wildlife Commission, LBJ and Caddo National Grasslands, Caddo WMA, Gus Engeling WMA; **Utah** Division of Wildlife Resources; **Vermont** Fish and Wildlife Department; **Virginia** Department of Wildlife Resources, Big Woods WMA, CF Phelps WMA, Dick Cross WMA, Fairystone Farms WMA, Mattaponi WMA, Merrimac Farm WMA, Powhatan WMA; and the **Wisconsin** Department of Natural Resources. Detailed locality information is provided in Supplementary Table 1.

Frogs were captured by hand and processed following protocols approved by the University of New Mexico Institutional Animal Care and Use Committee (UNM IACUC 20-201006-MC and 23-201375-MC). Upon capture, geographic coordinates in decimal degrees were recorded on handheld Garmin eTrex22x Handheld Navigator units. Individuals were euthanized with a two-step procedure involving an initial topical application of 20% benzocaine to the ventral surface. Once the individual was fully anesthetized and no longer exhibited righting behavior, sterile dissection scissors and forceps were used to remove vital organs. After removal of the heart, blood was collected in a microcapillary tube, used to prepare a thin blood smear, and the remainder was flash frozen in a liquid nitrogen dewar. Additional tissues including heart, lung, liver, thigh muscle, and toe (including skin) were flash frozen in liquid nitrogen. Blood smears were air-dried and fixed within 24 hours with 100% methanol; they were stained within two months with 10% Giemsa Blood Staining Solution (Avantor, Center Valley, PA). Specimens were preserved via injection of 95% ethanol, set in a standardized position, and later transferred to 70% ethanol for long-term storage. All specimens and parts were catalogued and archived at the Museum of Southwestern Biology, are available via loan request, and are searchable via the Arctos collection management system^4^ (arctos.database.museum; Supplementary Table 1).

Additional tissues and genomic DNA extracts were requested via loan from natural history collections and research laboratories. Samples were provided by the American Museum of Natural History (**AMNH**), Angelo State Natural History Collections (**ASNHC**), Auburn University Museum of Natural History (**AUM**), California Academy of Science (**CAS**), Cornell University Museum of Vertebrates (**CUMV**), Fort Hays State University Sternberg Museum (**FHSM**), Illinois Natural History Survey (**INHS**), University of Kansas Biodiversity Institute (**KU**), Los Angeles County Natural History Museum (**LACM**), Louisiana State University Museum of Natural Science (**LSUMZ**), University of New Mexico Museum of Southwestern Biology (**MSB**), University of California Berkeley Museum of Vertebrate Zoology (**MVZ**), New Brunswick Museum (**NBM**), North Carolina State Museum (**NCSM**), National Ecological Observatory Network (**NEON**), Oklahoma Collection of Genomic Resources (**OCGR**), Royal Alberta Museum (**RAM**), Royal Ontario Museum (**ROM**), Texas A&M University Biodiversity Research and Teaching Collections (**TCWC**), Texas Memorial Museum Texas Natural History Collection (**TNHC**), Florida Museum of Natural History Genetic Resources Repository (**UF**), University of Texas Arlington Amphibian and Reptile Diversity Research Center (**UTA**), University of Texas El Paso Biodiversity Collections (**UTEP**), University of Washington Burke Museum (**UWBM**), Yale Peabody Museum (**YPM**), E.A. Hoffman from University of Central Florida, and S.C. Lougheed from Queen’s University. Samples from the United States, Canada, and Mexico were included (Figs. 1-3) and these spanned the years 1975-2024, with most samples (~91%) collected in the last two decades (Supplementary Table 1).

**Library preparation and sequencing.** Sample preparation was conducted in a designated molecular lab with clean work surfaces (wiped with 10% bleach) and sterile, filtered pipette tips. We extracted whole genomic DNA from liver, toe, muscle, or blood samples using the Omega Bio-tek E.Z.N.A.^®^ Tissue DNA Kit (D3396, Omega Bio-tek, Norcross, GA) following the manufacturer’s protocols for either tissues (liver, toe, muscle) or whole blood. We performed the optional step to remove RNA using Invitrogen^™^ PureLink^™^ RNase A (Fisher Scientific, Denver, CO). We performed two elution steps with volumes of 75-200 µL each of Elution Buffer, depending on the starting quantity and freshness of the tissue, with lower volumes for older samples with smaller quantities of starting tissue. We quantified extractions using a Qubit™ Broad Range Quantification Assay Kit (Life Technologies Corporation, Eugene, OR) and either diluted samples to a concentration of 20 ng/µL with molecular grade water or concentrated samples on a ThermoSavant DNA120 SpeedVac^®^ Concentrator (Thermo Fisher Scientific, Waltham, MA) at 30 °C.

We arranged diluted samples (500 ng each) into 96-well plates and performed final checks of DNA quality via agarose gel electrophoresis. We prepared 1% agarose gels with 2.5 g Molecular Biology Grade Agarose (IBI Scientific, Dubuque, IA), 250 mL 10% TAE buffer (2 M Tris-Acetate, 0.05 M EDTA, pH 8.3; Omega Bio-tek, Norcross, GA), and 12.5 µL GelRed^®^ Gel Stain 10,000X in water (Biotium, Inc., Fremont, CA). We loaded 40 ng of DNA per sample mixed with an equal volume of 3X Gel Loading Dye, Purple (New England BioLabs, Ipswich, MA) in individual wells. We loaded 2 µL of Quick-Load Purple 1 Kb Plus DNA Ladder (New England BioLabs, Ipswich, MA) in the first and last well of each row as size standards. Gels were run at 80 V for 70 minutes on a Bio-Rad Horizontal Electrophoresis System and were imaged with a Bio-Rad GelDoc XR+ System.

Library preparation and sequencing were performed at the University of Wisconsin Biotechnology Center DNA Sequencing Core Facility (RRID:SCR_017759) using their Genotype by Sequencing (GBS) full-service option for double-digest restriction-site associated DNA (ddRAD) sequencing^5^. Samples were digested with the restriction enzymes MspI and SbfI-HF, and size selection was conducted on a PippinHT (Sage Science, Inc., Beverly, MA) targeting fragments of 300-450 base pairs (bp) in length. Sequencing was performed on the Illumina NovaSeq X Plus with a target of 4 million reads (2 x 150 bp) per individual (384 million reads per plate). Read quality was assessed for each plate via MultiQC v1.14^6^.

**Sequence assembly pipeline.** We processed raw sequence data using the ipyrad v. 0.9.102 toolkit^7^ and resources from the UNM Center for Advanced Research Computing (CARC). Raw reads for each plate were demultiplexed by individual barcodes ranging in length from 5-10 bp with 0 mismatches allowed. Sorted reads were filtered with default settings: maximum low-quality bases (parameter 9) = 5, Phred offset (parameter 10) = 33, strict filtering of adapters (parameter 16) = 2, minimum trimmed length (parameter 17) = 35, and no read trimming (parameter 25) = 0, 0, 0, 0. We then conducted a series of *de novo* assemblies and quality-checking steps including assemblies with paired and R1 reads only, ultimately choosing to work with only the R1 reads because those assemblies retained far more shared loci among individuals. As more reference genomes become available for these species in the future, we encourage others to explore the performance of reference-based assemblies with these data.

Our workflow included initial assemblies with multiple species at the genus or subgenus level to perform checks for outliers, misidentified samples, potential hybrids, and samples with poor sequencing performance. For these multi-species assemblies (27 total), we set the clustering threshold (parameter 14) at 0.85 and the minimum samples per locus (parameter 21) to 4. All other parameters were kept at default settings for ipyrad v. 0.9.102: minimum depth for statistical (parameter 11) and majority-rule (parameter 12) base-calling = 6, maximum clustering depth within samples (parameter 13) = 10000, maximum alleles per site in consensus (parameter 18) = 2, maximum uncalled bases (parameter 19) and heterozygotes (parameter 20) in consensus = 0.05, maximum SNPs per locus (parameter 22) = 0.2, maximum indels per locus (parameter 23) = 8, maximum heterozygous sites per locus (parameter 24) = 0.5, and all output formats produced (parameter 27).

For each multi-species assembly, we conducted a combination of exploratory analyses—a Pearson principal component analysis (PCA), a sparse non-negative matrix factorization (sNMF) analysis^8^, and a distance-based split network in SplitsTree App v6.4.11^9^—to verify species identities and identify potential problematic samples for removal. For PCA, we modified the STRUCTURE-formatted file including one randomly selected variable site per locus (‘.ustr’) to include a population column that assigned each sample to the field-identified species code. We then conducted a PCA using the “gl.pcoa()” function in the *dartR.base* R package^10,11^. We created interactive plots with the “plot_ly()” function in the *plotly* R package^12^ for each two-way combination of the first three PC axes and colored samples by their species code. We assessed every sample that did not group with its expected species to flag (1) misidentified samples (if they clearly grouped with a different species than their field-assigned identity; for samples that fell within the expected geographic range of both species and were morphologically similar, we verified voucher specimens when possible); (2) potential hybrids (if they were intermediate between species on the PCA plots and had a large number of reads and loci); and (3) samples with poor sequencing success (if they drifted towards the middle of the PCA plots and had less than 10% of the average number of loci for that species in the assembly).

To further verify species identities and assess admixture, we conducted sNMF analyses using the “snmf()” function in the R package *LEA*^13^. We converted the ‘.ustr’ file to ‘.geno’ format using the “struct2geno()” function in *LEA*, ran 100 repetitions each of the sNMF analysis setting the number of ancestral populations (K) from 1-10, and calculated the cross-entropy criterion for each run. We plotted the ancestry coefficients as pie charts to examine the “best” replicate (minimizing the cross-entropy value) of the K value that matched the number of species in the dataset. We verified the misidentified samples and noted any samples with mixed ancestry as potential hybrids if they had many loci or poorly sequenced samples if they had few loci.

As a final check for outliers, we opened the PHYLIP-formatted file (‘.phy’) in SplitsTree App v6.4.11^9^, obtained a distance matrix using the P Distance method^14^, and generated a split network using the Neighbor Net method^15,16^. We searched for each species prefix and set them to different colors to easily find misidentified samples and noted any samples that were intermediate between species (potential hybrids) or that had very long branches (poorly sequenced).

Misidentified samples were moved to their correct species folder for subsequent assemblies. Potential hybrids were kept with their original species for subsequent assemblies but were removed from downstream analyses to avoid the inflation of genomic diversity metrics because of mixed ancestry. Samples with poor sequencing success were dropped from subsequent assemblies and further analyses. All samples are included with these notes in Supplementary Table 1, and raw ‘.fastq’ files for all samples are uploaded to the NCBI Sequence Read Archive (SRA BioProject Number PRJNA1423219).

For each species dataset, we generated ipyrad assemblies that tested different values for the clustering threshold (parameter 14) and evaluated several metrics to identify the “optimal” threshold. This value determines how similar sequences should be to merge them into the same locus; a value too low can merge paralogous loci into the same locus, while a value too high can split naturally occurring allelic variation into multiple loci when they belong to the same locus. Our workflow was modified from McCartney-Melstad et al.^17^. We tested clustering thresholds from 0.85-0.97 and evaluated the following five metrics: (1) the correlation between pairwise divergence and pairwise missingness, (2) the slope of genetic divergence versus geographic distance, (3) the cumulative variance explained by the main principal components of genetic variation, (4) the total number of SNPs, and (5) heterozygosity. Metrics were generated and plotted using vcftools^18^ and the following packages in R^19^: *SNPRelate*^20^, *pheatmap*^21^, *grImport2*^22^, *ggplot2*^23^, and *scales*^24^.

For each metric, we recorded the value that was considered “optimal” following the interpretation of McCartney-Melstad et al.^17^; however, since the five metrics did not always agree on the same optimum, we chose the median value. Furthermore, since there was no clear optimum for some datasets, we repeated downstream analyses with assemblies based on two clustering thresholds for several species to determine how consistent results were when different assembly parameters were used. For each species, we chose the “optimal” value (hereafter, *clustOpt*) and then repeated analyses using the same value for all species (i.e., 93; hereafter, *clustSame*). To determine how consistent results were across datasets including different numbers of loci and amounts of missing data, we also generated datasets for each species setting the minimum number of samples per locus (parameter 21) to 50% of the samples for that species (hereafter, *min50*) and to 80% of the samples for that species (hereafter, *min80*).

**Mapping genomic diversity within species.** We generated continuous maps of genetic diversity for each species using the *WINGEN* R package v2.2.0^25^ (https://github.com/AnushaPB/wingen). *WINGEN* uses a moving window approach to calculate genetic diversity for each cell on a raster of the study region based on genetic data from the sampled individuals within the window surrounding and including that focal cell. Additional functions are used to create a smooth diversity surface via kriging and to mask the final map to a suitable background layer.

We used the ‘.vcf’ files output from ipyrad and randomly selected one SNP per locus using the Python script “vcf_single_snp.py” (https://github.com/pimbongaerts/radseq). We projected sample coordinates using the Conus Albers Equal Area projection (crs = 5070) and set the resolution of the background North America layer to 50 km based on our sampling distances. For each species, we used the range map from the Center for North American Herpetology (<https://www.cnah.org/>; accessed 24 September 2025; maintained by Travis W. Taggart) to create a background layer and set the extent to the range of that species. We manually edited range maps using QGIS v3.22.5^26^ when needed to encompass all sampled points and to exclude large unsampled portions of the range.

To run the moving window in *WINGEN*, we used the “window_gd()” function and set the window size to 3 (wdim = 3) to encompass 150 km, aiming to include at least two samples for calculating genetic diversity. We calculated average observed heterozygosity across all sites (stat = “Ho”) and set the minimum number of individuals to 1 (min_n = 1) to calculate diversity even if a window only included a single individual. To create a smooth map of genetic diversity, we used the “wkrig_gd()” function and disaggregated by a factor of 5 to achieve a final interpolated resolution of 10 km. We used the raster of sample counts per cell to weight the kriged prediction raster by sample size. Finally, we used the “mask_gd()” function to create a final continuous raster of genetic diversity that masked the kriged layer by the sampled range for each species. We used the following additional packages to read, project, and plot data: *vcfR*^27^, *ggplot2*^23^, *terra*^28^, and *sf*^29,30^.

**Identifying “hotspots” of high diversity and “coldspots” of low diversity across species.** One of our goals was to identify geographic areas of overlap across species that harbored either high or low genomic diversity (hereafter, “hotspots” or “coldspots”). We used different thresholds to summarize the areas of highest or lowest diversity within each species and then summarized across species to build composite maps as follows.

For each species, we generated binary maps to represent the area with the highest or lowest 5%, 10%, and 20% of genomic diversity for that species. We calculated the value for each threshold using the “global()” and “quantile()” functions in the *terra* R package and the final kriged, masked raster saved from *WINGEN*. The threshold was calculated using the probabilities (probs) of 95%, 90%, or 80%, and with NA values removed from the raster prior to calculation. Once the threshold value was determined, the binary raster was created by setting values greater than or equal to the threshold to 1 and all other values to 0. In other words, the 5% high diversity binary raster depicts the top 5% of grid cells to identify the areas with the highest observed heterozygosity for that species.

Once binary maps were generated for each species, we created composite maps across species at the 5%, 10%, and 20% thresholds for the highest diversity and lowest diversity. For each threshold, we first created composite maps by summing the binary rasters together to visualize the number of species with the highest or lowest diversity across the study region. These “sum of species” maps highlight common areas of high diversity or low diversity across multiple species. Second, because species richness varies across the study region, we also generated “proportion of species” maps to highlight areas with high or low diversity for the greatest proportion of species occurring in that area, regardless of the underlying species richness. To generate these maps, we divided each grid cell of the summed raster by the number of focal species that occur in that grid cell. We excluded grid cells that only included a single focal species from our final maps because our goal was to identify regions of high or low diversity across multiple species.

We applied global and local tests of spatial association to the composite rasters to determine whether regions of high and low genomic diversity across species clustered significantly in geographic space and, if so, to visualize the locations of significant clustering. First, we calculated the global Getis-Ord *G* statistic for spatial autocorrelation^31^ using the “global.Gtest()” function in the *spdep* R package^32^. We used this test to determine whether there was significant clustering of regions with high or low genomic diversity across the entire study area. Second, we employed a local G test^33^ with the “local_gi_perm()” function in the *sfdep* R package^34^ to identify and visualize the locations of potential “hotspots” of high diversity and “coldspots” of low diversity. We conducted analyses using both the “sum of species” and the “proportion of species” rasters for both high and low diversity. We exported final maps to visualize areas with significant clustering of species with high diversity (“hotspots”) or low diversity (“coldspots”) based on the classification of grid cells with gi > 0 and p-values from a folded permutation test of p < 0.05. We used the following additional packages to manipulate and plot data: *sf*^29,30^, *terra*^28^, *tidyterra*^35^, *ggplot2*^23^, *dplyr*^36^, and *tidyr*^37^.

**Testing for a Latitudinal Diversity Gradient within species.** Latitude is a commonly invoked correlate of *species* diversity on global and continental scales, typically referred to as the Latitudinal Diversity Gradient (LDG^38–41^). More recently, studies have shown similar patterns of increasing *intraspecific genetic* diversity with decreasing latitude on a global scale for various taxonomic groups^42–44^. Thus far, these studies have primarily been limited to the single locus, organellar genetic information available for thousands of taxa. Furthermore, these data are usually summarized across species by latitudinal bands or by grid cells at different resolutions. As Gratton et al.^44^ points out, these approaches “did not account for species identities, thus potentially conflating within-species gradients and differences among species occupying different regions.” Here, we leverage population-level nuclear data within species to investigate the relationship between latitude and genomic diversity, enabling us to investigate how universal the LDG in genomic diversity is across North American frogs and whether species-level attributes or geographic region explain the presence or absence of the LDG within species.

According to the LDG, we expect high diversity areas to occur at lower (more southern) latitudes and low diversity areas to occur at higher (more northern) latitudes within the northern hemisphere, resulting in a negative association between latitude and diversity. For each species, we calculated individual-based observed heterozygosity (Ho_ind) using the “gl.report.heterozygosity()” function in the *dartRverse* R package^10,11^. We compared values from the full datasets and those including one random SNP per locus (“.usnps”) and found they were highly correlated (correlation coefficients all > 0.8); we therefore used the “.usnps” dataset for subsequent analyses. We also performed a final check for outliers and tested for a correlation between H_O_ and the number of loci for each sample. We dropped any samples that had both extreme values of H_O_ and < 50% of the average number of loci for that dataset out of caution that the H_O_ values were influenced by the low number of loci rather than reflecting true patterns of biological diversity.

For samples from the same locality (with the exact same coordinates), we calculated the average observed heterozygosity per locality. We checked for normality in H_O_ within each species using histograms, quantile-quantile plots, and a Shapiro-Wilk test^45^. Since some datasets violated assumptions of normality, we tested for a correlation between H_O_ and latitude using a Spearman correlation^46^, recording the correlation coefficient (rho) and p-values for each species. Because spatial autocorrelation would violate the assumption of independence among samples, we also used spatial regressions to test the relationship between latitude and H_O_ when accounting for the spatial location of samples. We first computed Moran’s I autocorrelation coefficient of H_O_ using the “Moran.I()” function in the *ape* R package^47,48^. We created the weight matrix as the inverse of pairwise distances between samples, calculating the geodesic distances (fun = distGeo) between coordinate pairs projected in WGS84 with the “distm()” function of the *geosphere* R package^49,50^. We specified the alternative hypothesis as “greater” to test for positive autocorrelation.

We then used spatial regression models to investigate the relationship between latitude and H_O_ while incorporating spatially correlated error terms. We fit a series of generalized least squares (GLS) models with and without spatial correlation structure to identify and remove the effects of spatial autocorrelation. We fit an initial linear regression model (OLS) and plotted the residuals in geographic space to visualize the degree of autocorrelation. We then fit a variogram to the residuals, which measures the spatial dependence between observations as a function of distance, using the “variogram()” function in the *gstat* R package^51,52^. As an initial visualization of model fit, we examined exponential, spherical, and gaussian variogram models with the “fit.variogram()” function. We then fit GLS models with exponential, spherical, and gaussian correlation structure in the *nlme* R package^53,54^.

We also fit spatial error and lag models using functions in the *spatialreg* R package^55,56^. We created a distance-based neighbor list using the range of the best-fitted variogram, excluding any models that had convergence errors. Lagrange Multiplier diagnostics were used to test for spatial dependence in the original OLS model using the “lm.RStests()” function in the *spdep* R package^32^. We fit both spatial error (SEM) and spatial lag (SLM) models, compared the spatial models and the original GLS model based on Akaike Information Criterion (AIC), and selected the model with the lowest AIC value^57^ to assess the relationship between H_O_ and latitude after accounting for spatial autocorrelation.

Finally, we plotted model residuals in geographic space and computed Moran’s I of the residuals to confirm whether the final model reduced spatial autocorrelation relative to the original model. We used the “moran.test()” function in the *spdep* R package^58^. We created the spatial weights matrix by defining the upper limit of neighbor distances based on the range from the variogram (“dnearneigh()” function), using row standardized weights, and allowing weights of zero length for regions without neighbors in the specified distance (style = “W” and zero.policy = TRUE in the “nb2listw()” function).

**Testing the influence of anthropogenic disturbance on genomic diversity.** Previous studies have found mixed results regarding the possible association between intraspecific genetic diversity and human disturbance (Miraldo et al.^43^ and Millette et al.^59^ for global examples in animals with mitochondrial DNA; Exposito-Alonso et al.^60^ for examples in 20 plant and animal species with range-wide genomic data). Following a similar approach as above for latitude, we investigated the relationship between human disturbance and genomic diversity within North American frogs.

We obtained human population density and land use estimates from the History Database of the Global Environment (HYDE v3.5^61^) for the years 1970–2025. For each sample in our dataset, we extracted the associated ‘anthrome’ based on collection year (1975–2024) using the classification system of ^62–64^. The anthrome classes range from the most disturbed (e.g., 11 Urban) to the least disturbed (e.g., 61–63 Wild, remote). If human activity and disturbance levels are associated with wildlife population declines, we expect to find higher genomic diversity levels in areas with less human disturbance (i.e., higher anthrome values). Following the same approach described for latitude above, within each species, we tested for a positive relationship between H_O_ and anthromes using Spearman correlations and GLS models with and without spatial correlation structure.

We conducted 176 total tests for latitude and anthropogenic disturbance using both Spearman correlations and spatial regressions for each of the 44 species with at least n = 20 samples. To account for multiple tests, we applied a Benjamini-Hochberg correction^65^ to adjust the p-values from each test. We set a false discovery rate of 0.05 and report the original and adjusted results in Supplementary Table 2.

**Comparative analyses – What factors, if any, predict whether species support the LDG?** Few species had genomic diversity patterns associated with aspects of human disturbance, but 19 of 44 (43.2%) species exhibited patterns consistent with a Latitudinal Diversity Gradient (LDG) in genomic diversity after correction for multiple tests. We tested predictors in a phylogenetic comparative framework to assess whether factors related to sampling effort, geographic distribution, or biological characteristics explain why species do or do not support the LDG.

As response variables, we tested model sets with either a binary response (Yes/No) to examine predictors of whether a species supported the LDG, or a continuous response (correlation coefficients) to examine predictors of the strength of the relationship between latitude and H_O_. We constructed binary model sets based on the results from the standard correlation tests and the best spatial regression model for each species.

We included predictors related to sampling effort (number of samples), geographic distribution (sampled range size, minimum latitude, and primary ecoregion), and biological characteristics (maximum body size, thermal tolerance). We expected species with larger sample sizes and larger ranges would provide more power to detect an LDG pattern. We expected species with distributions at more northern latitudes and within the Northern Forests ecoregion would be more likely to exhibit an LDG pattern than those in southern latitudes and in other ecoregions (Eastern Temperate Forests, Great Plains, and North American Deserts). Body size was used as a proxy for dispersal ability, with the prediction that larger frogs would have higher dispersal and less clear signatures of an LDG pattern. We expected species with higher thermal tolerances might be more capable of rapid range shifts associated with historical climate change and would therefore be more likely to exhibit patterns consistent with the LDG.

Sampled range size was obtained by first generating a minimum convex polygon around the sample coordinates for each species using the “mcp()” function in the *adehabitatHR*^66^ R package. The area of the resulting polygon was then computed using the “areaPolygon” function of the *geosphere*^50^ R package. Species were assigned to ecoregions based on the Level I classification of the Commission for Environmental Cooperation^67^. If the species range extended into Northern Forests, the species was assigned to that ecoregion; otherwise, the species was assigned to the ecoregion containing the greatest proportion of the species range (either Eastern Temperate Forests or Great Plains/North American Deserts combined). Maximum body sizes in snout-vent length (SVL) were obtained from the AmphiBIO database^68^ or the relevant field guide^69,70^. Upper thermal limits were obtained from the AmphiTherm database, which includes thermal trait estimates for hundreds of species^71^. We corrected taxonomy based on the reported sample coordinates (i.e., *Acris crepitans* localities from the range of *Acris blanchardi*; *Pseudacris triseriata* localities from the range of *Pseudacris maculata*; *Anaxyrus woodhousii* localities from the range of *Anaxyrus fowleri*; *Rana berlandieri* localities from the range of *Rana blairi*; and *Rana pipiens* localities from the range of *Rana sphenocephala*). We used critical thermal maximum (CTmax) values for all species. We prioritized wild-caught, adult frogs from experiments that used the loss of righting response (LRR) as the endpoint where possible. For species with multiple records for upper thermal limit at the same life stage and endpoint, we chose the maximum recorded value.

We fit phylogenetic generalized linear mixed models using *MCMCglmm*^72^, a Bayesian statistics package in R that uses Markov chain Monte Carlo (MCMC) sampling and incorporates phylogenetic information into models. We downloaded 100 phylogeny subsets for the 46 species in our dataset from VertLife.org, which is based on the relationships from Jetz and Pyron^73^ for the amphibian tree of life. Given the consistency of relationships among replicates, we used the first random tree for our analyses. We standardized continuous predictors to a mean of zero and a standard deviation of one. We built models with the full dataset of 44 species and only five predictors and with a reduced dataset including only the 25 species that had thermal tolerance data to include all six predictors. We set priors of *V* = 1, nu = 0.02 for the phylogenetic random effect, used a categorical family for binary responses, and used a gaussian family for continuous responses. We ran each model across four chains of 5,000,000 iterations with a burn-in period of 1,000,000 and thinning every 4,000 steps. We visually inspected trace plots, checked Gelman diagnostic statistics using the “gelman.diag()” function^74^, and checked effective sample sizes (ESS) values to evaluate convergence. We combined results from the four chains to plot the posterior means and 95% credible intervals for each predictor.

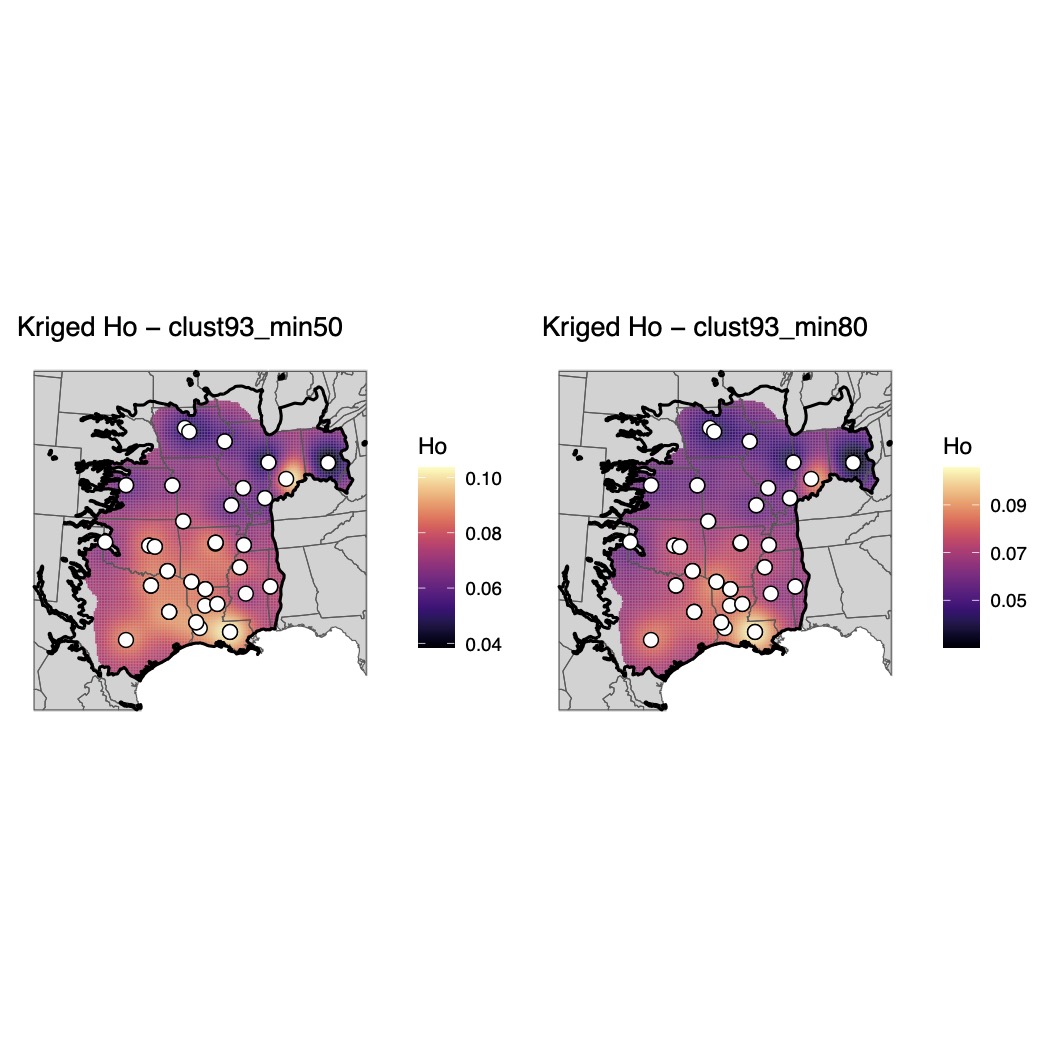

**Supplementary Fig. 1.**

Maps of genomic diversity for *Acris blanchardi* generated with the *WINGEN* R package.

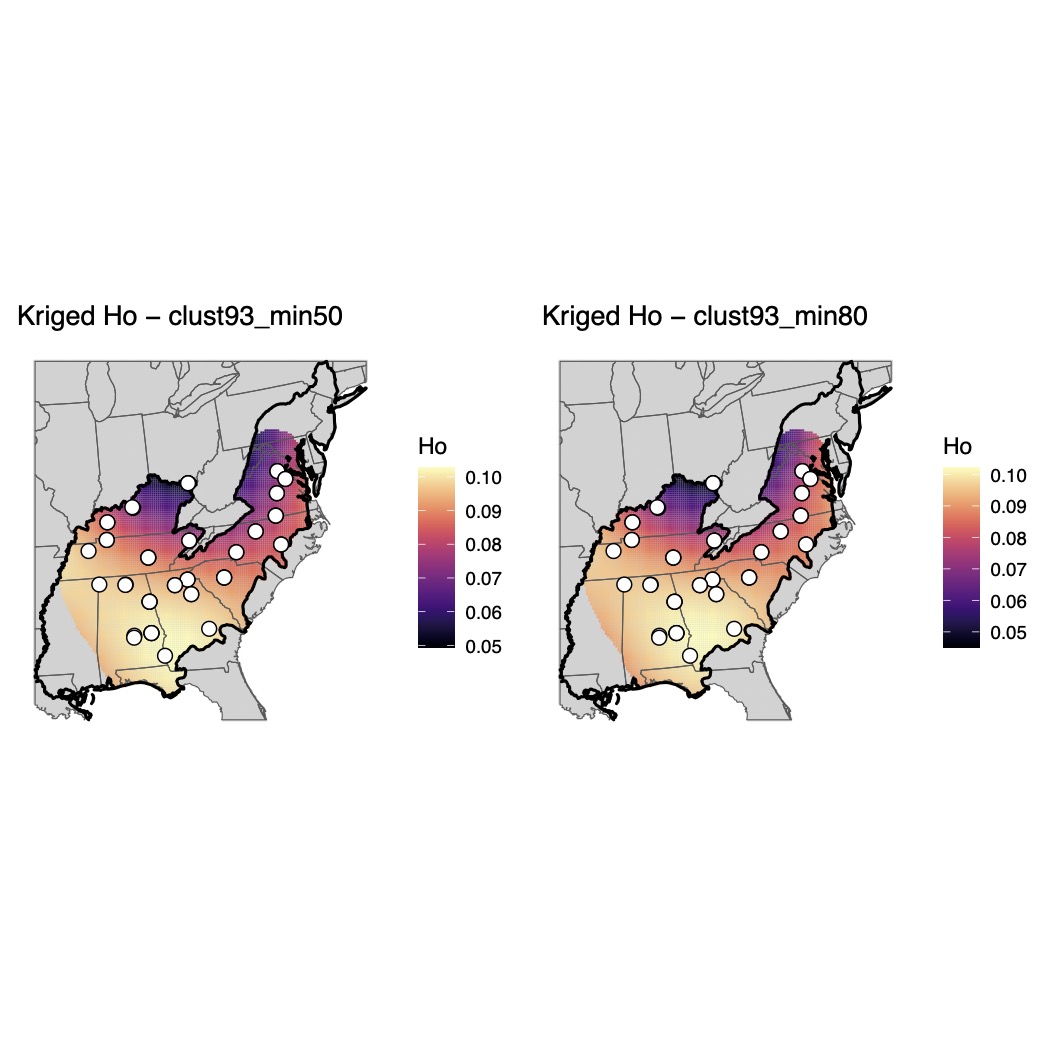

Supplementary Fig. 2.

Maps of genomic diversity for *Acris crepitans* generated with the *WINGEN* R package.

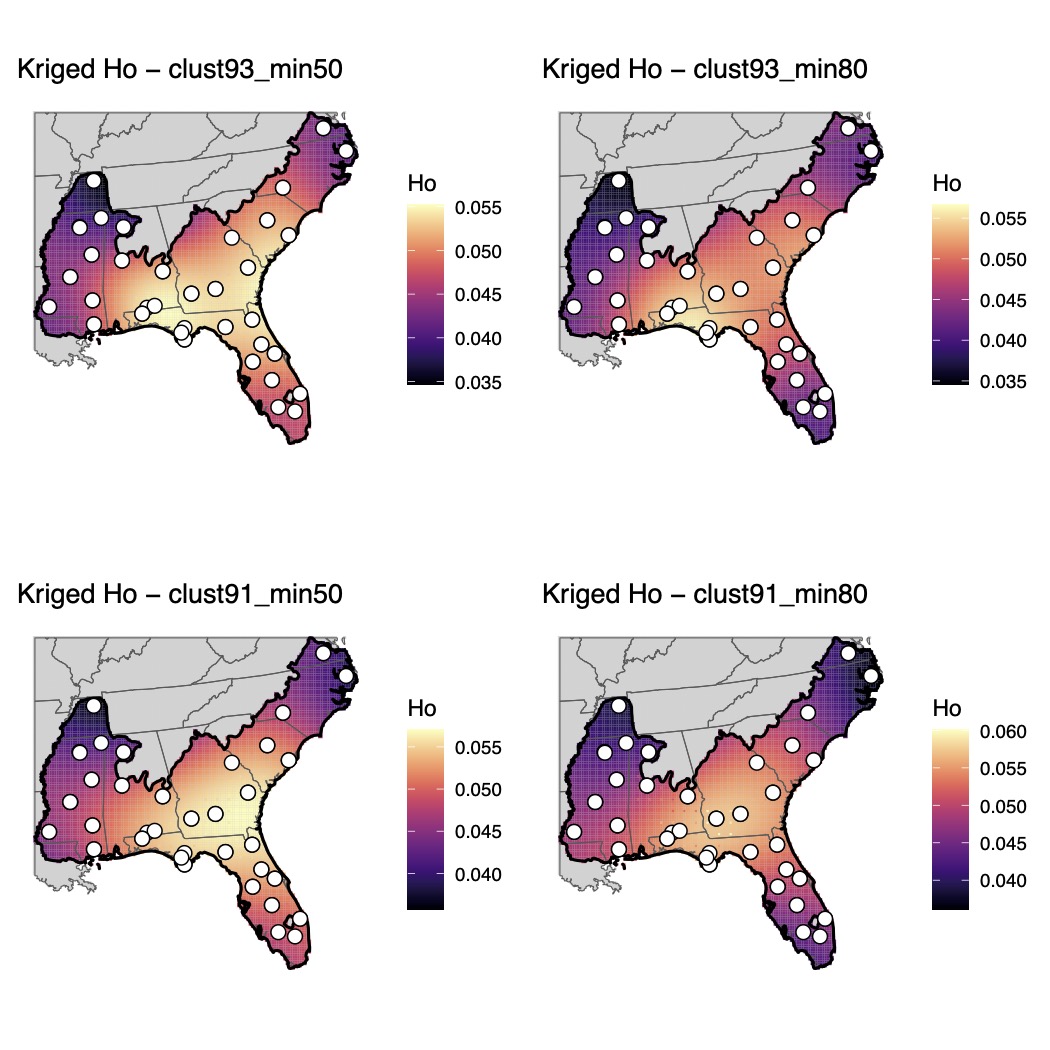

Supplementary Fig. 3.

Maps of genomic diversity for *Acris gryllus* generated with the *WINGEN* R package.

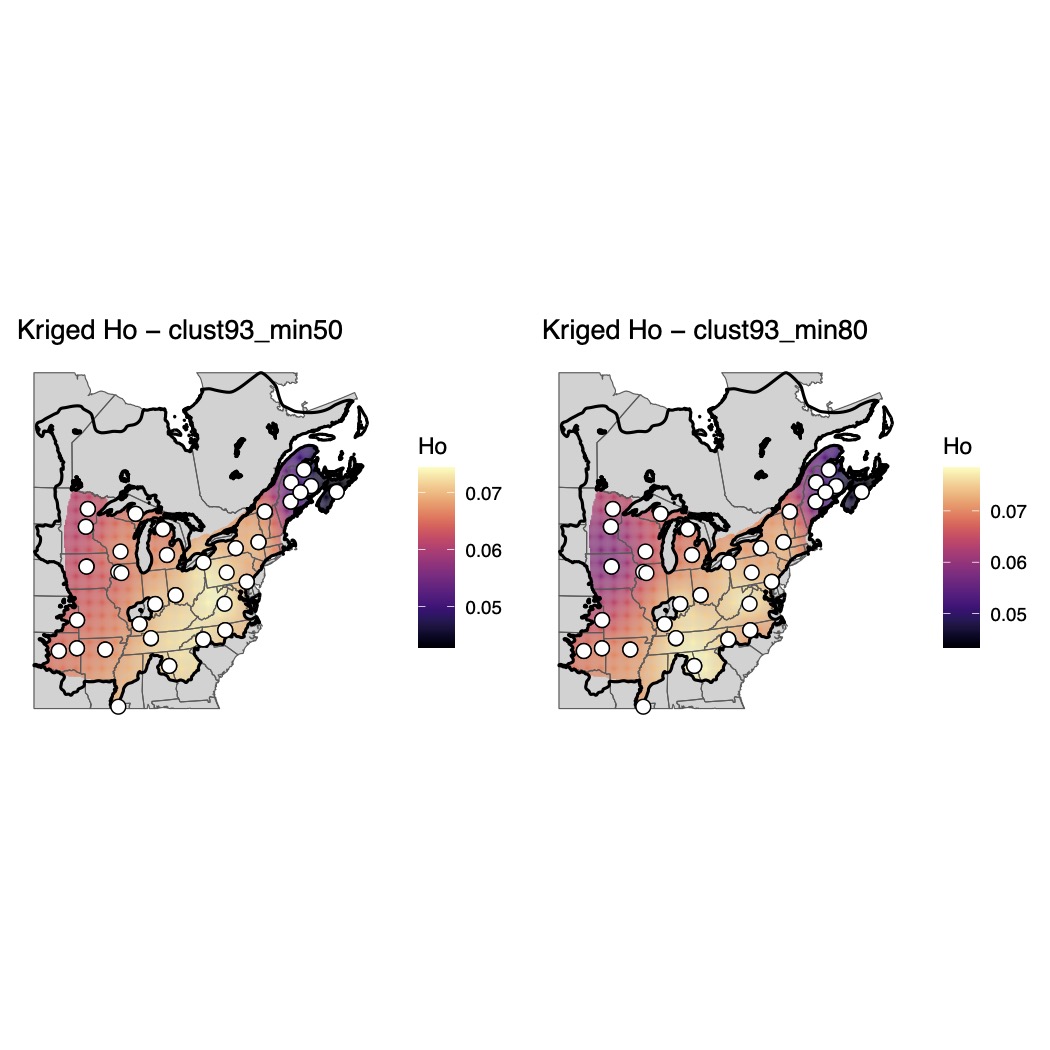

Supplementary Fig. 4.

Maps of genomic diversity for *Anaxyrus americanus* generated with the *WINGEN* R package.

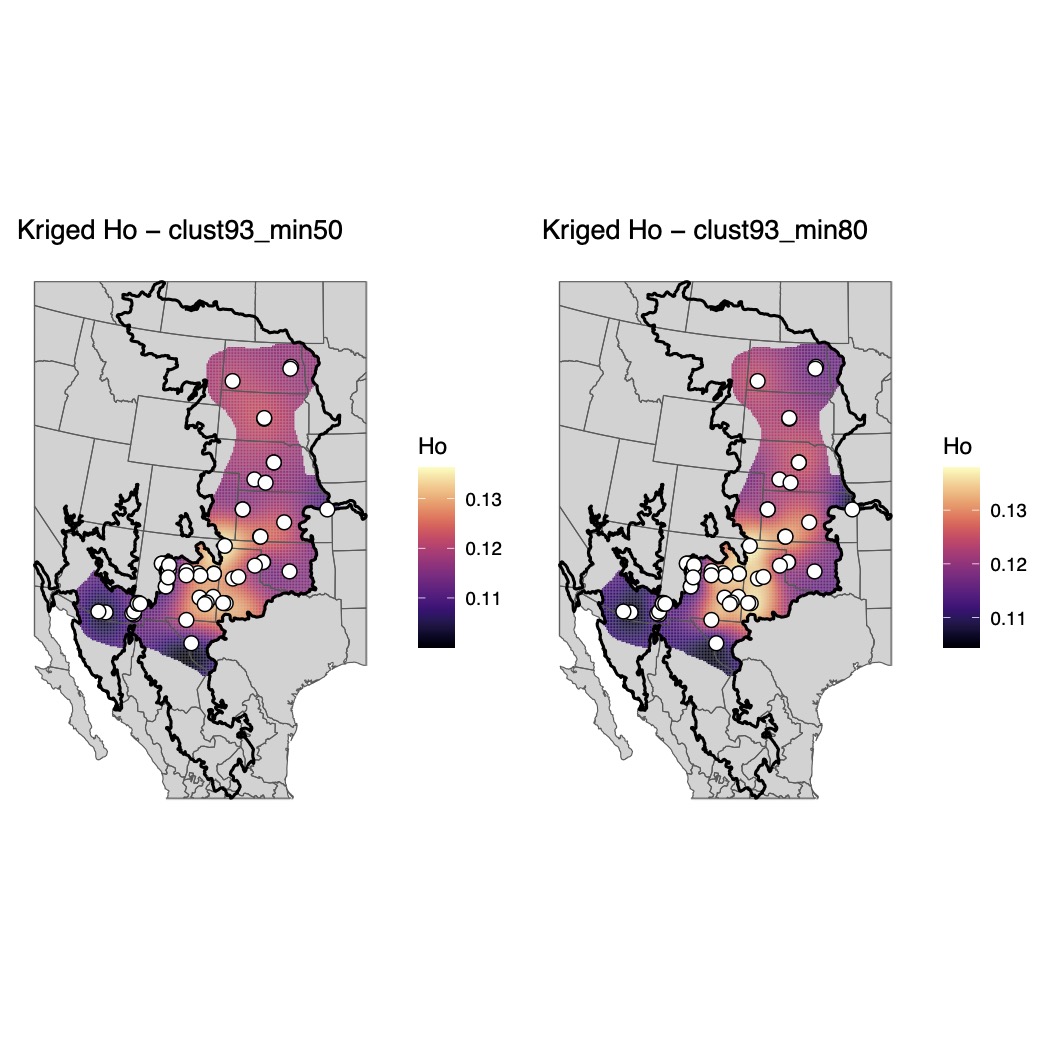

Supplementary Fig. 5.

Maps of genomic diversity for *Anaxyrus cognatus* generated with the *WINGEN* R package.

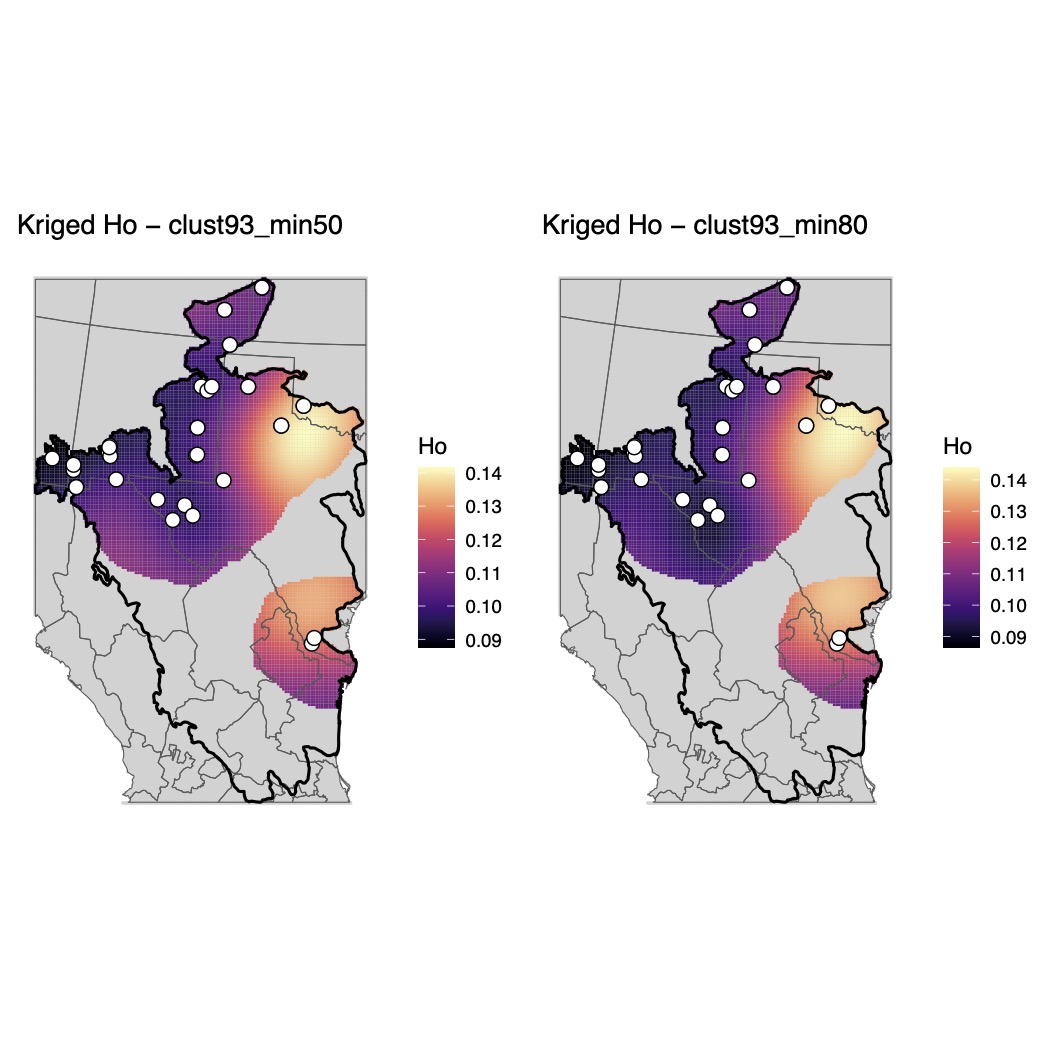

Supplementary Fig. 6.

Maps of genomic diversity for *Anaxyrus debilis* generated with the *WINGEN* R package.

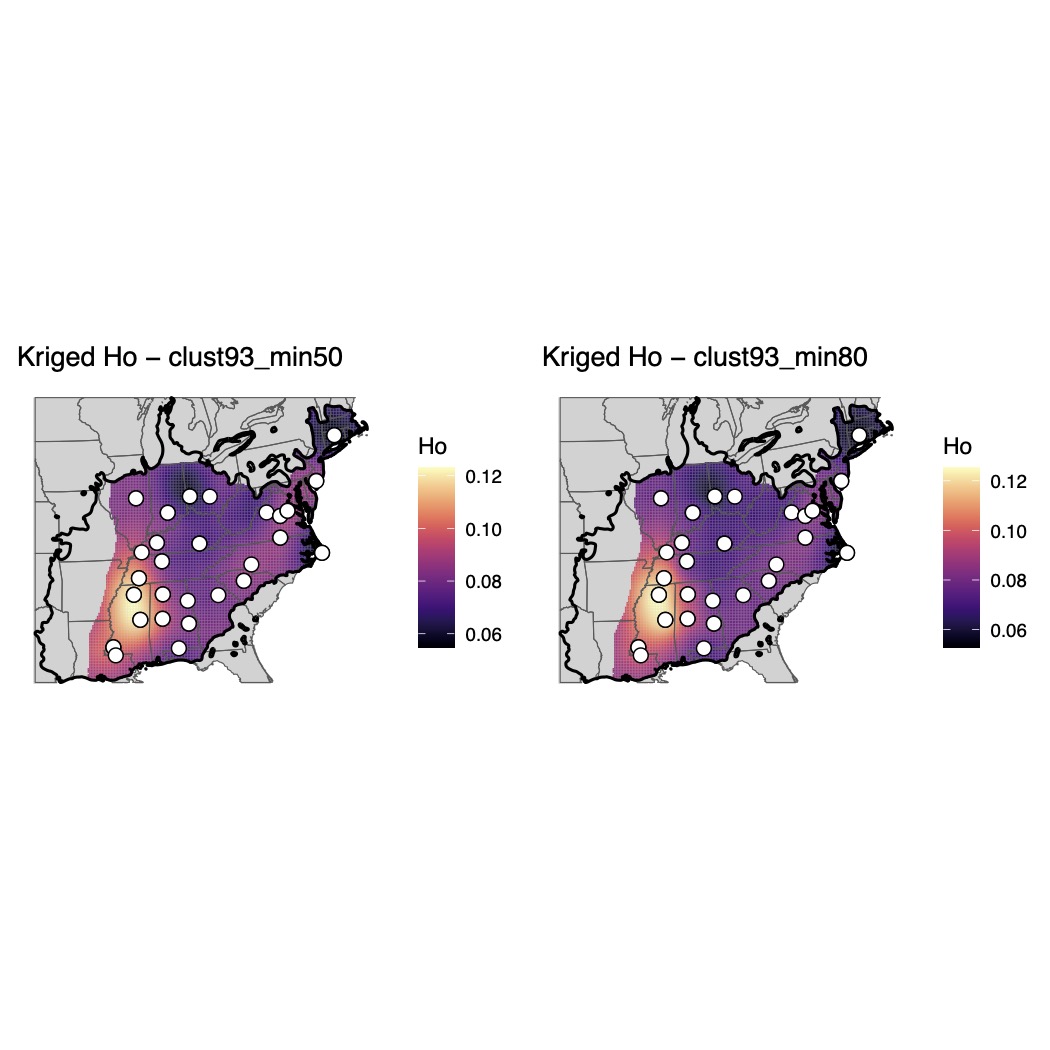

Supplementary Fig. 7.

Maps of genomic diversity for *Anaxyrus fowleri* generated with the *WINGEN* R package.

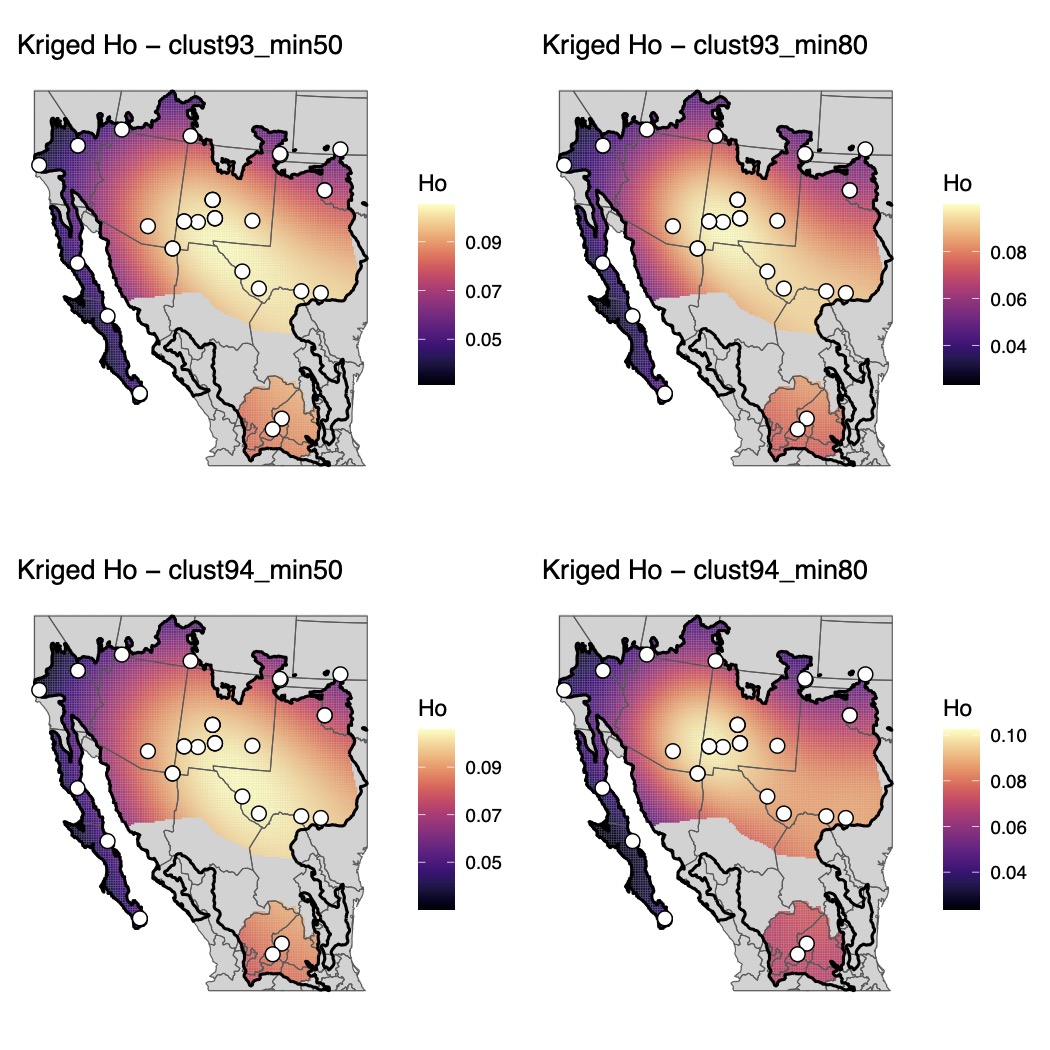

Supplementary Fig. 8.

Maps of genomic diversity for *Anaxyrus punctatus* generated with the *WINGEN* R package.

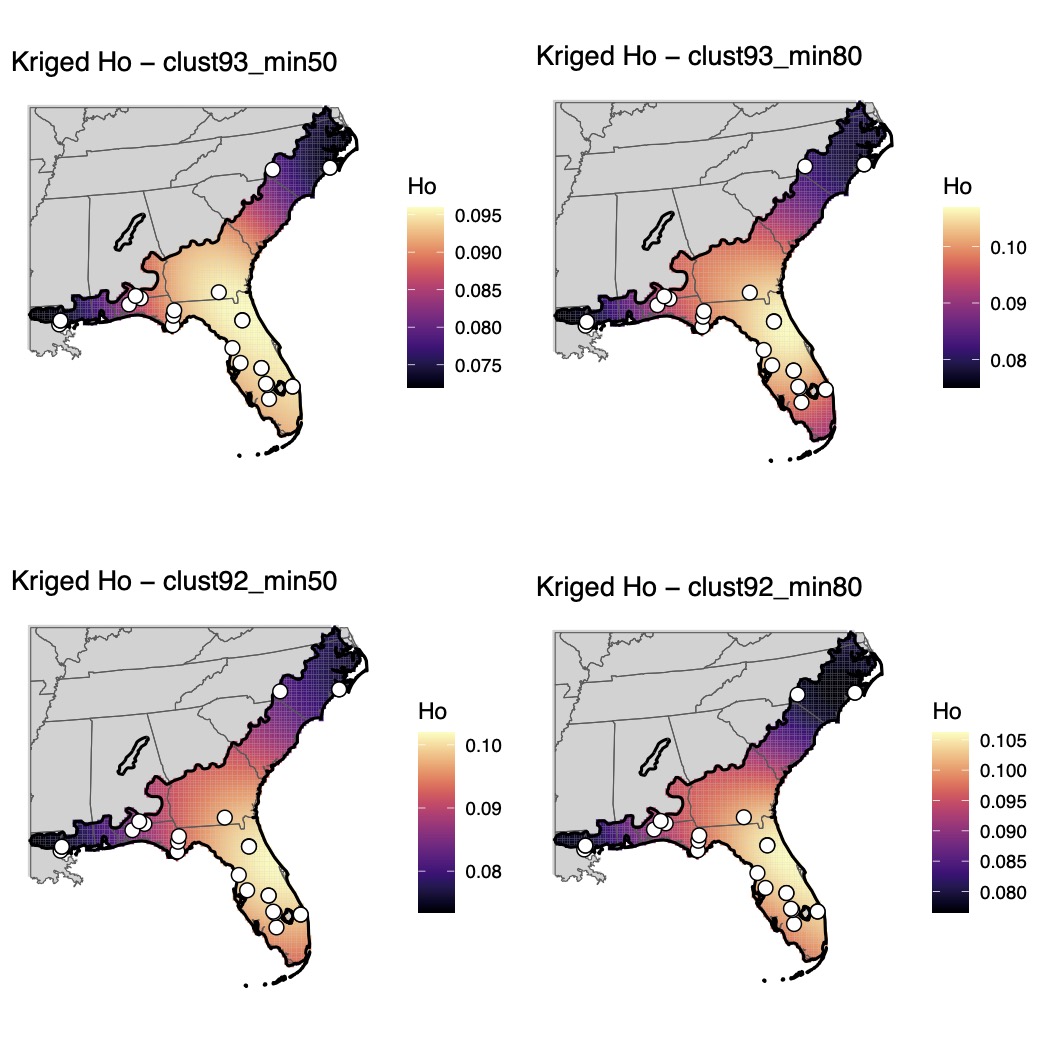

Supplementary Fig. 9.

Maps of genomic diversity for *Anaxyrus quercicus* generated with the *WINGEN* R package.

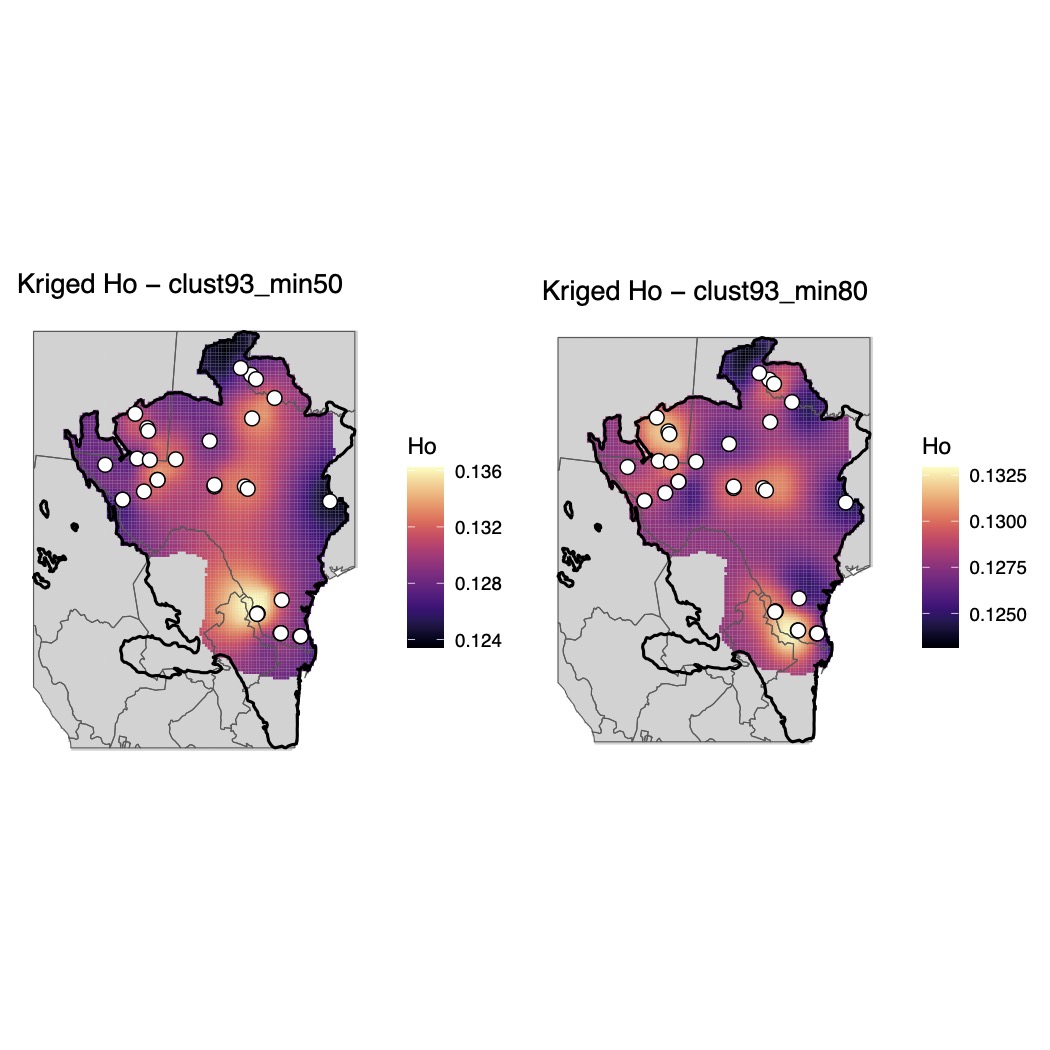

Supplementary Fig. 10.

Maps of genomic diversity for *Anaxyrus speciosus* generated with the *WINGEN* R package.

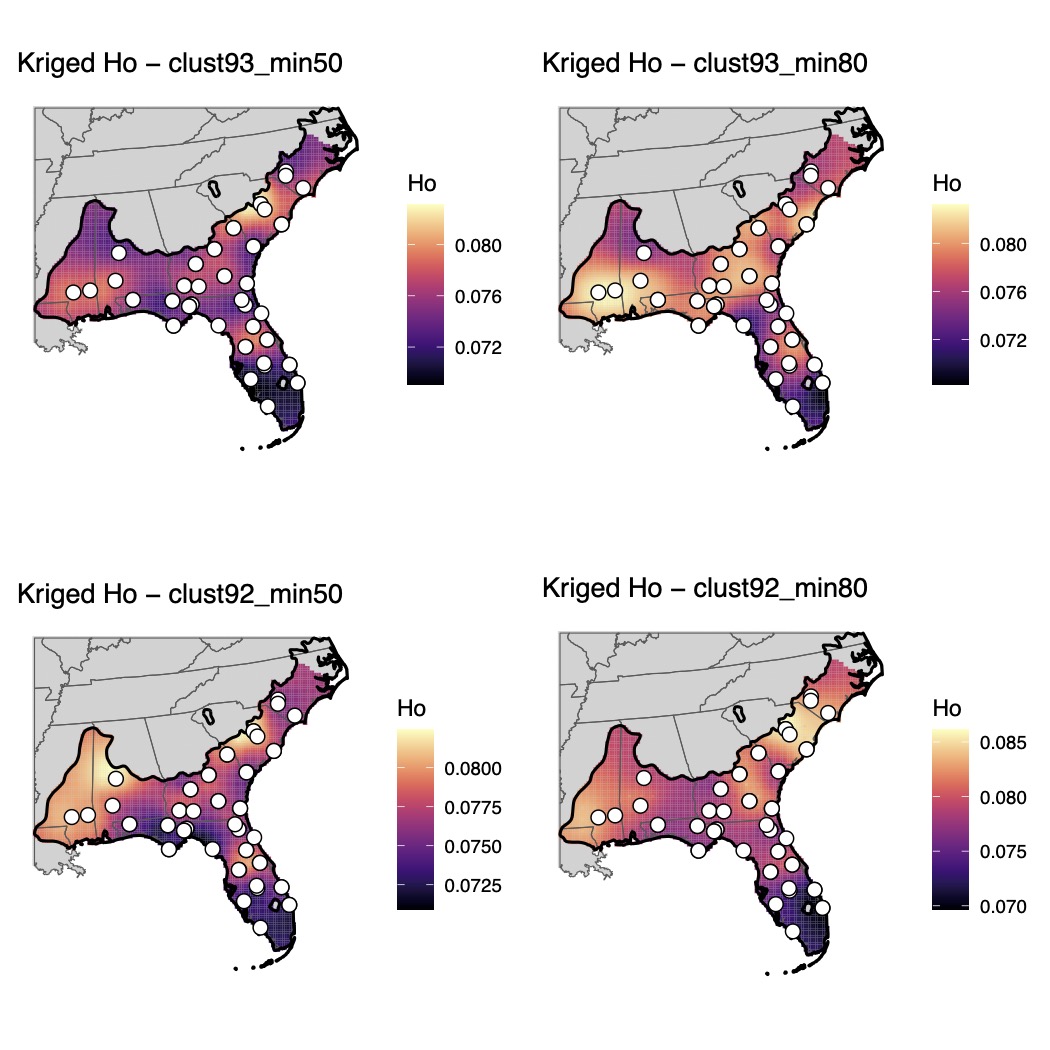

Supplementary Fig. 11.

Maps of genomic diversity for *Anaxyrus terrestris* generated with the *WINGEN* R package.

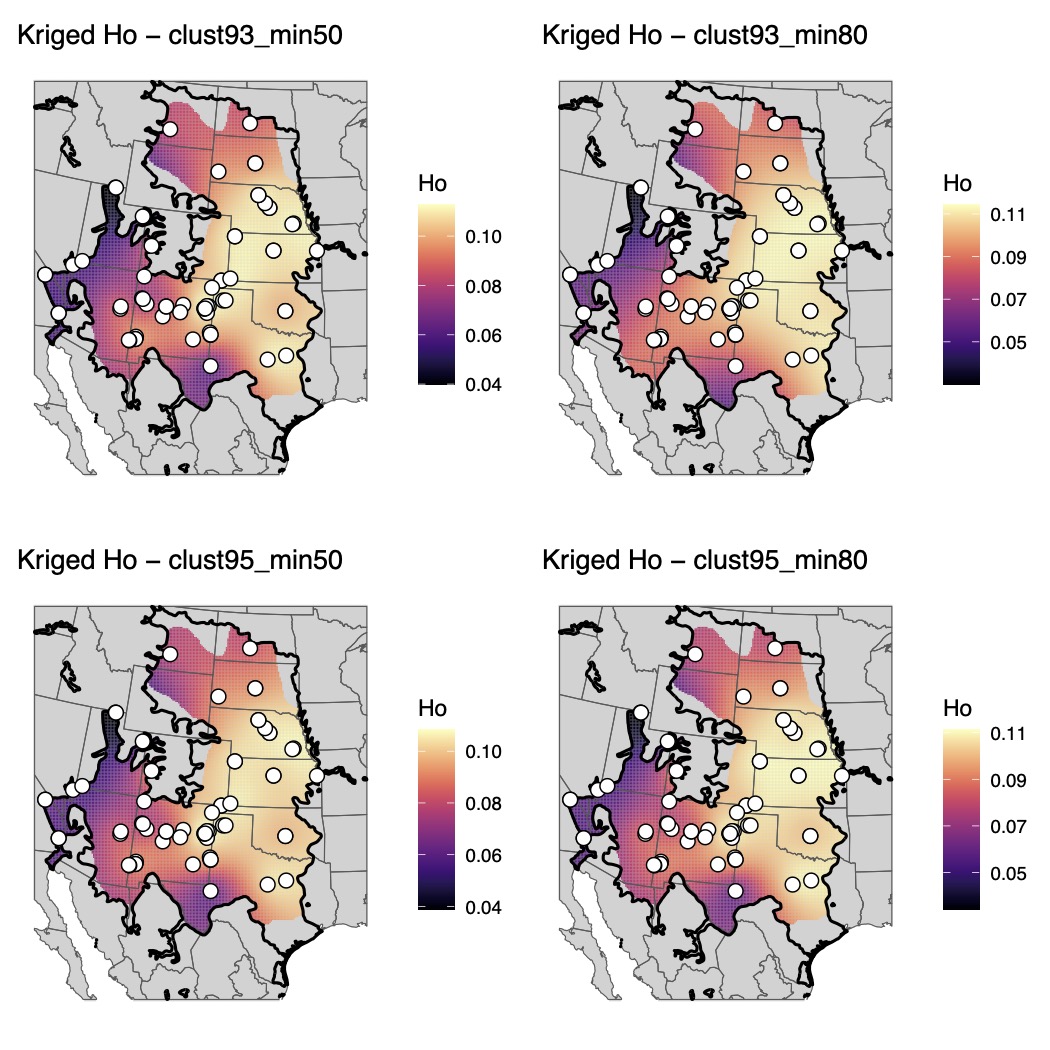

Supplementary Fig. 12.

Maps of genomic diversity for *Anaxyrus woodhousii* generated with the *WINGEN* R package.

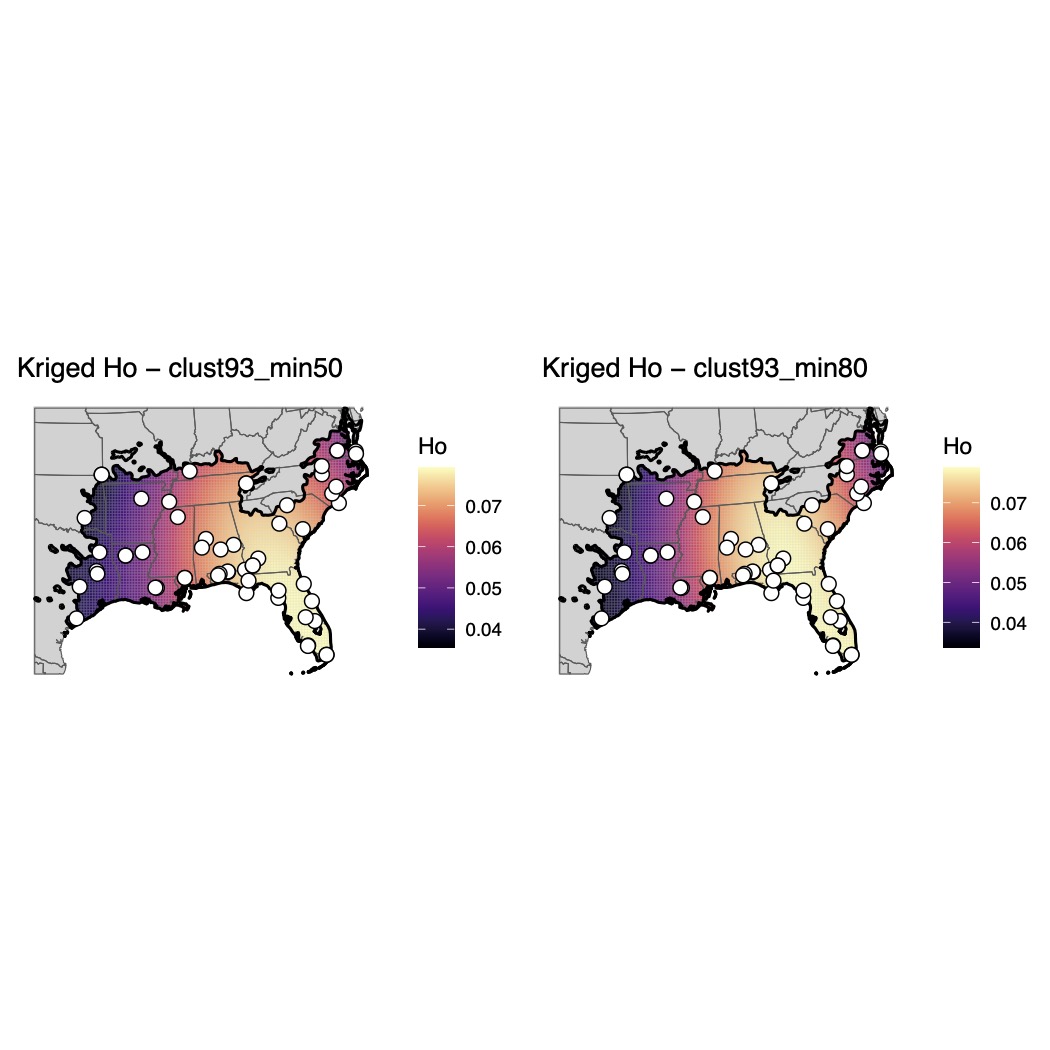

Supplementary Fig. 13.

Maps of genomic diversity for *Gastrophryne carolinensis* generated with the *WINGEN* R package.

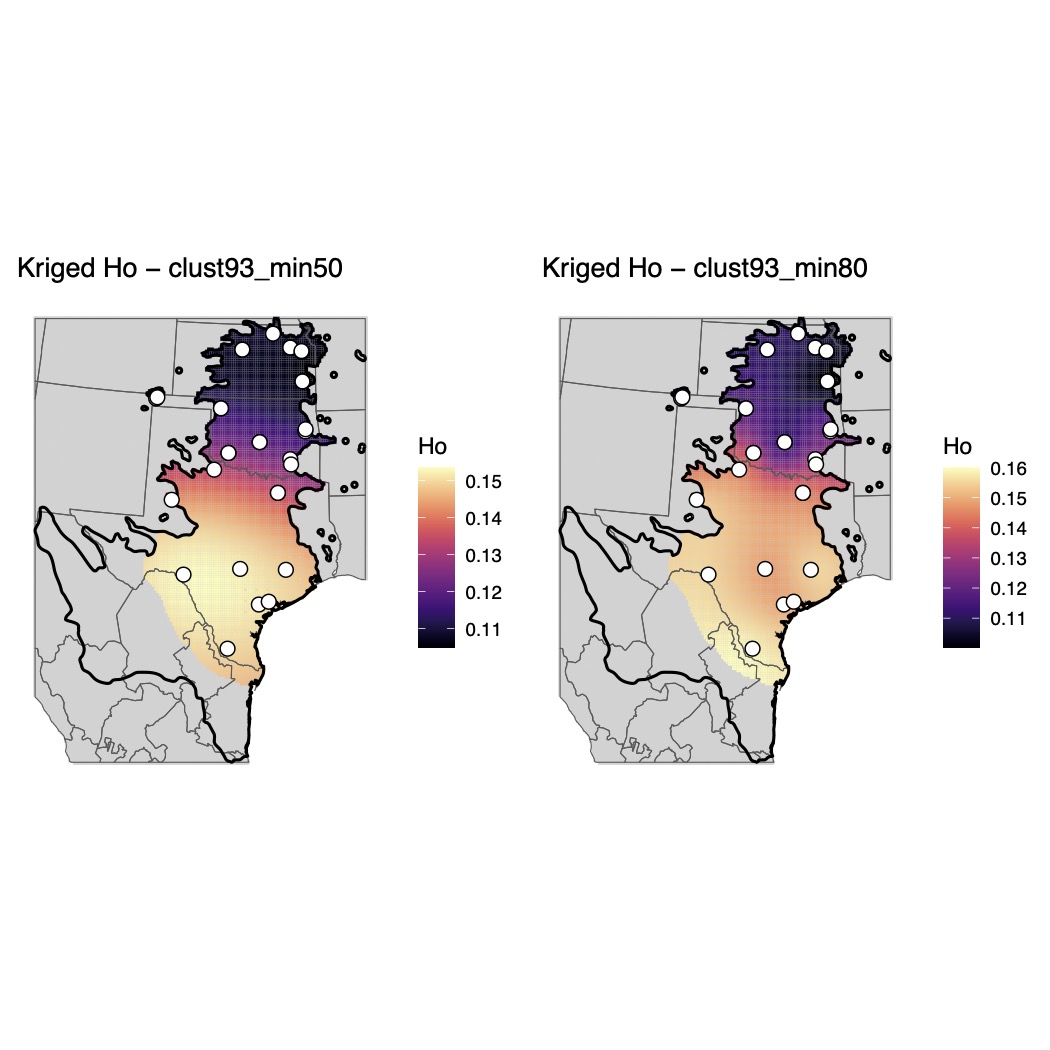

Supplementary Fig. 14.

Maps of genomic diversity for *Gastrophryne olivacea* generated with the *WINGEN* R package.

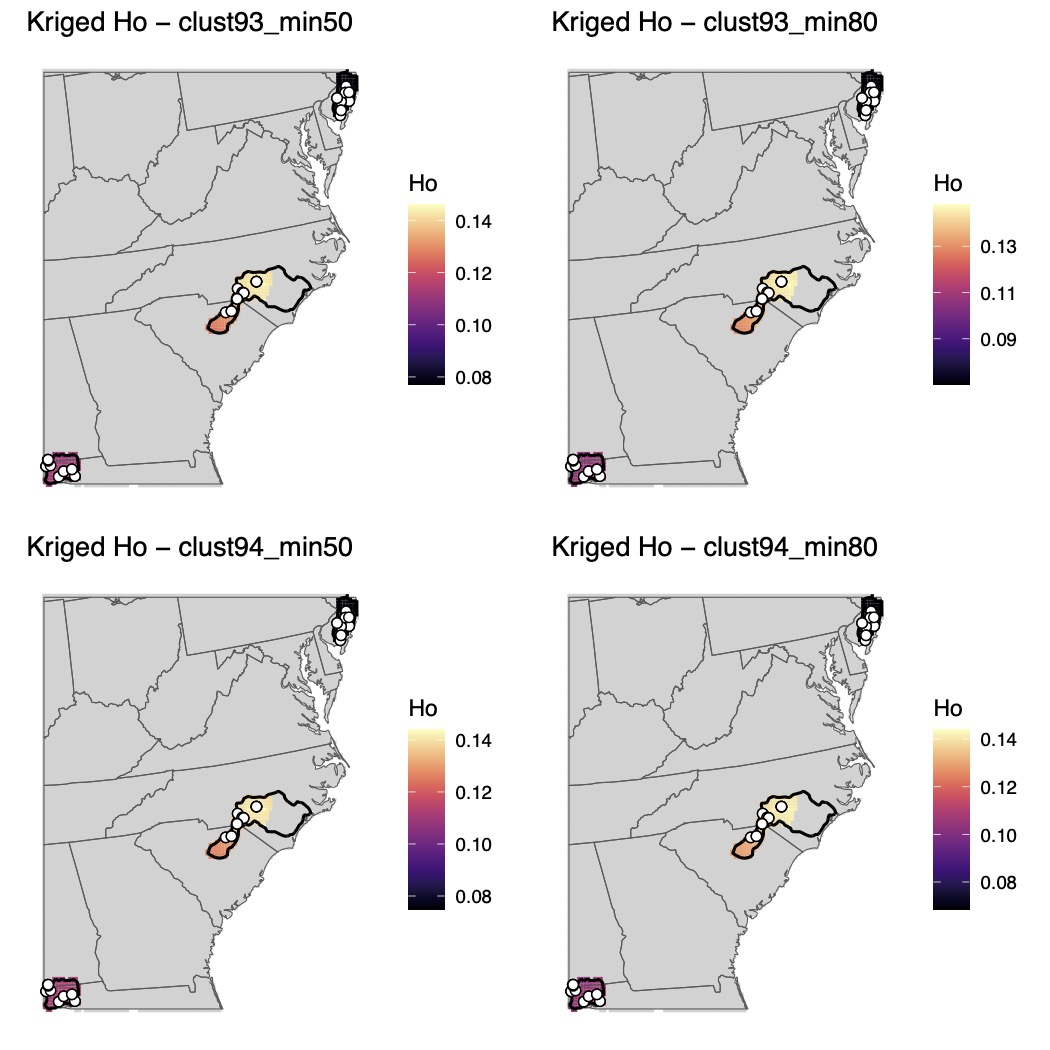

Supplementary Fig. 15.

Maps of genomic diversity for *Hyla andersonii* generated with the *WINGEN* R package.

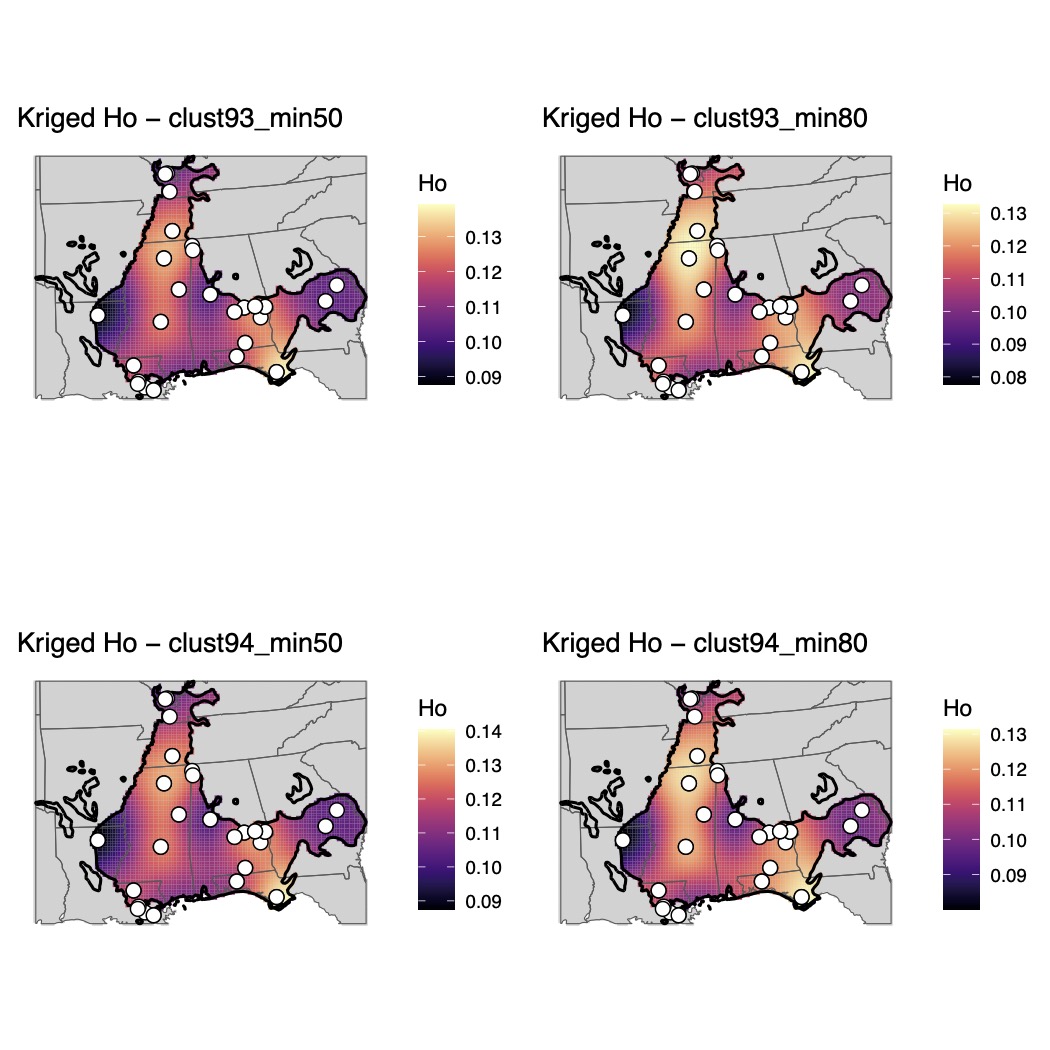

Supplementary Fig. 16.

Maps of genomic diversity for *Hyla avivoca* generated with the *WINGEN* R package.

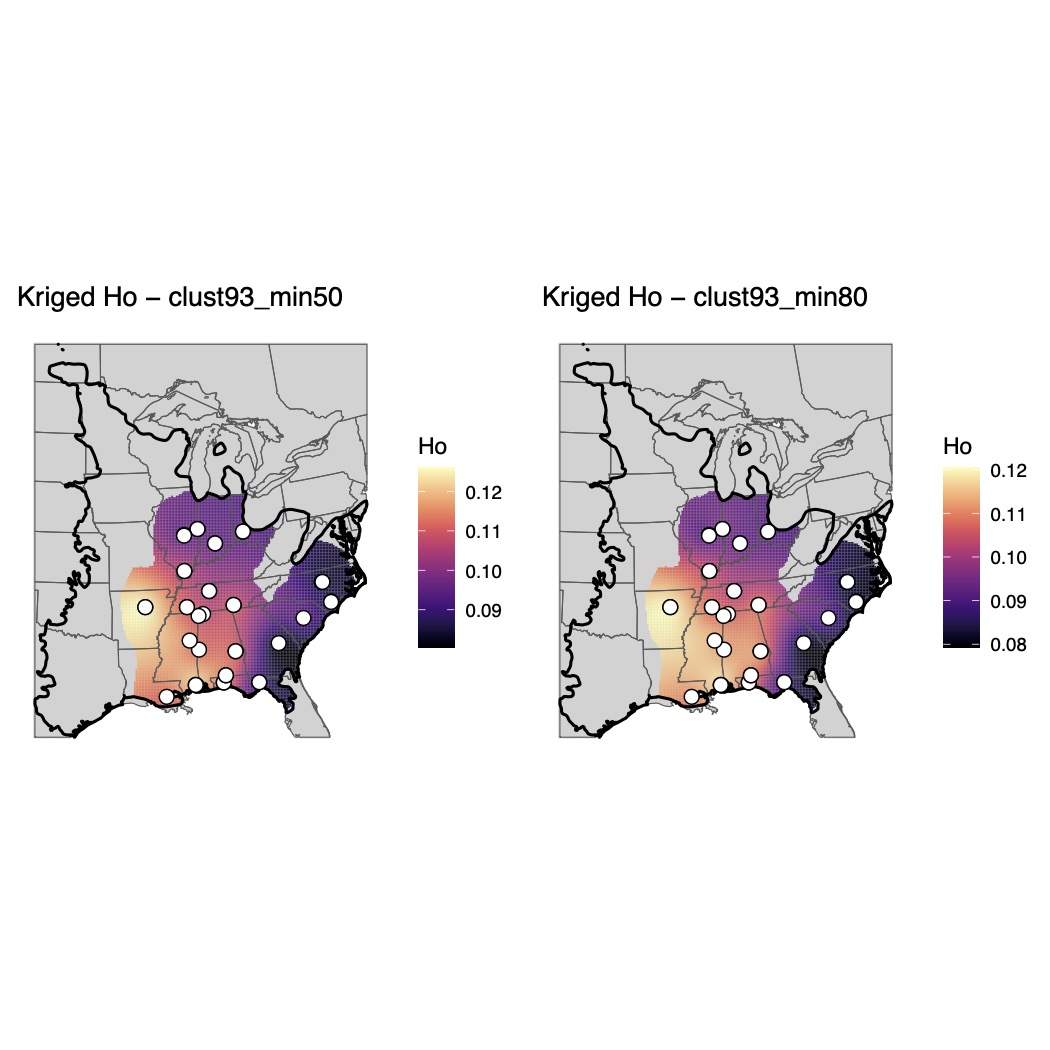

Supplementary Fig. 17.

Maps of genomic diversity for *Hyla chrysoscelis* generated with the *WINGEN* R package.

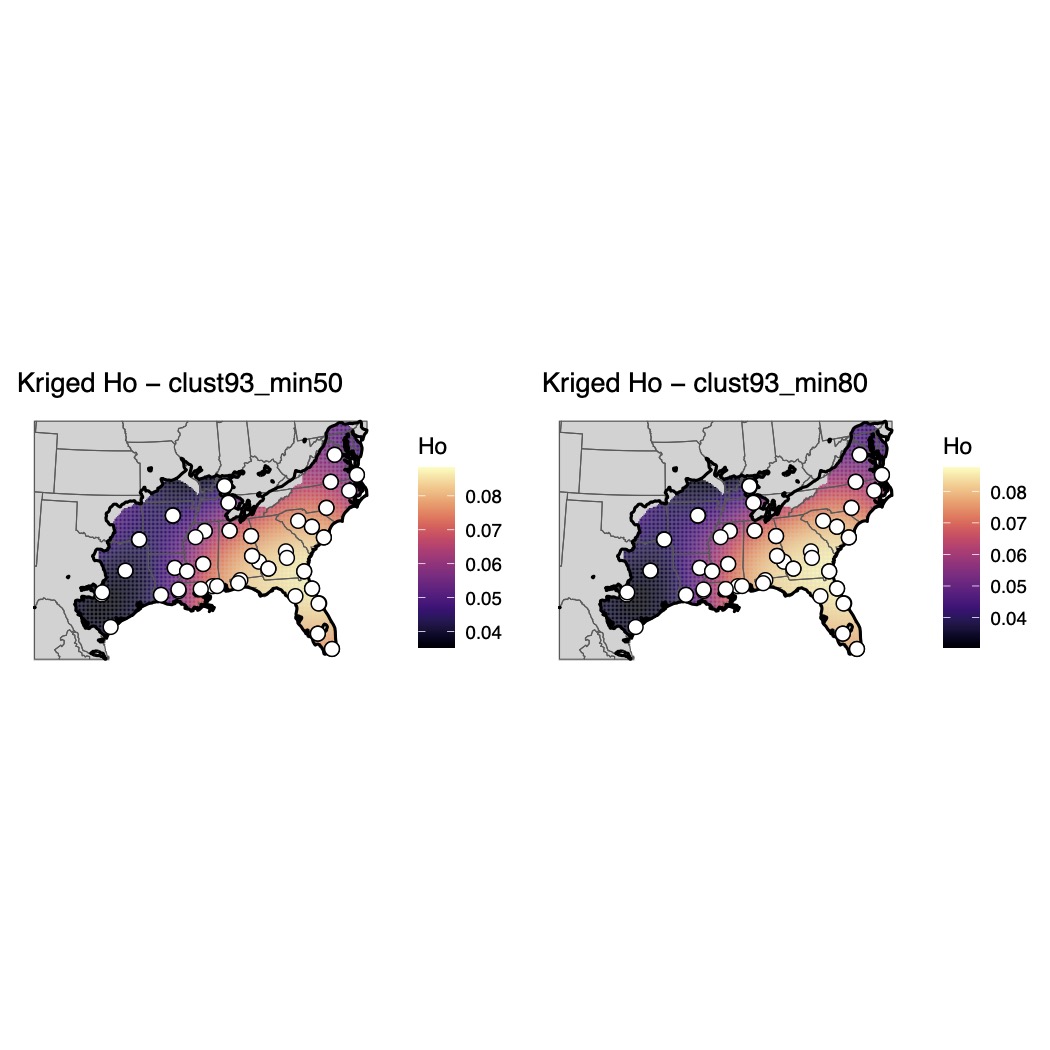

Supplementary Fig. 18.

Maps of genomic diversity for *Hyla cinerea* generated with the *WINGEN* R package.

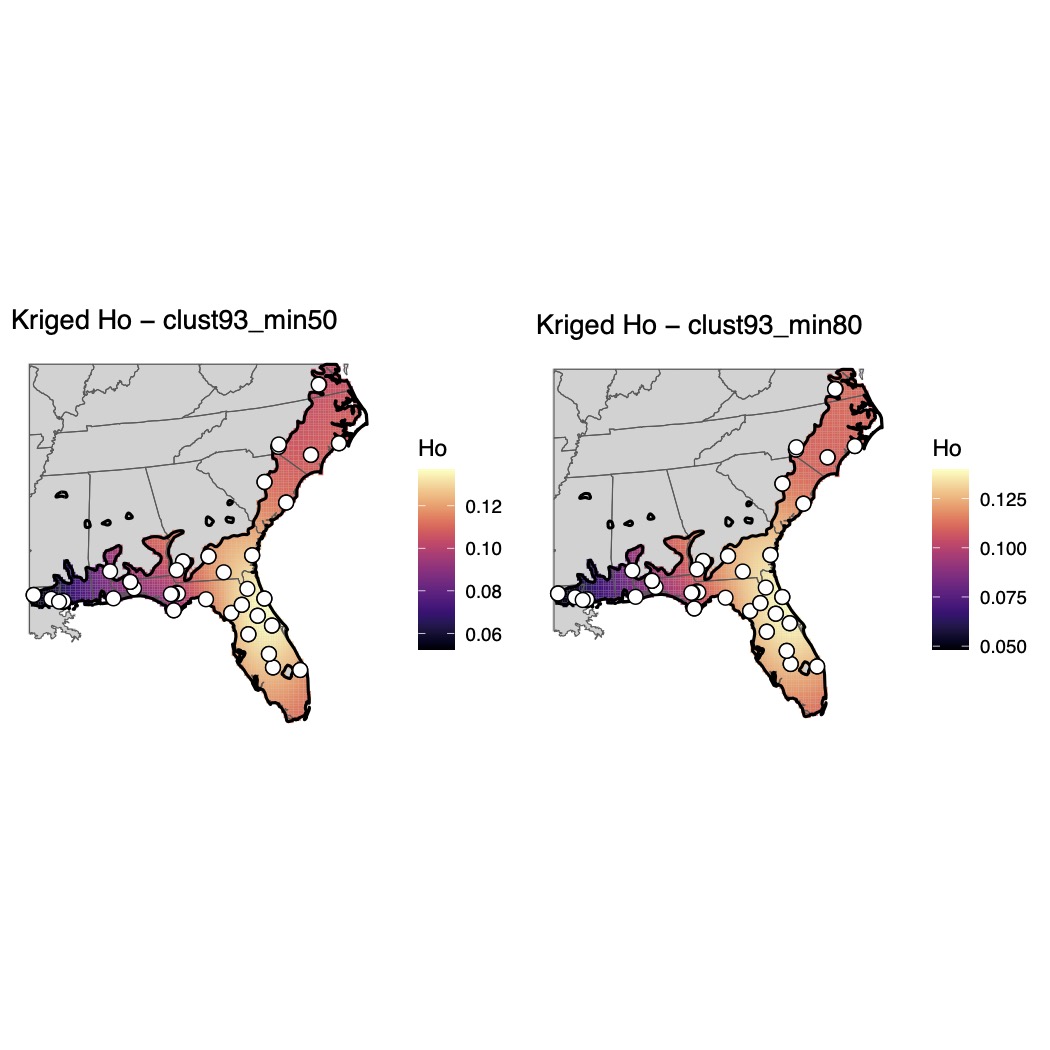

Supplementary Fig. 19.

Maps of genomic diversity for *Hyla femoralis* generated with the *WINGEN* R package.

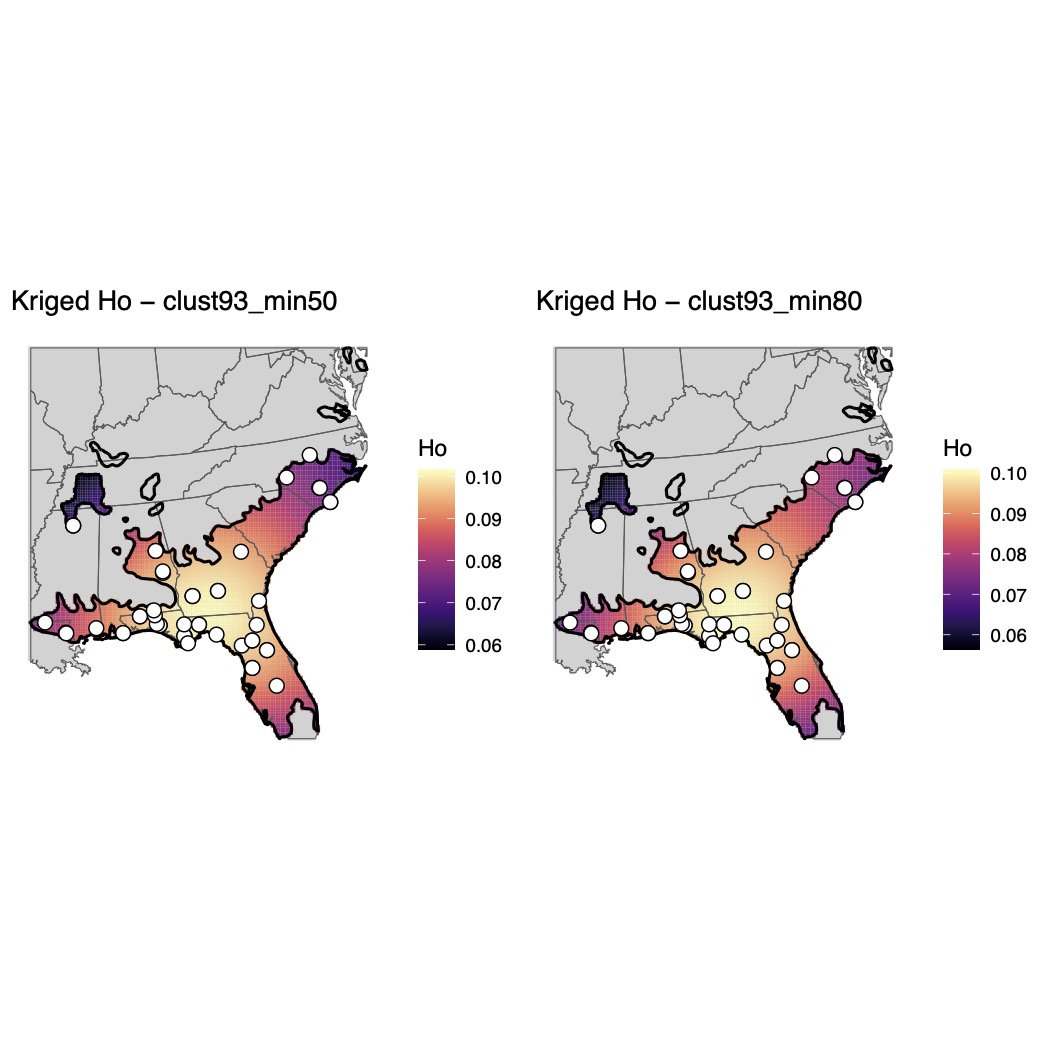

Supplementary Fig. 20.

Maps of genomic diversity for *Hyla gratiosa* generated with the *WINGEN* R package.

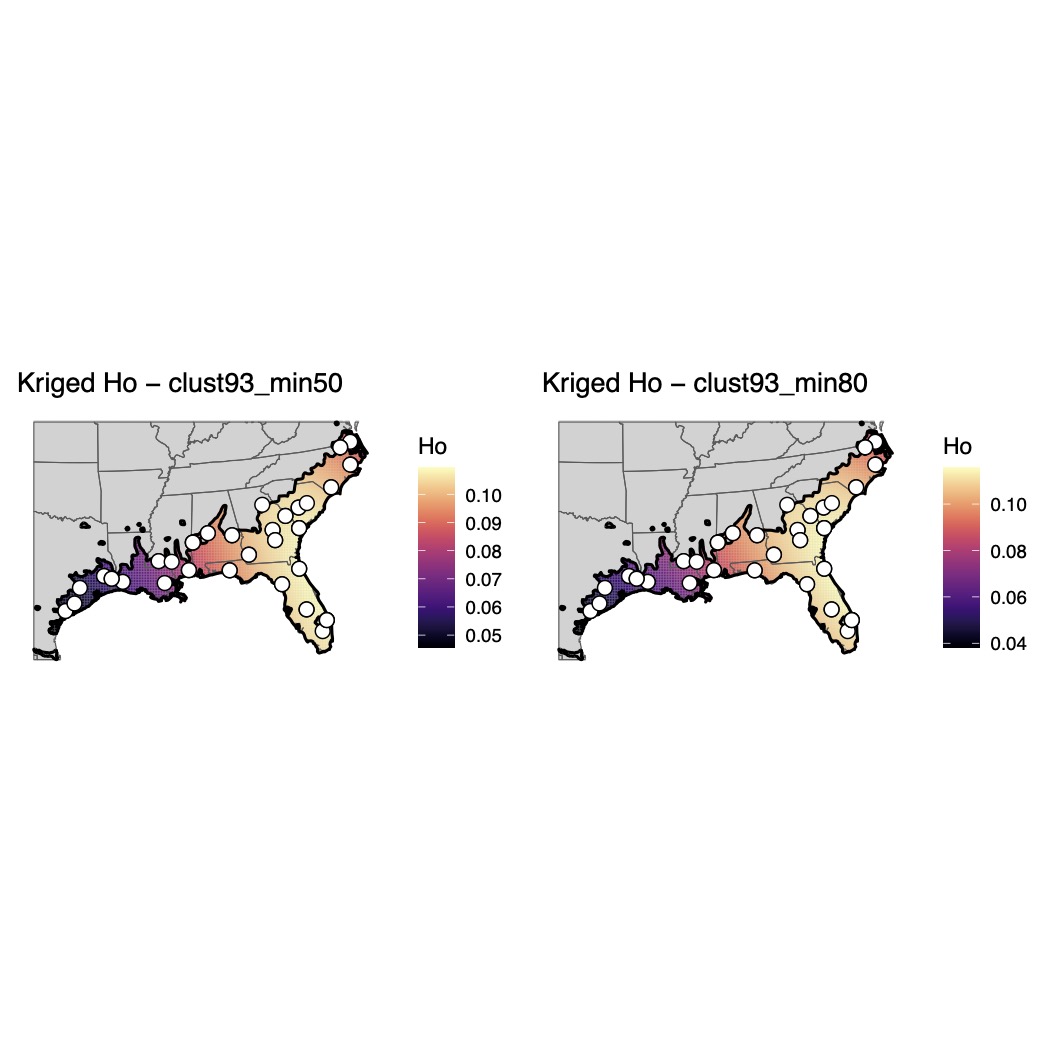

Supplementary Fig. 21.

Maps of genomic diversity for *Hyla squirella* generated with the *WINGEN* R package.

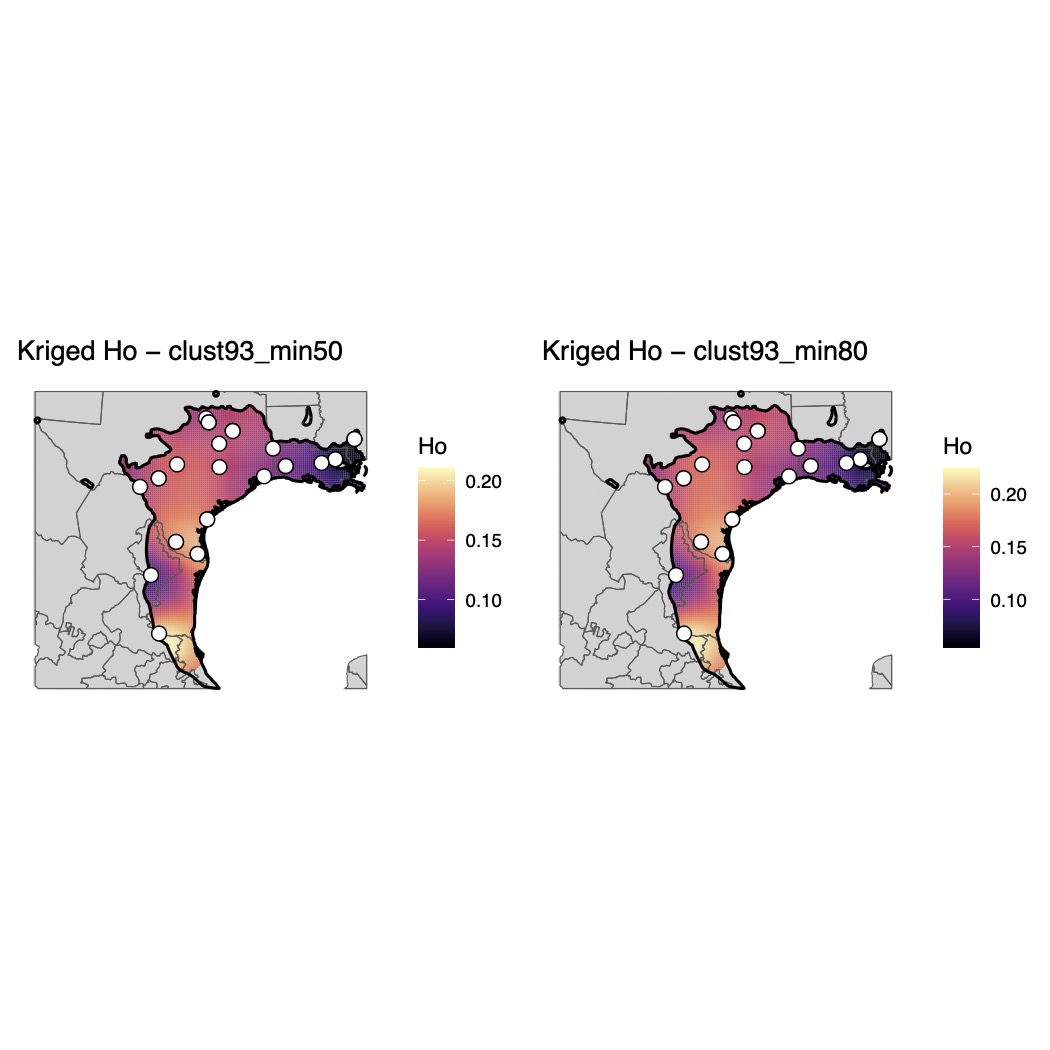

Supplementary Fig. 22.

Maps of genomic diversity for *Incilius nebulifer* generated with the *WINGEN* R package.

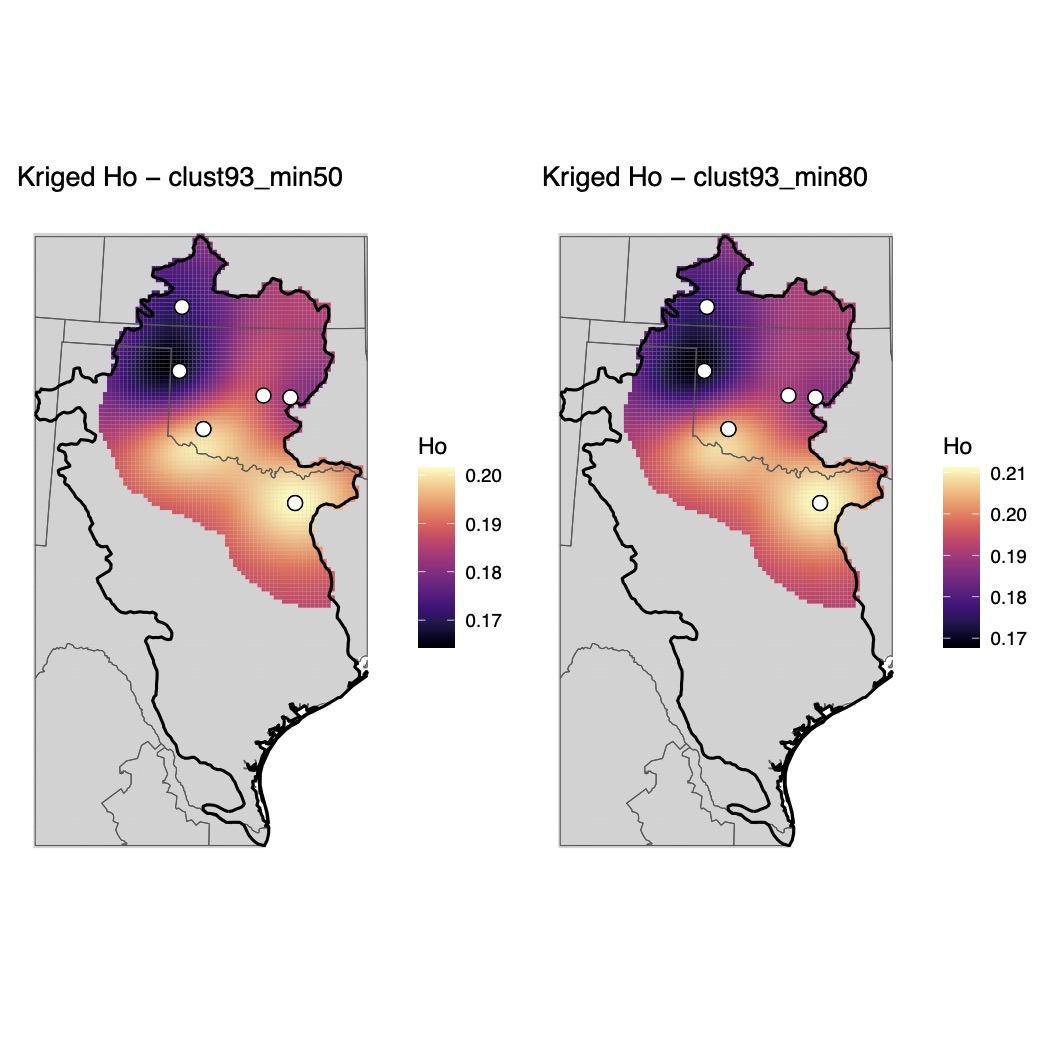

Supplementary Fig. 23.

Maps of genomic diversity for *Pseudacris clarkii* generated with the *WINGEN* R package.

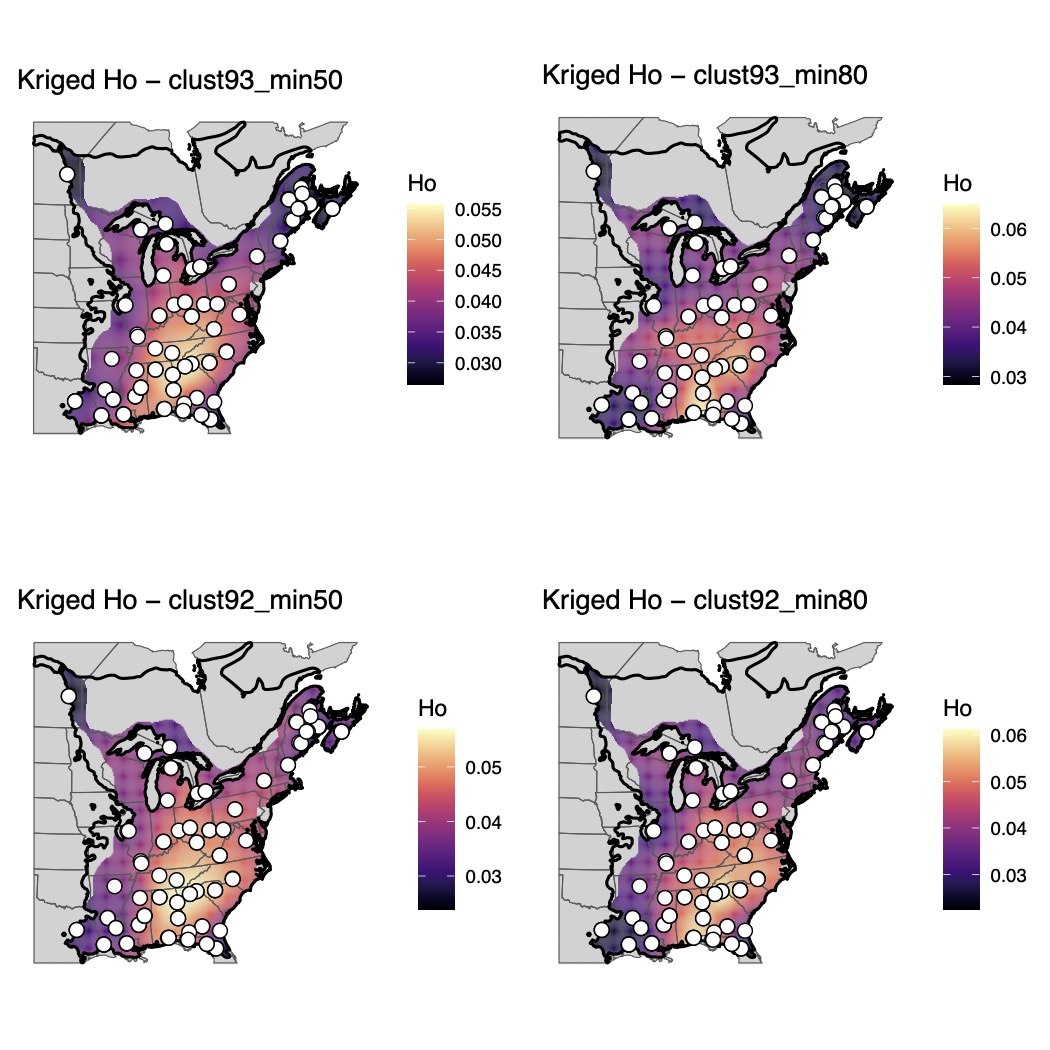

Supplementary Fig. 24.

Maps of genomic diversity for *Pseudacris crucifer* generated with the *WINGEN* R package.

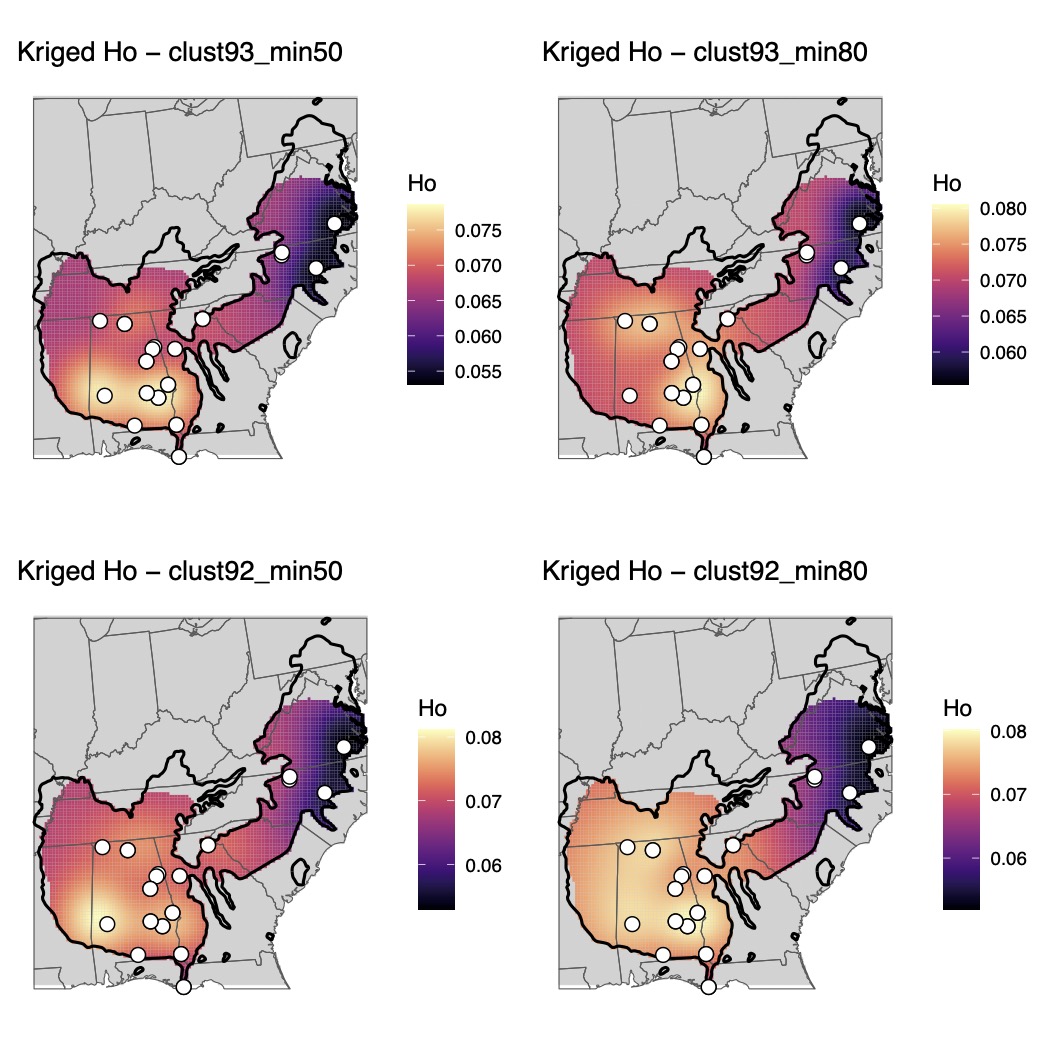

Supplementary Fig. 25.

Maps of genomic diversity for *Pseudacris feriarum* generated with the *WINGEN* R package.

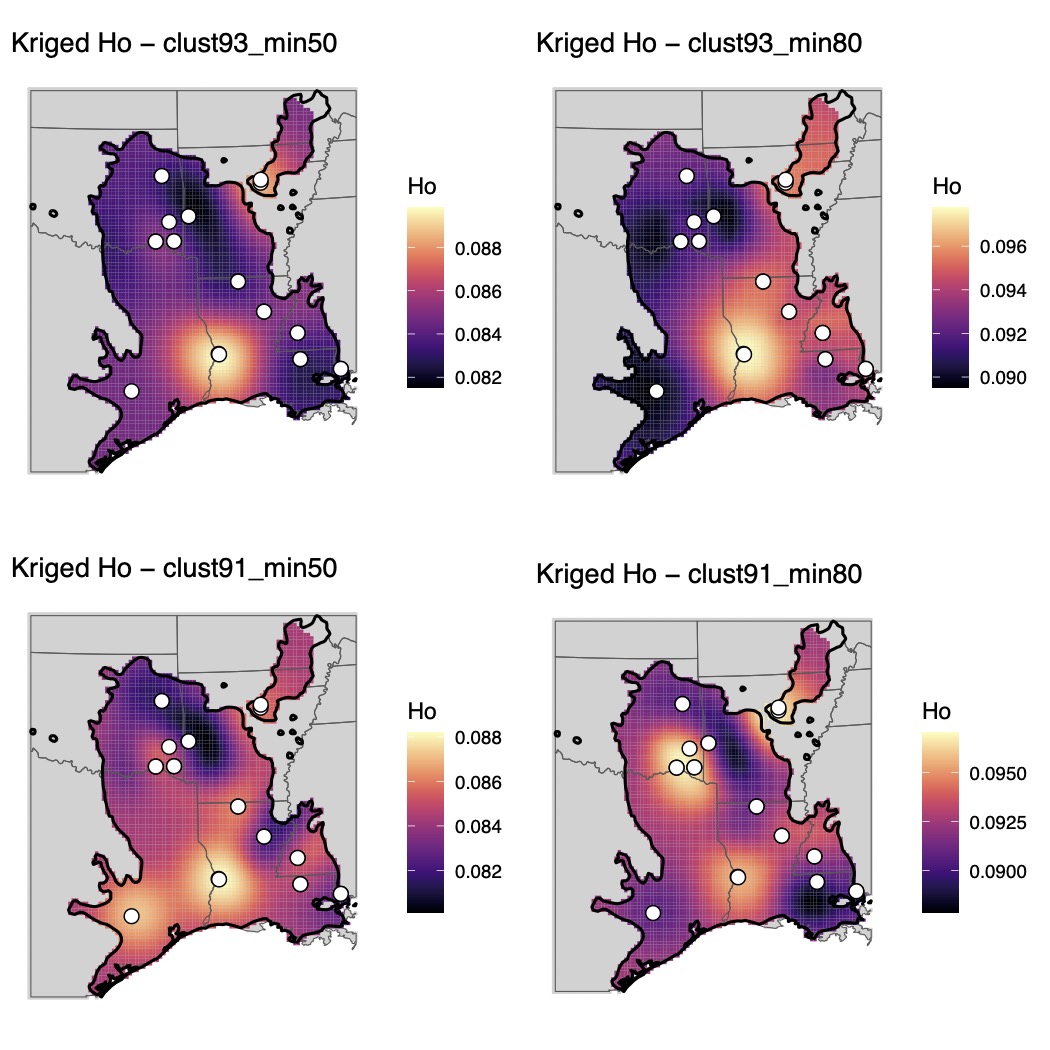

Supplementary Fig. 26.

Maps of genomic diversity for *Pseudacris fouquettei* generated with the *WINGEN* R package.

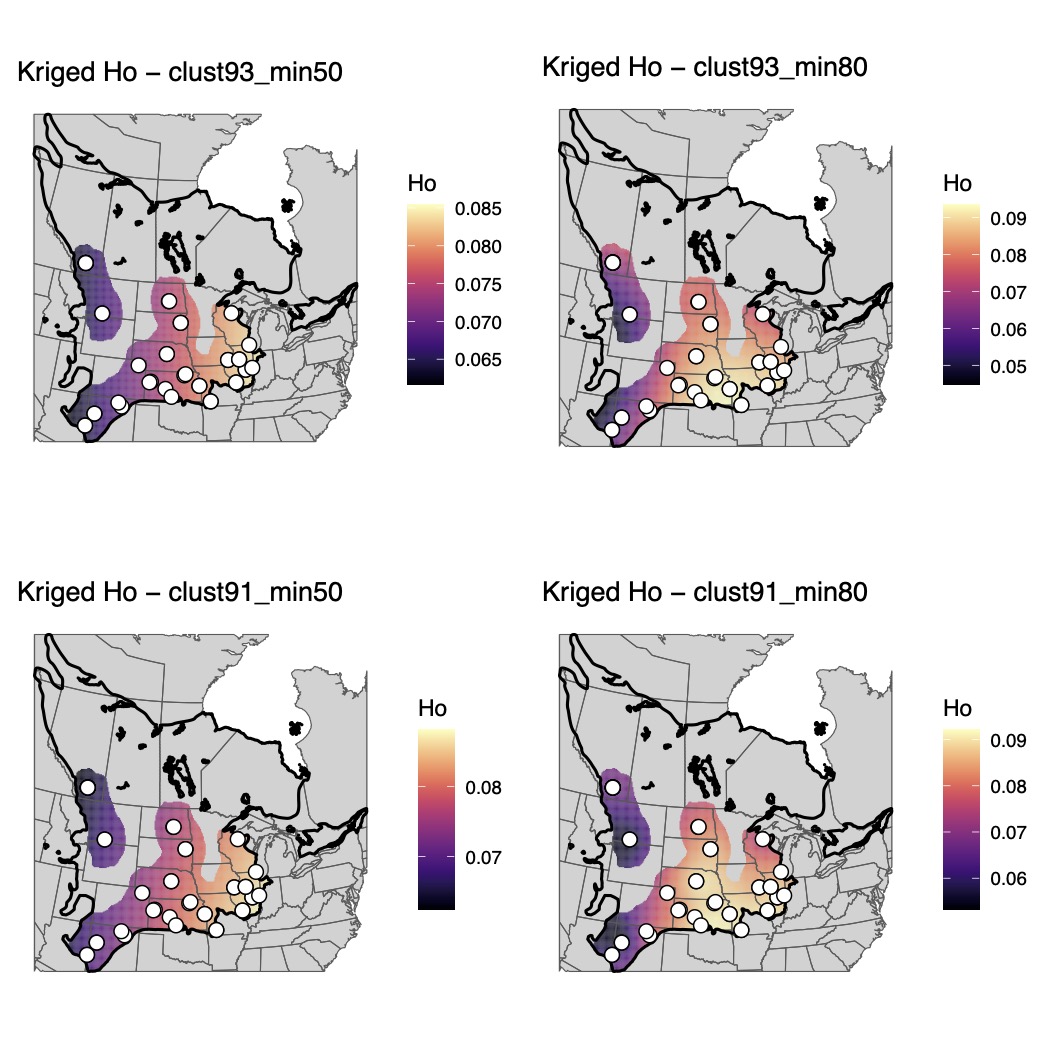

Supplementary Fig. 27.

Maps of genomic diversity for *Pseudacris maculata* generated with the *WINGEN* R package.

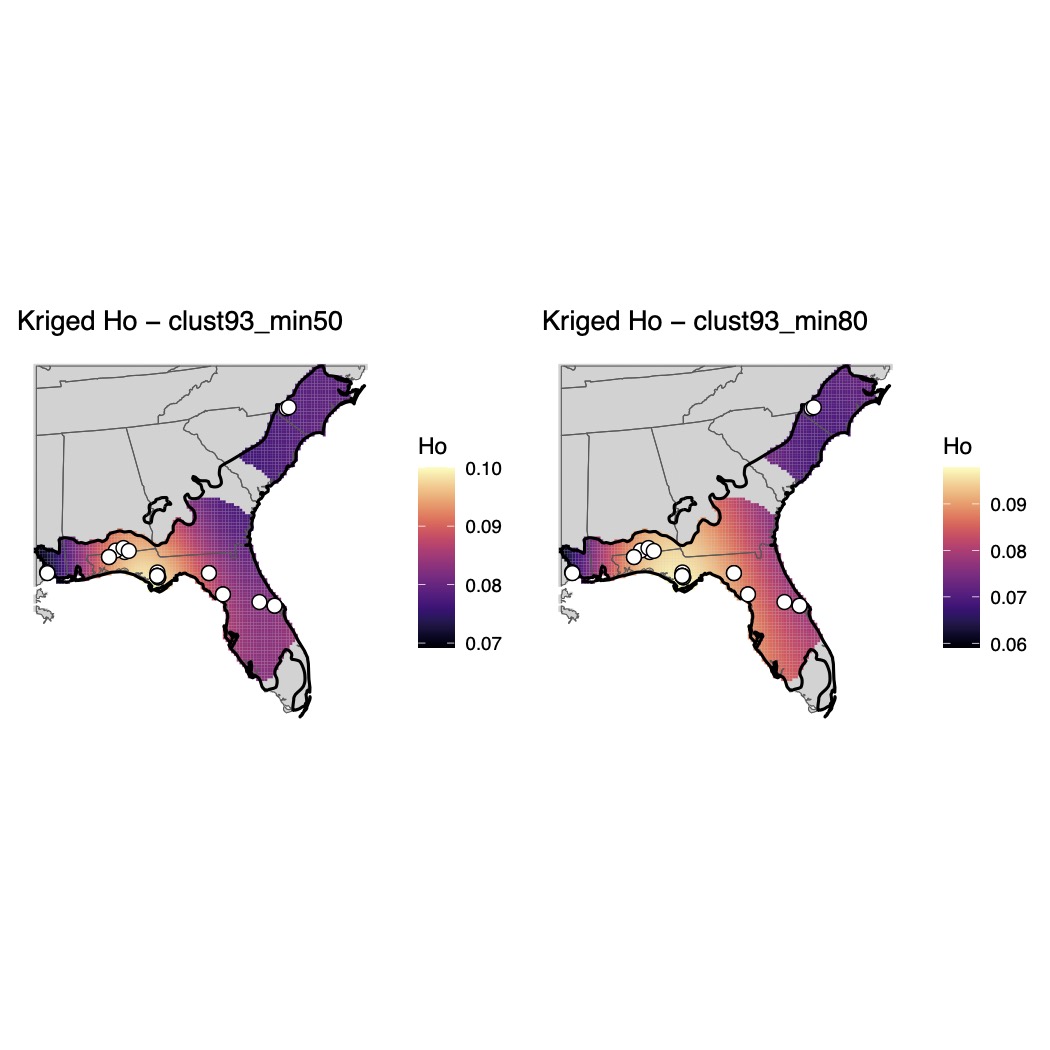

Supplementary Fig. 28.

Maps of genomic diversity for *Pseudacris nigrita* generated with the *WINGEN* R package.

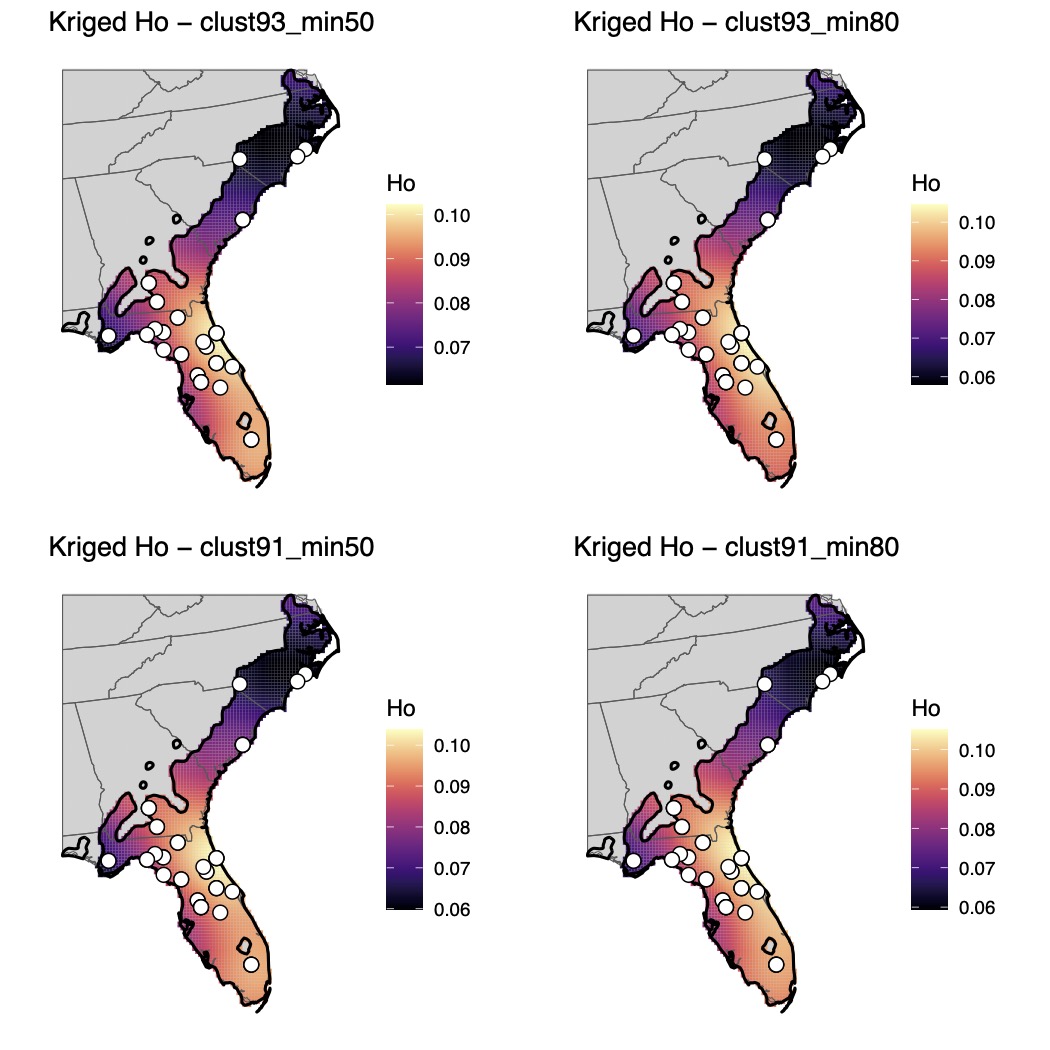

Supplementary Fig. 29.

Maps of genomic diversity for *Pseudacris ocularis* generated with the *WINGEN* R package.

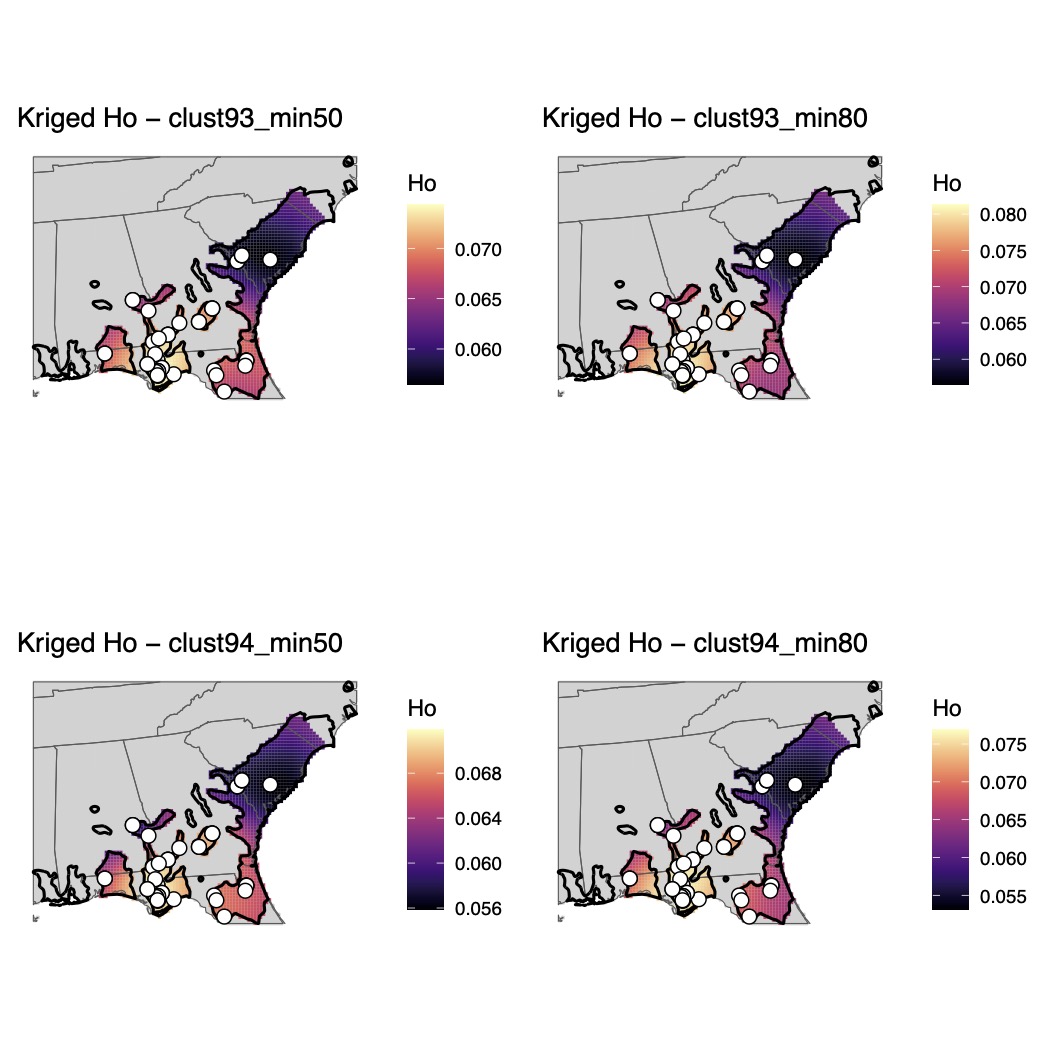

Supplementary Fig. 30.

Maps of genomic diversity for *Pseudacris ornata* generated with the *WINGEN* R package.

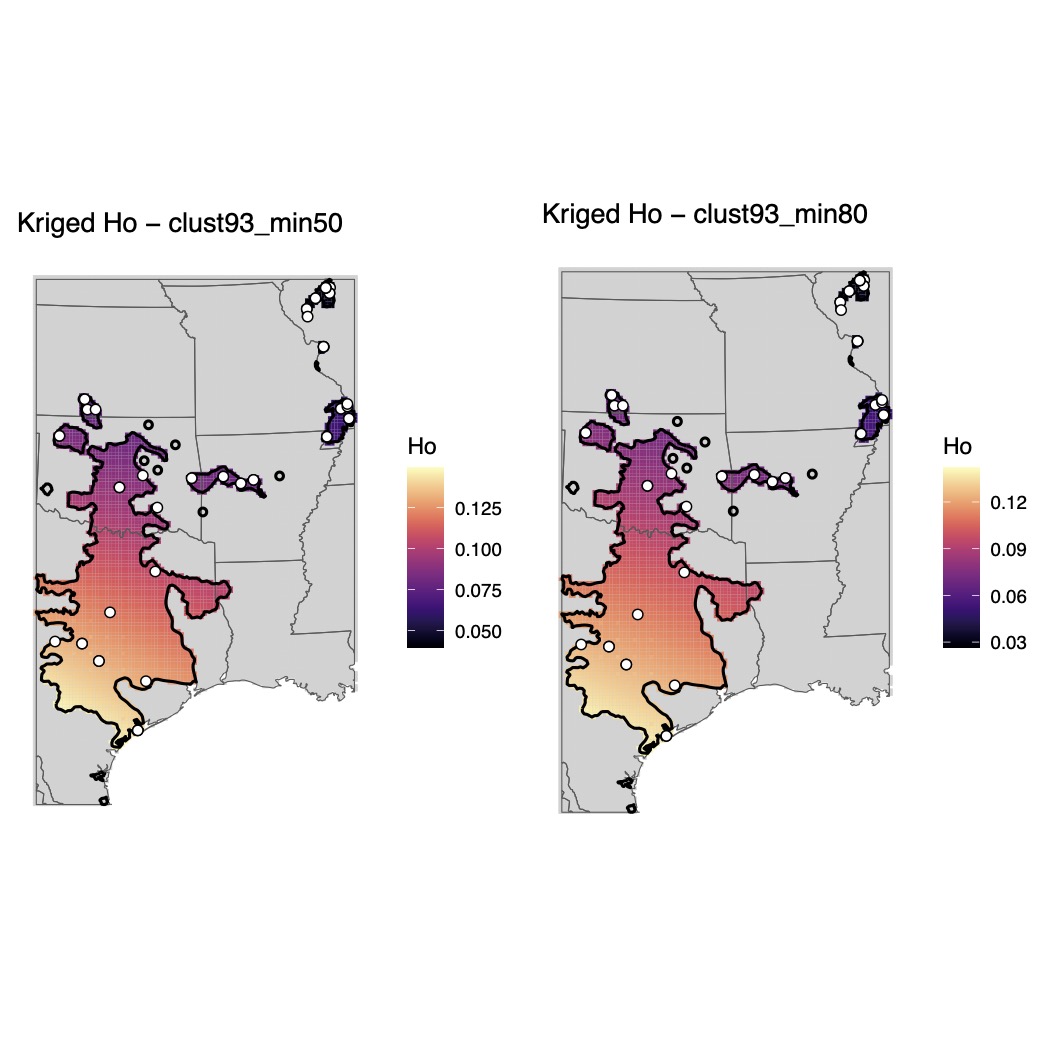

Supplementary Fig. 31.

Maps of genomic diversity for *Pseudacris streckeri* generated with the *WINGEN* R package.

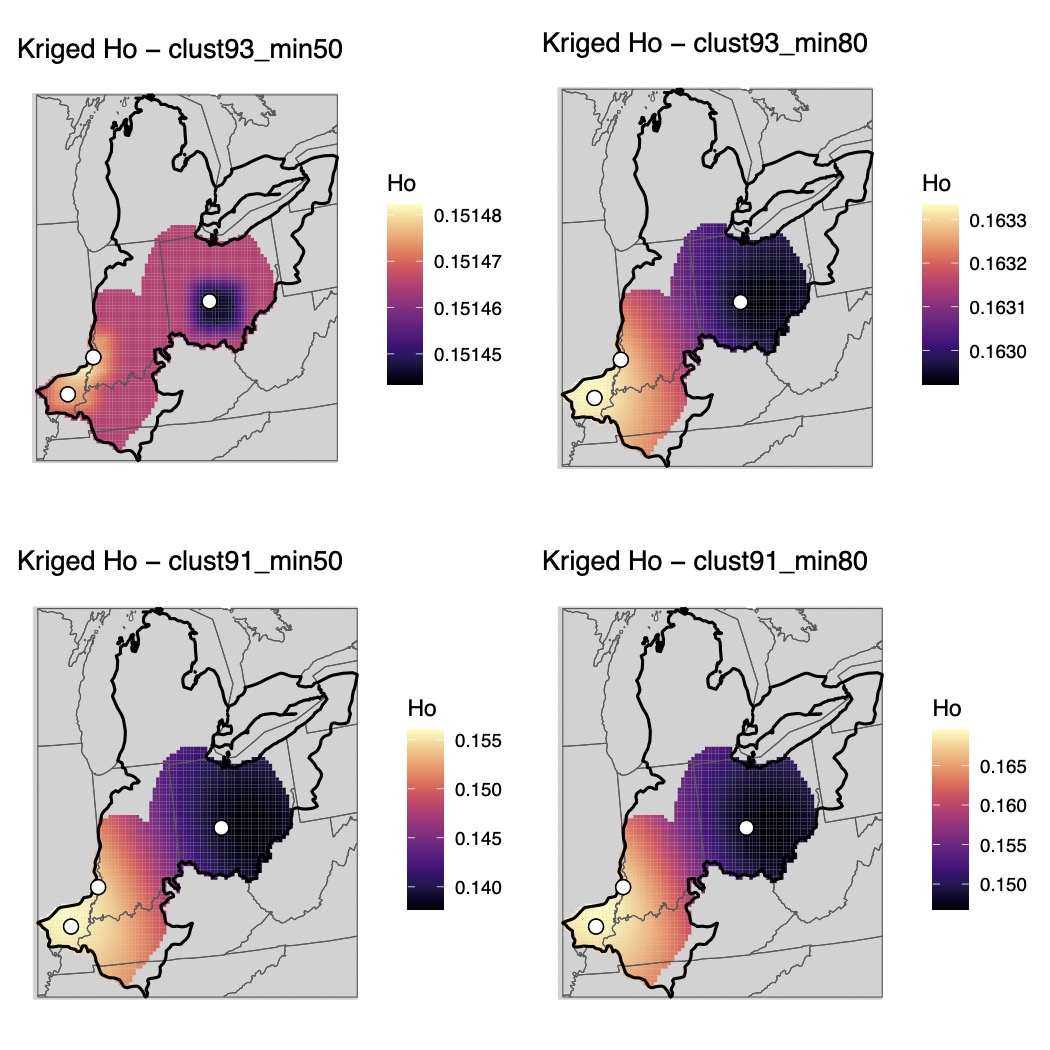

Supplementary Fig. 32.

Maps of genomic diversity for *Pseudacris triseriata* generated with the *WINGEN* R package.

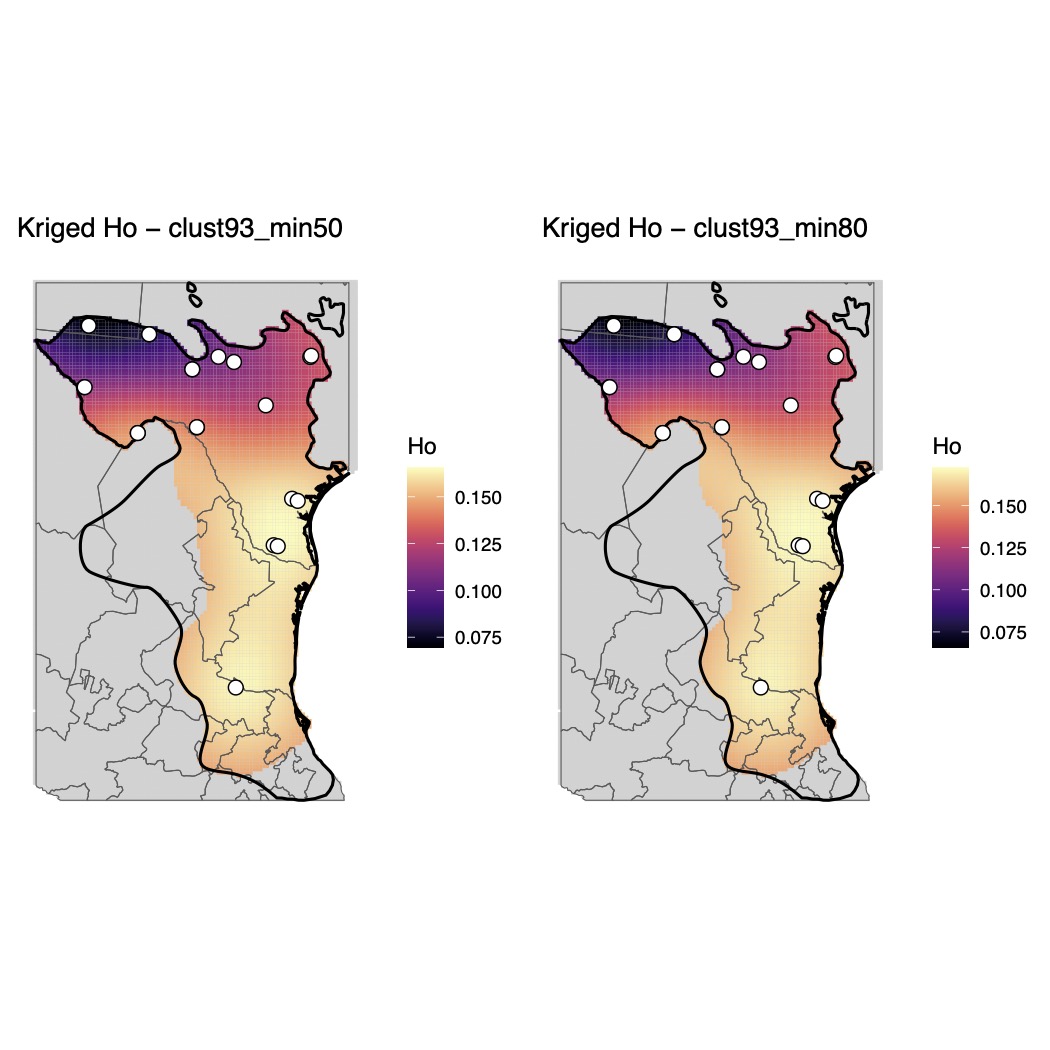

**Supplementary Fig. 33.**

Maps of genomic diversity for *Rana berlandieri* generated with the *WINGEN* R package.

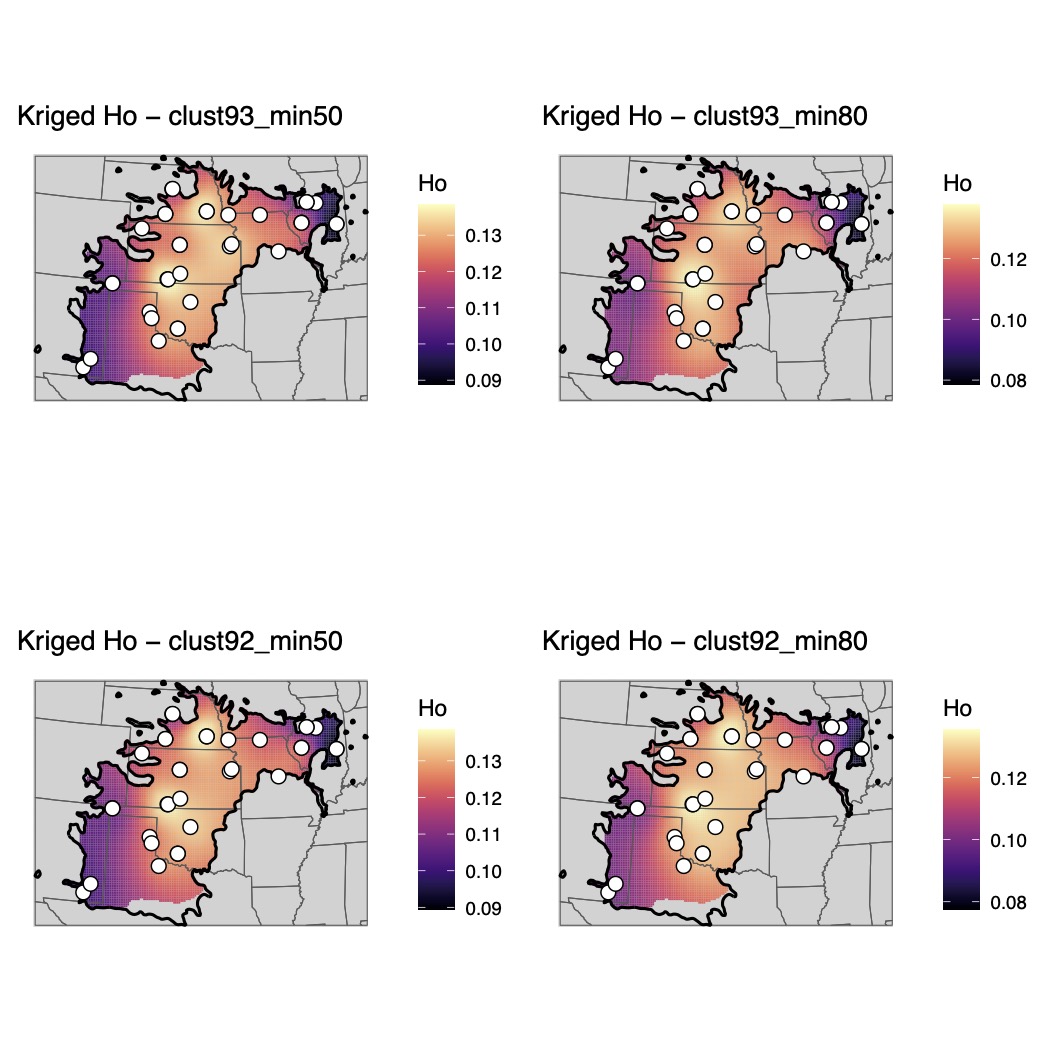

**Supplementary Fig. 34.**

Maps of genomic diversity for *Rana blairi* generated with the *WINGEN* R package.

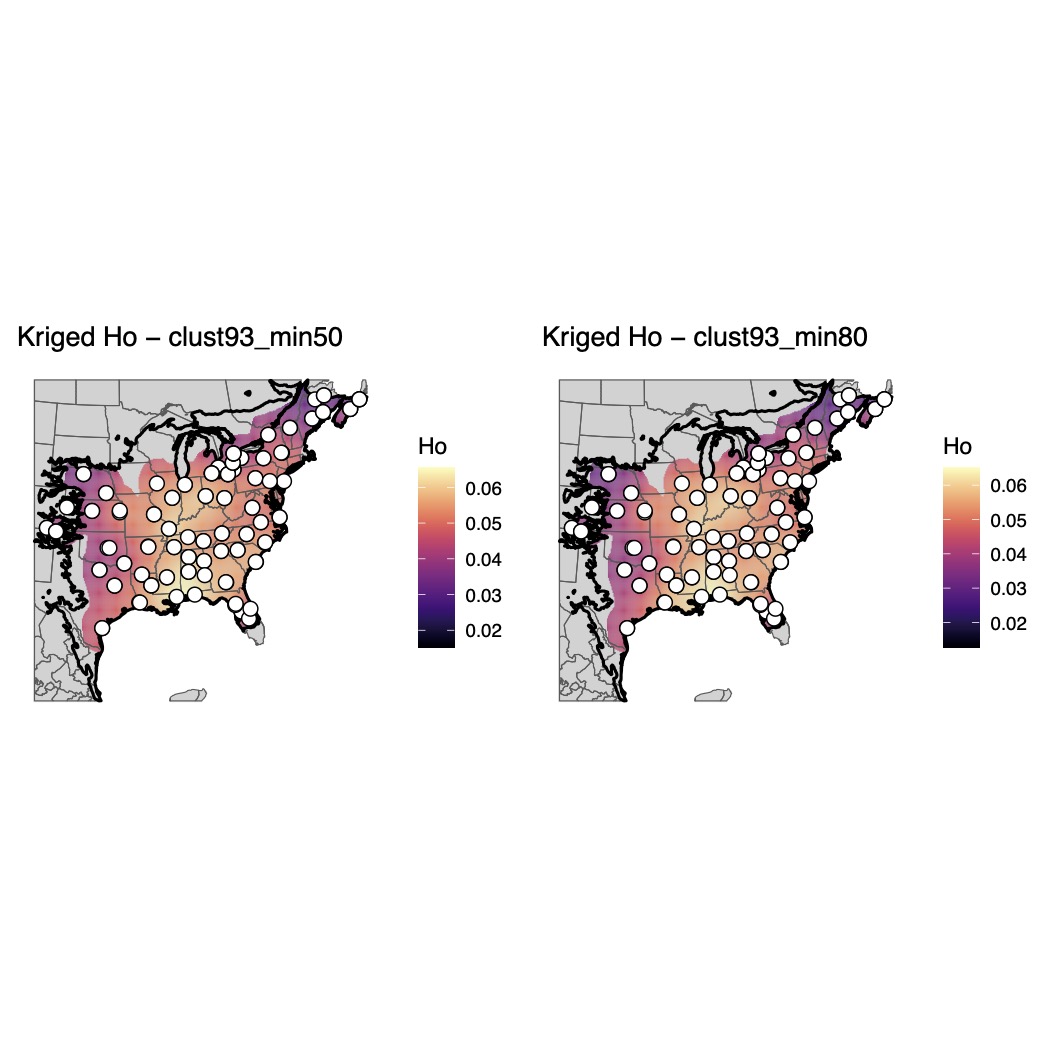

**Supplementary Fig. 35.**

Maps of genomic diversity for *Rana catesbeiana* generated with the *WINGEN* R package.

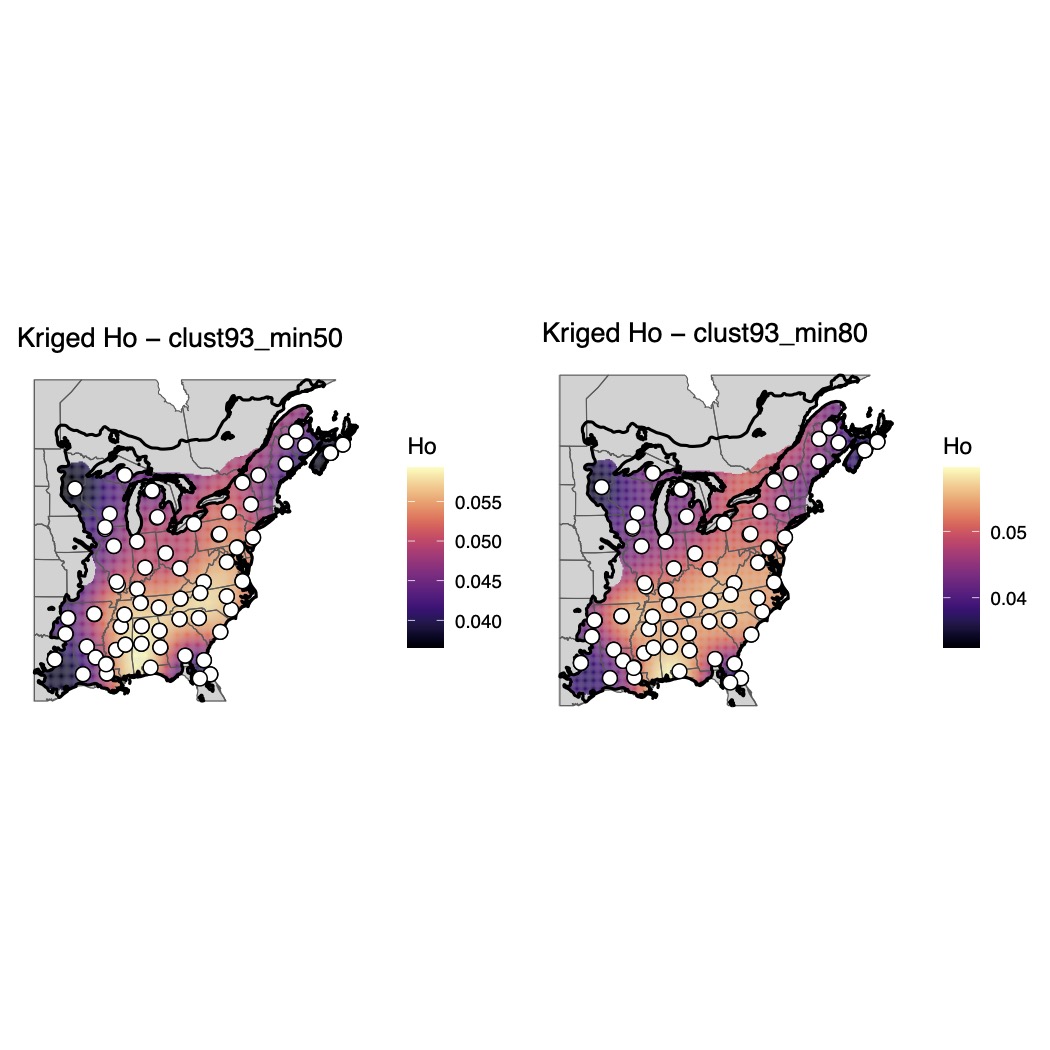

**Supplementary Fig. 36.**

Maps of genomic diversity for *Rana clamitans* generated with the *WINGEN* R package.

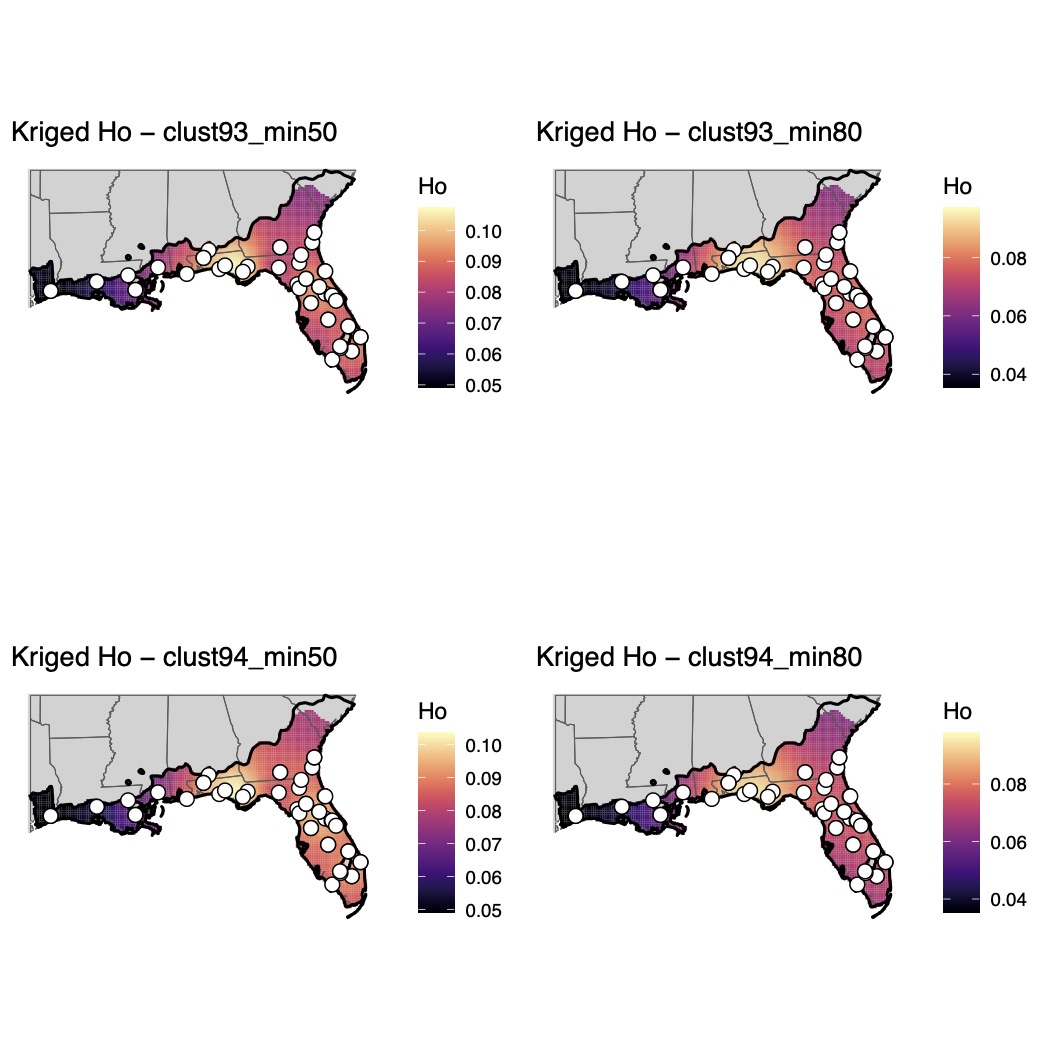

**Supplementary Fig. 37.**

Maps of genomic diversity for *Rana grylio* generated with the *WINGEN* R package.

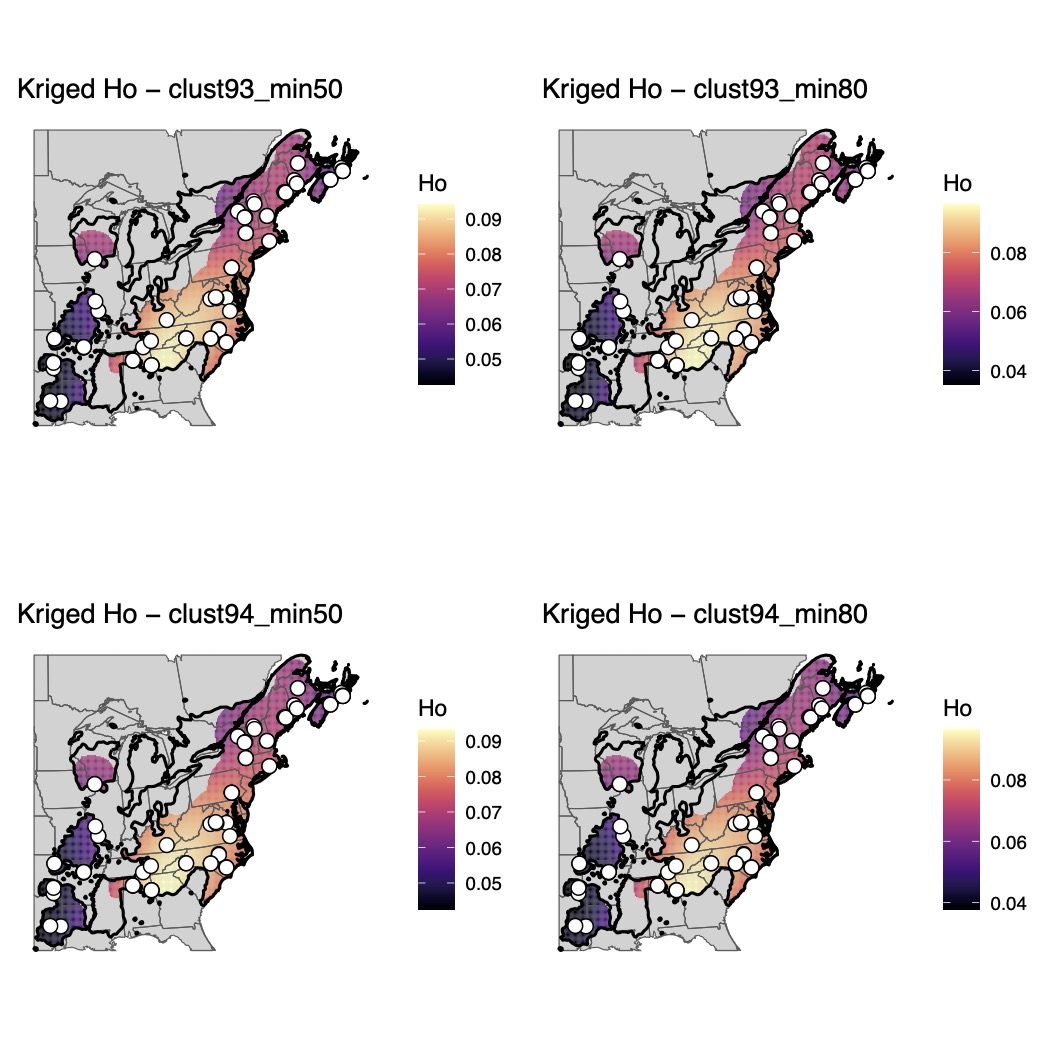

**Supplementary Fig. 38.**

Maps of genomic diversity for *Rana palustris* generated with the *WINGEN* R package.

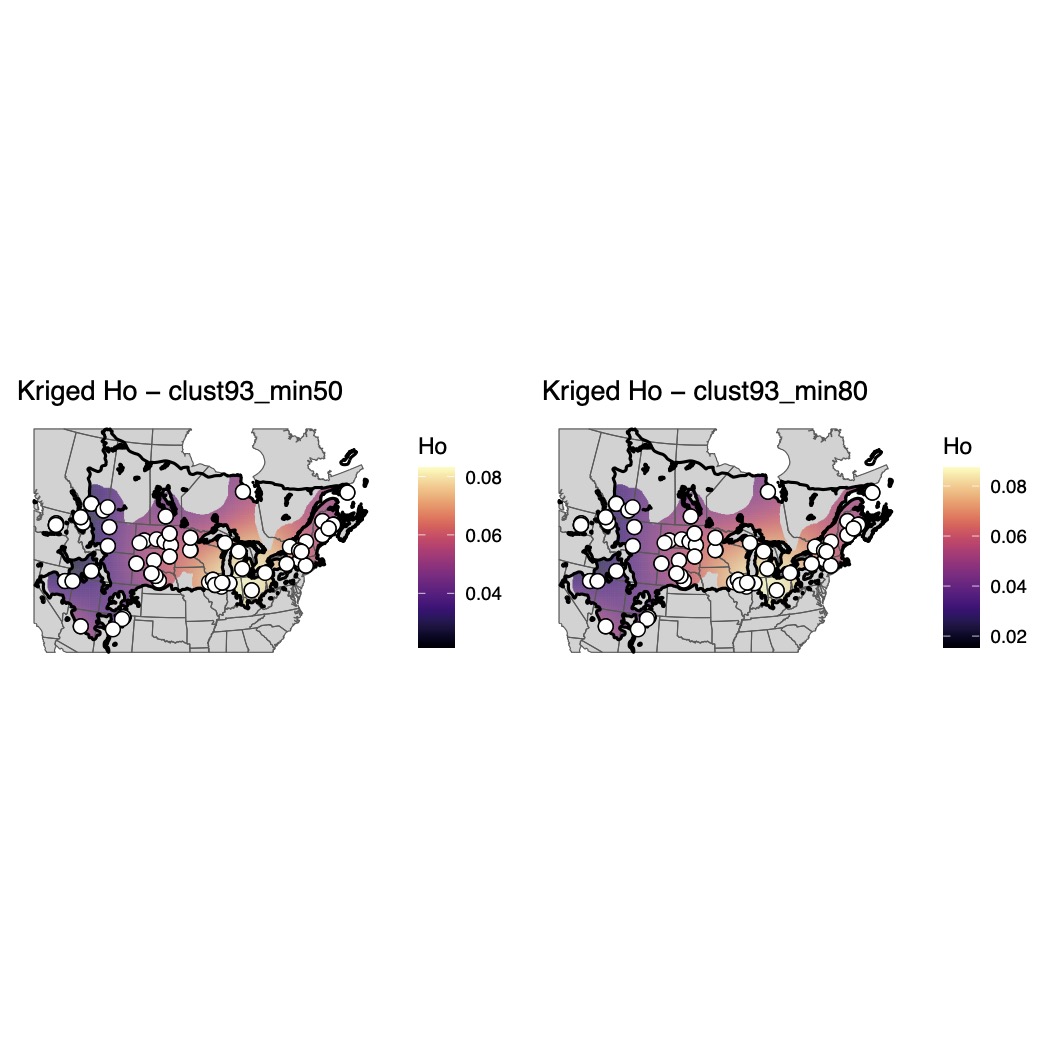
Supplementary Fig. 39.

Maps of genomic diversity for *Rana pipiens* generated with the *WINGEN* R package.

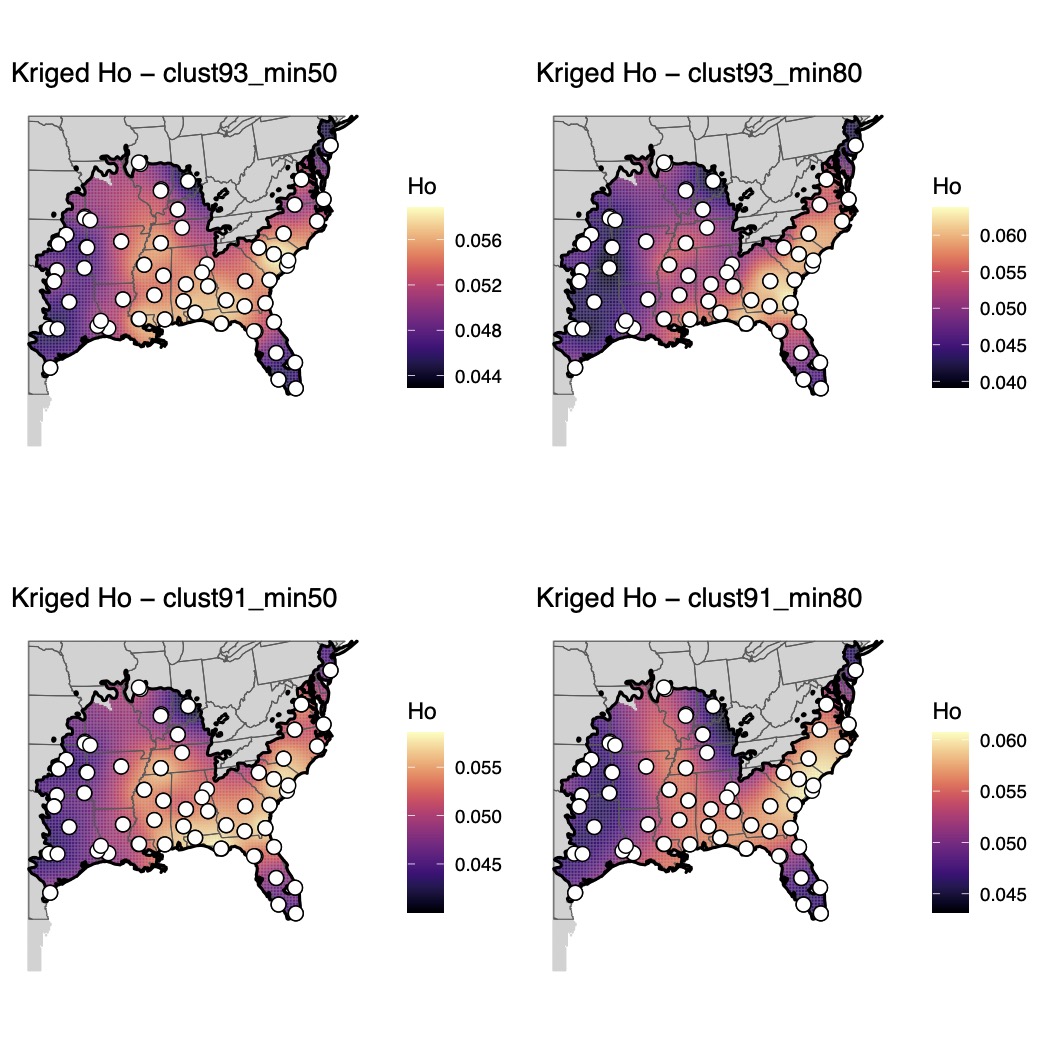

Supplementary Fig. 40.

Maps of genomic diversity for *Rana sphenocephala* generated with the *WINGEN* R package.

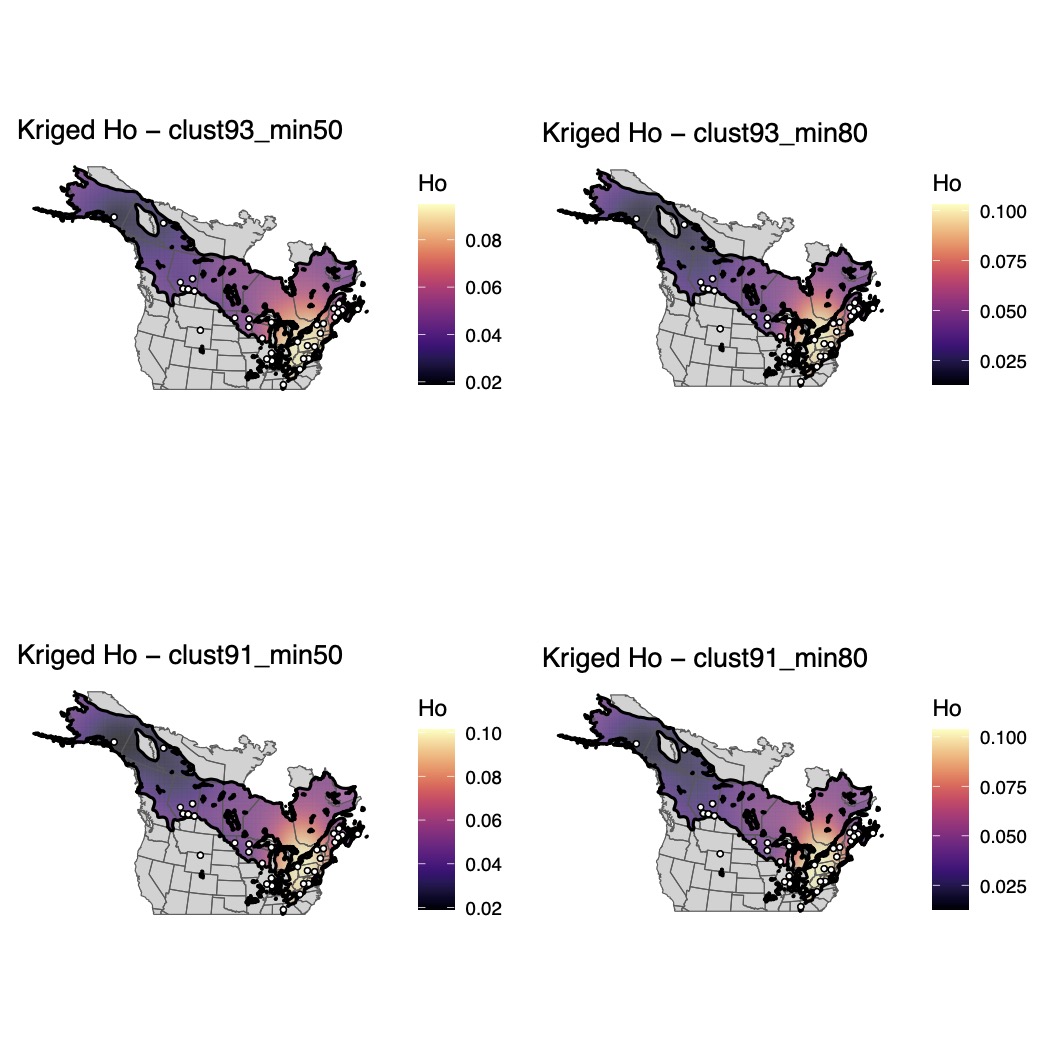

Supplementary Fig. 41.

Maps of genomic diversity for *Rana sylvatica* generated with the *WINGEN* R package.

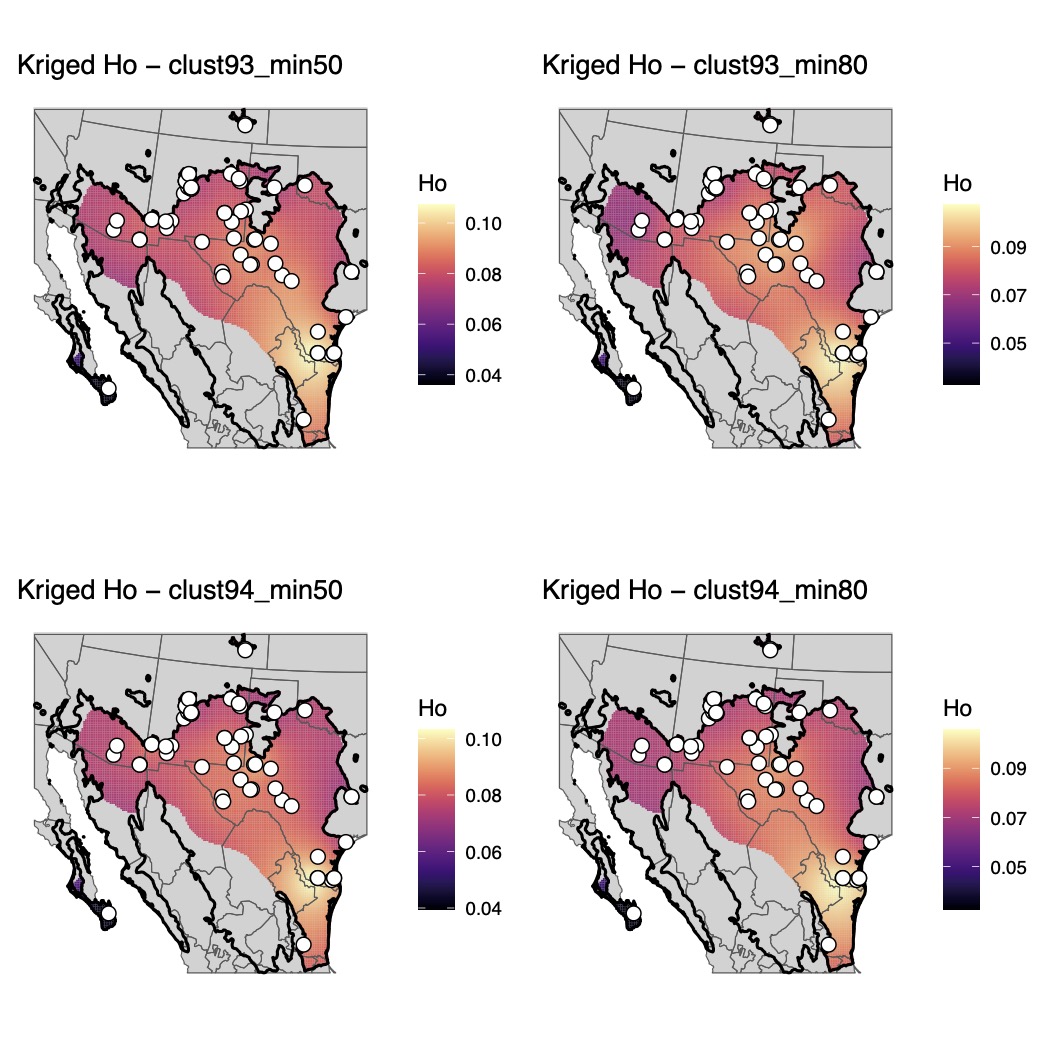

Supplementary Fig. 42.

Maps of genomic diversity for *Scaphiopus couchii* generated with the *WINGEN* R package.

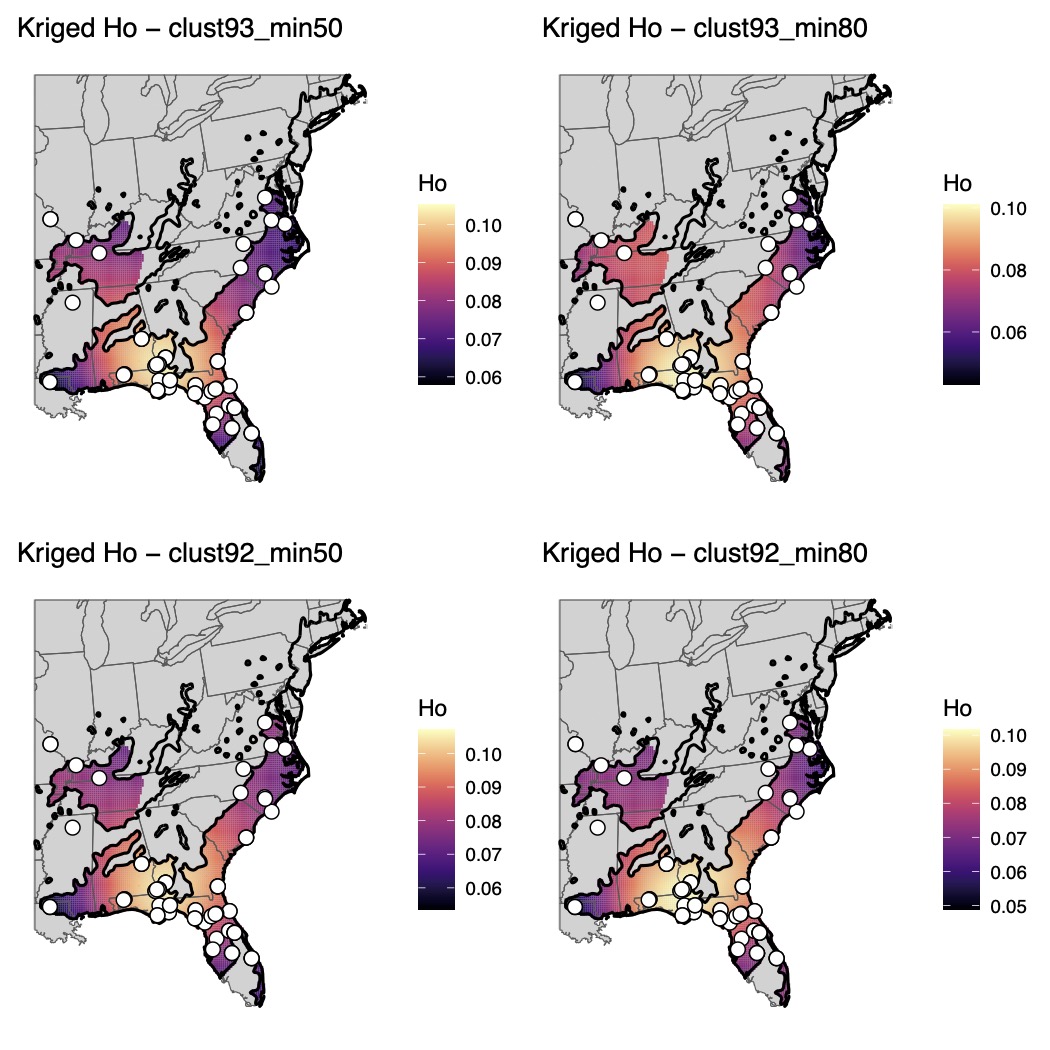

Supplementary Fig. 43.

Maps of genomic diversity for *Scaphiopus holbrookii* generated with the *WINGEN* R package.

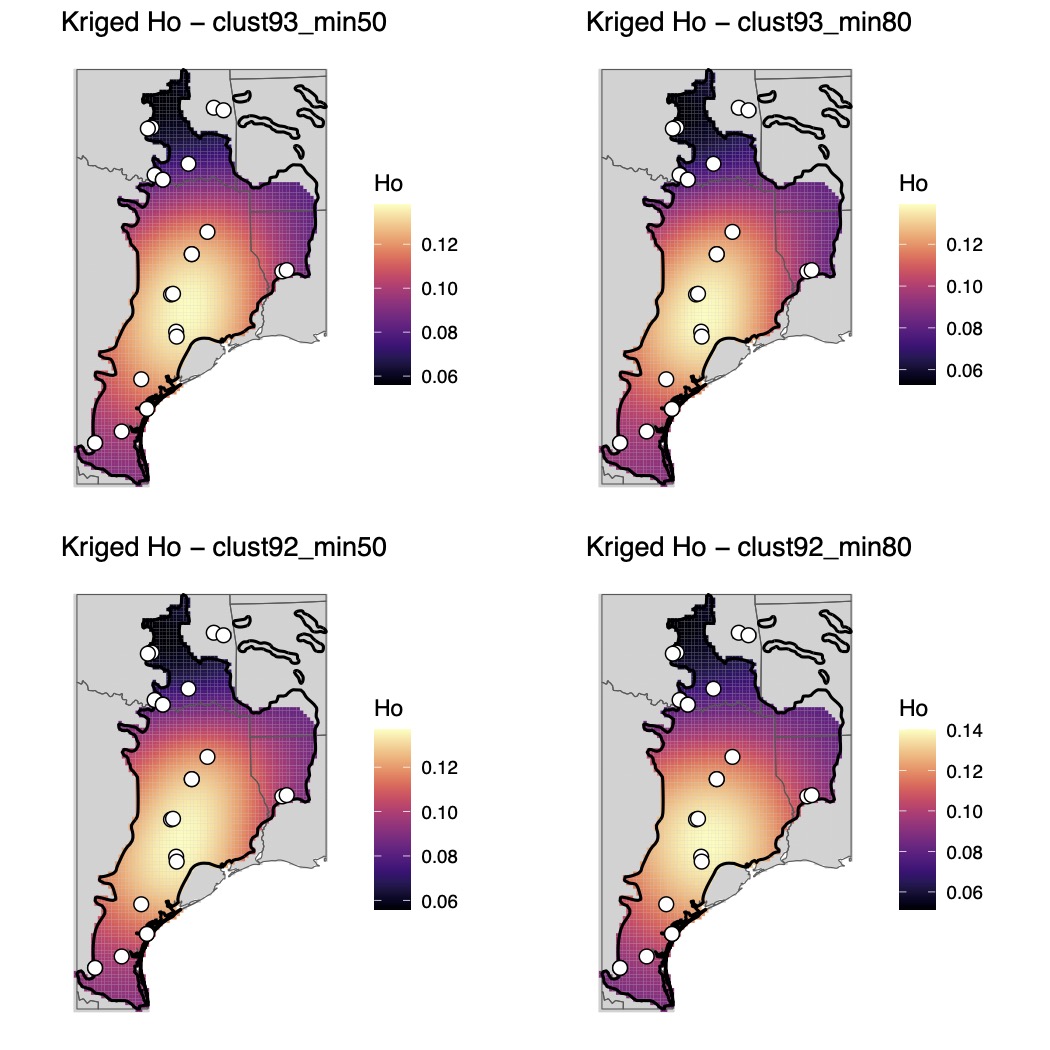

Supplementary Fig. 44.

Maps of genomic diversity for *Scaphiopus hurterii* generated with the *WINGEN* R package.

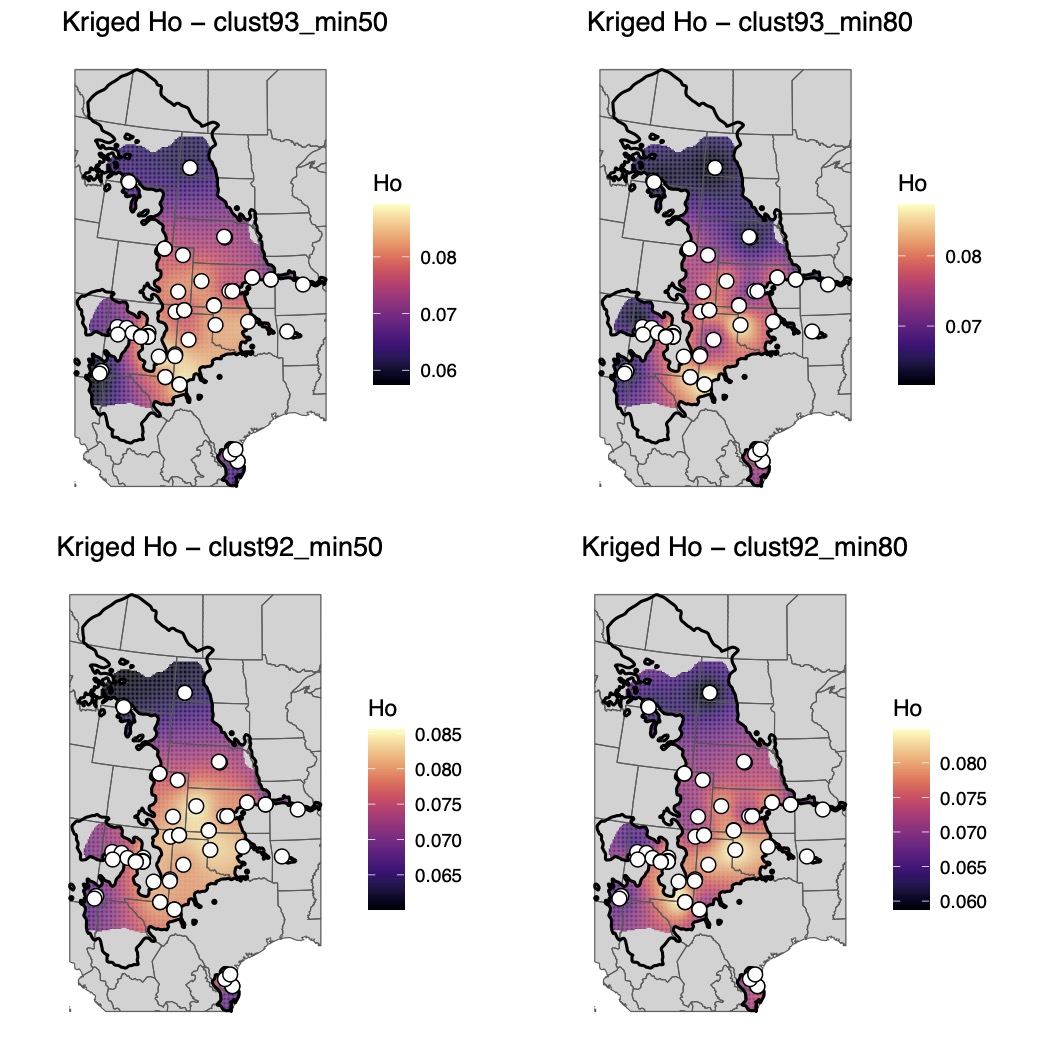

Supplementary Fig. 45.

Maps of genomic diversity for *Spea bombifrons* generated with the *WINGEN* R package.

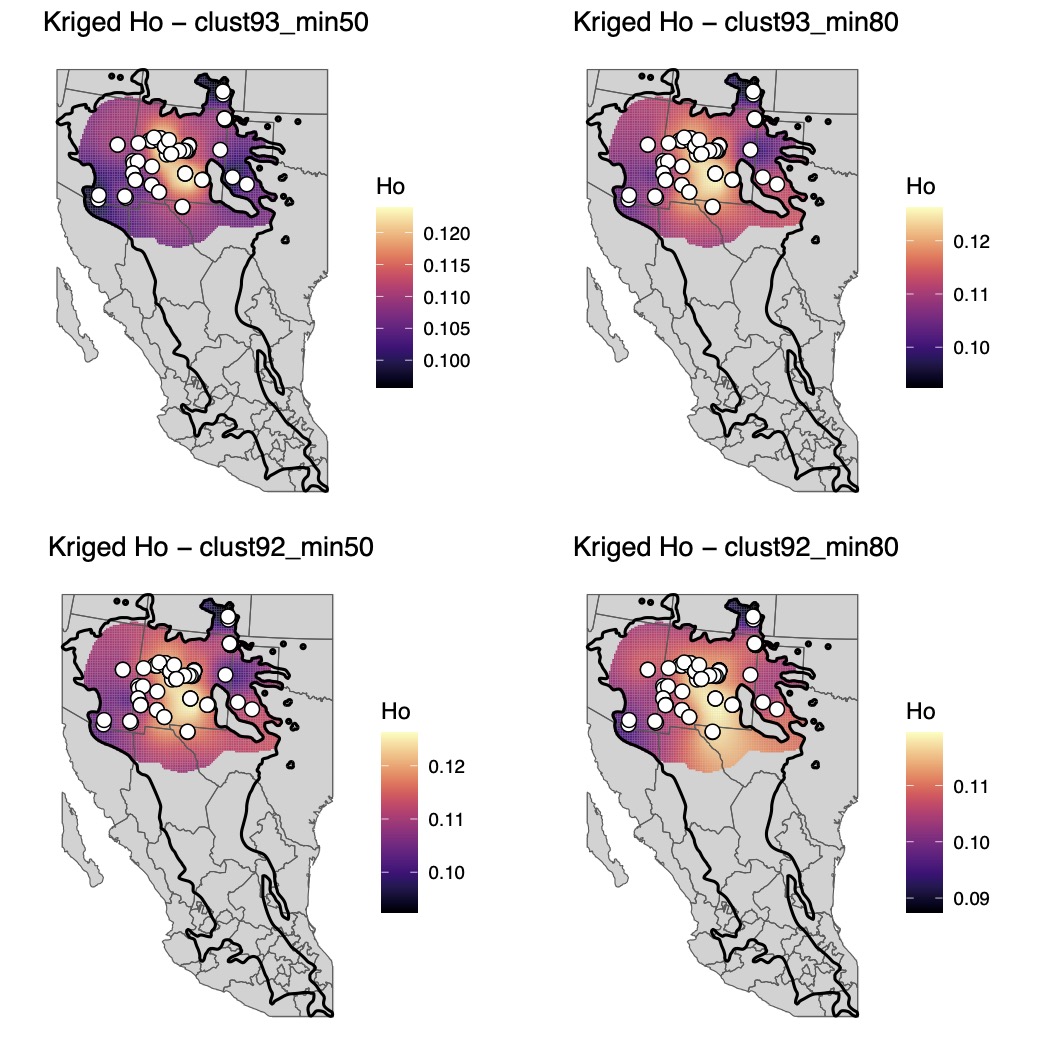

Supplementary Fig. 46.

Maps of genomic diversity for *Spea multiplicata* generated with the *WINGEN* R package.

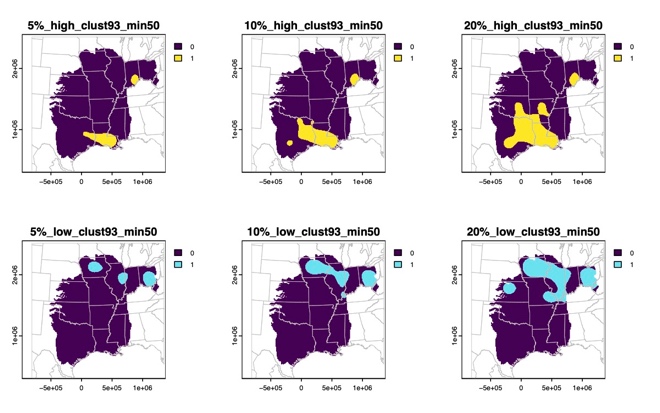

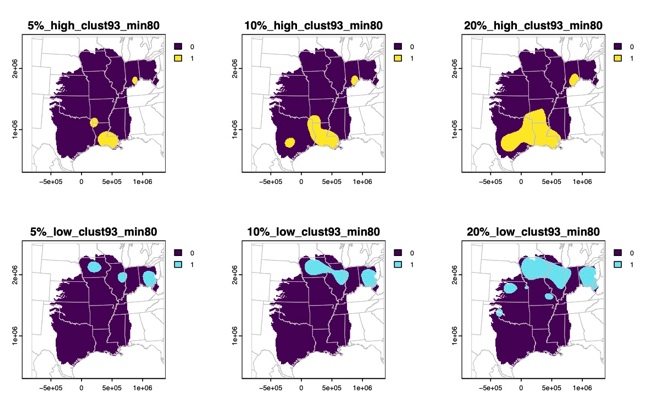

Supplementary Fig. 47.

Binary maps of the highest (yellow) or lowest (blue) genomic diversity for *Acris blanchardi* based on 5%, 10%, or 20% quantile thresholds.

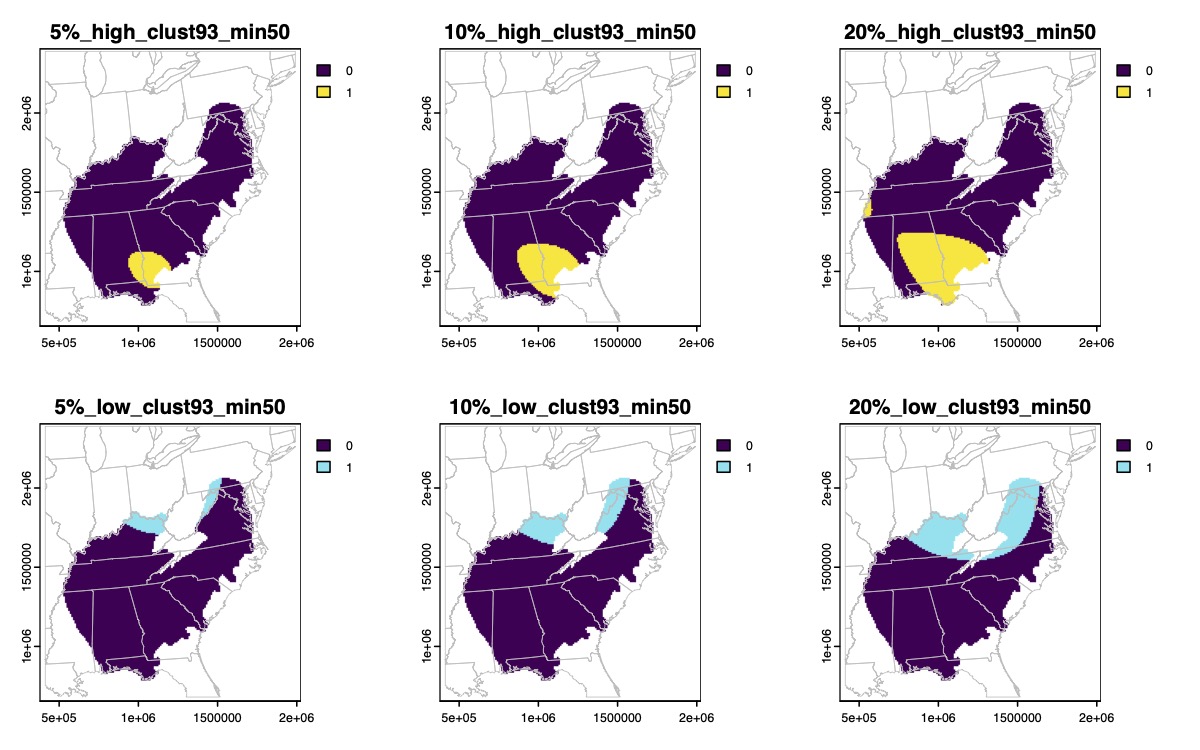

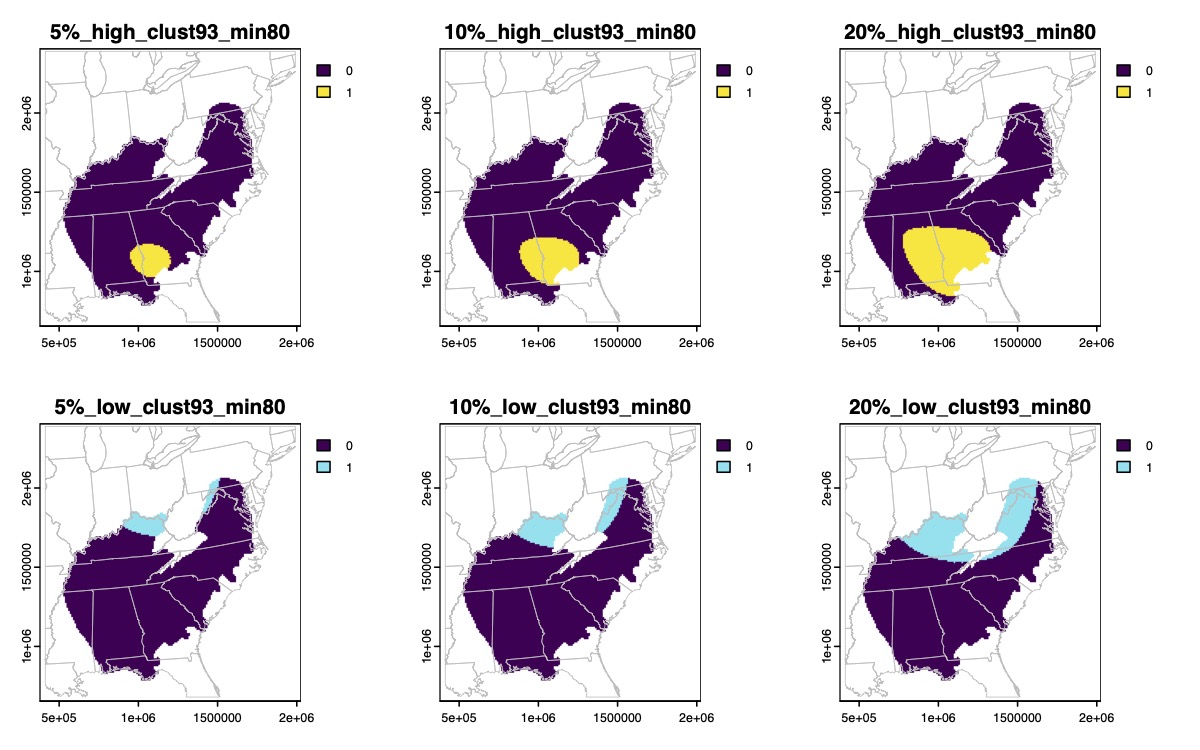

Supplementary Fig. 48.

Binary maps of the highest (yellow) or lowest (blue) genomic diversity for *Acris crepitans* based on 5%, 10%, or 20% quantile thresholds.

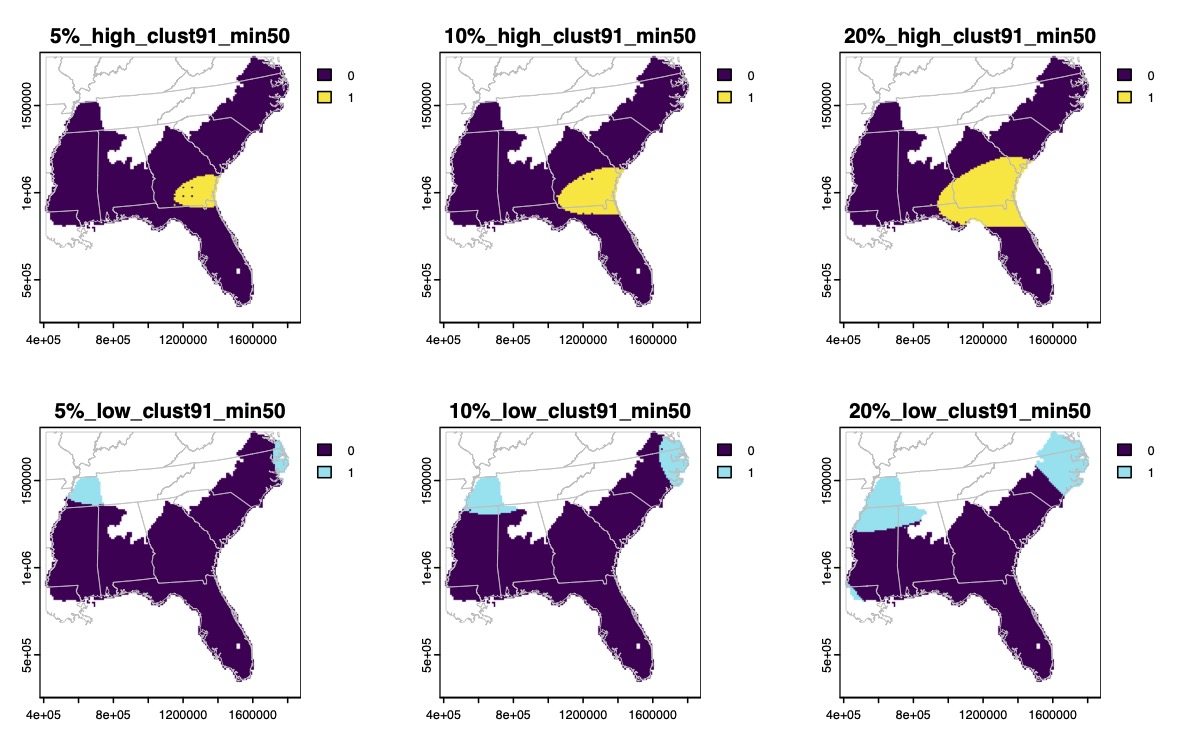

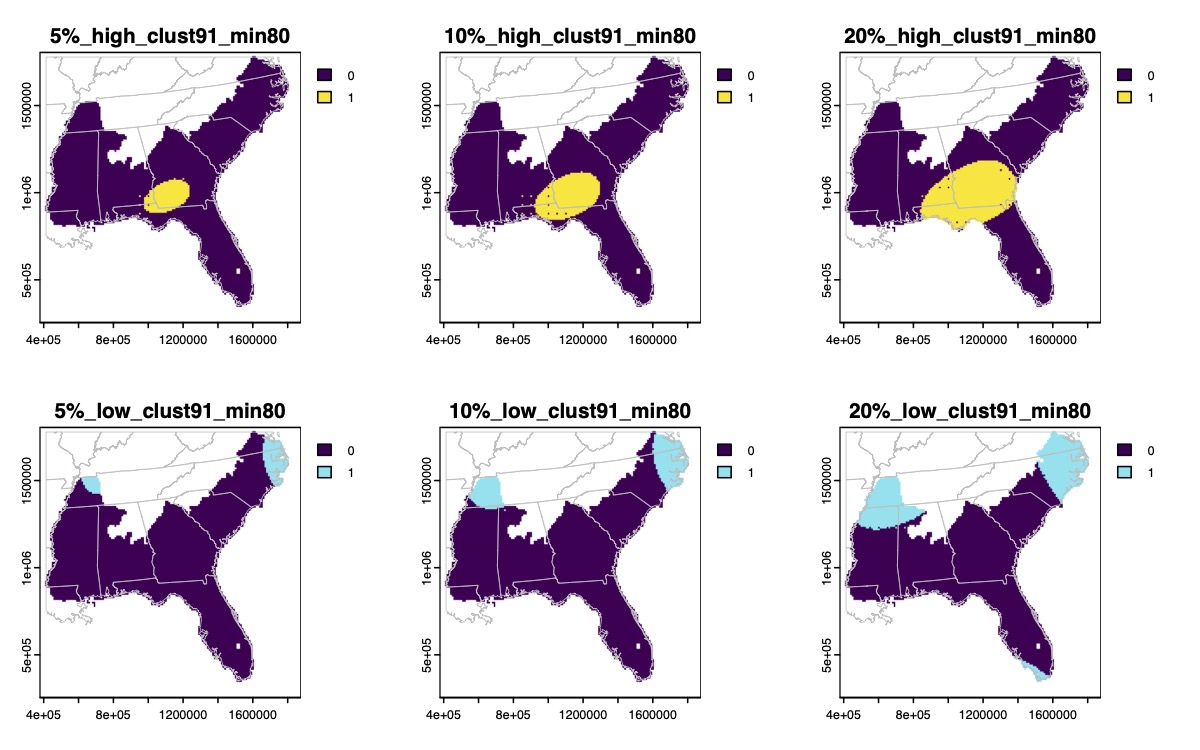

Supplementary Fig. 49.

Binary maps of the highest (yellow) or lowest (blue) genomic diversity for *Acris gryllus* based on 5%, 10%, or 20% quantile thresholds.

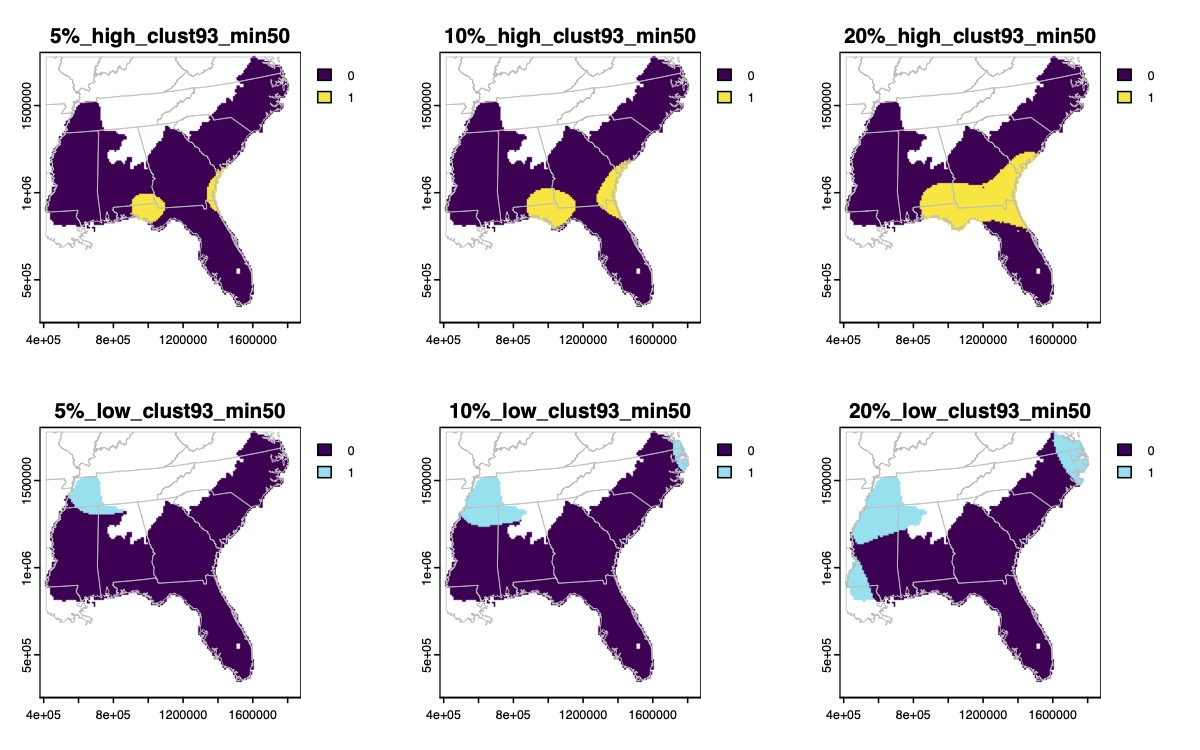

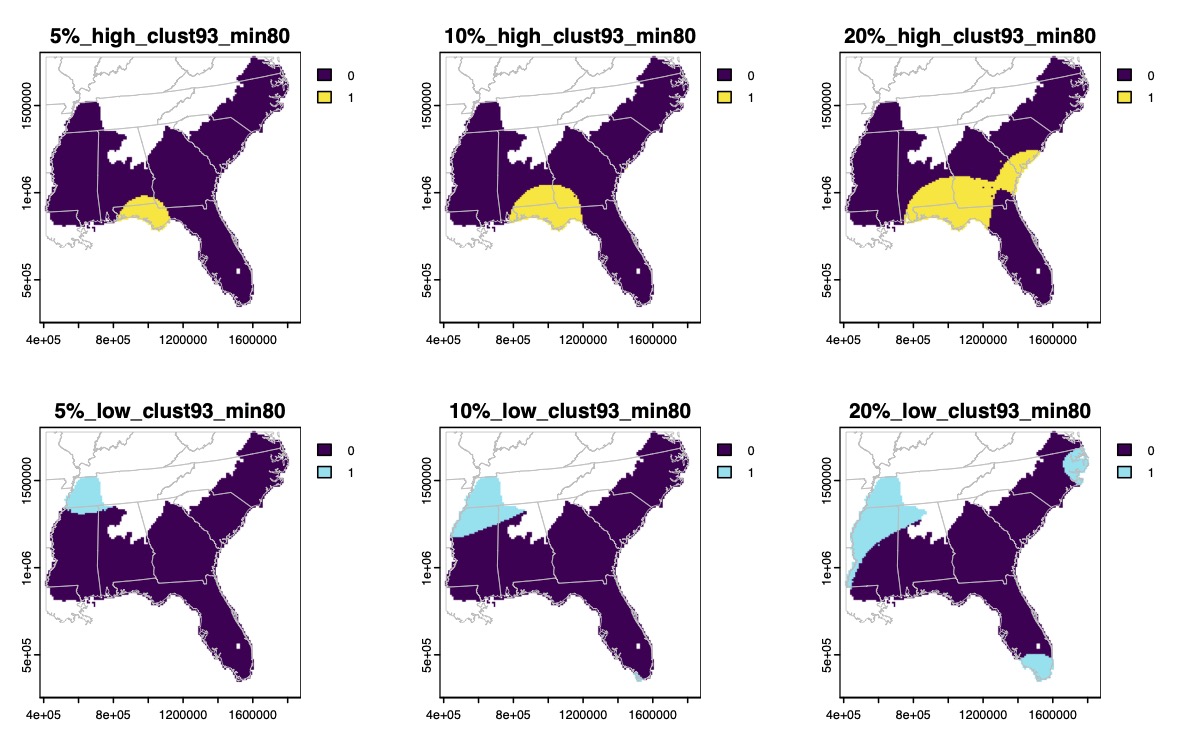

Supplementary Fig. 49. (continued)

Binary maps of the highest (yellow) or lowest (blue) genomic diversity for *Acris gryllus* based on 5%, 10%, or 20% quantile thresholds.

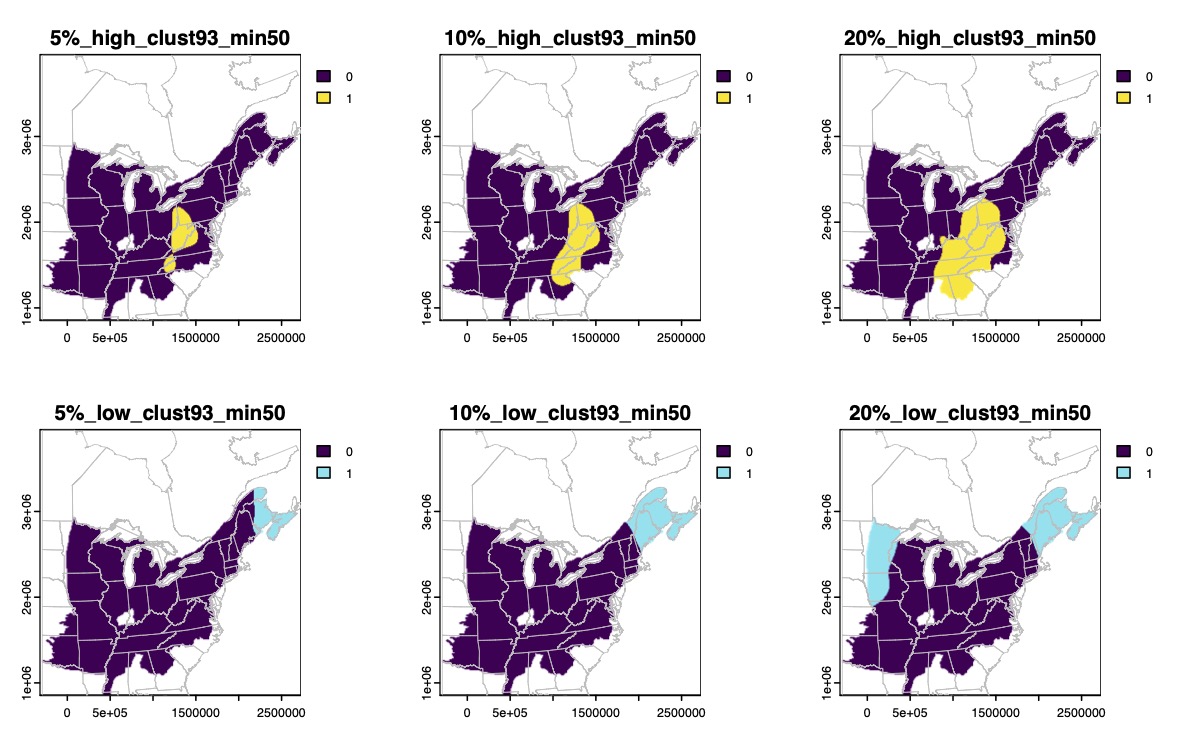

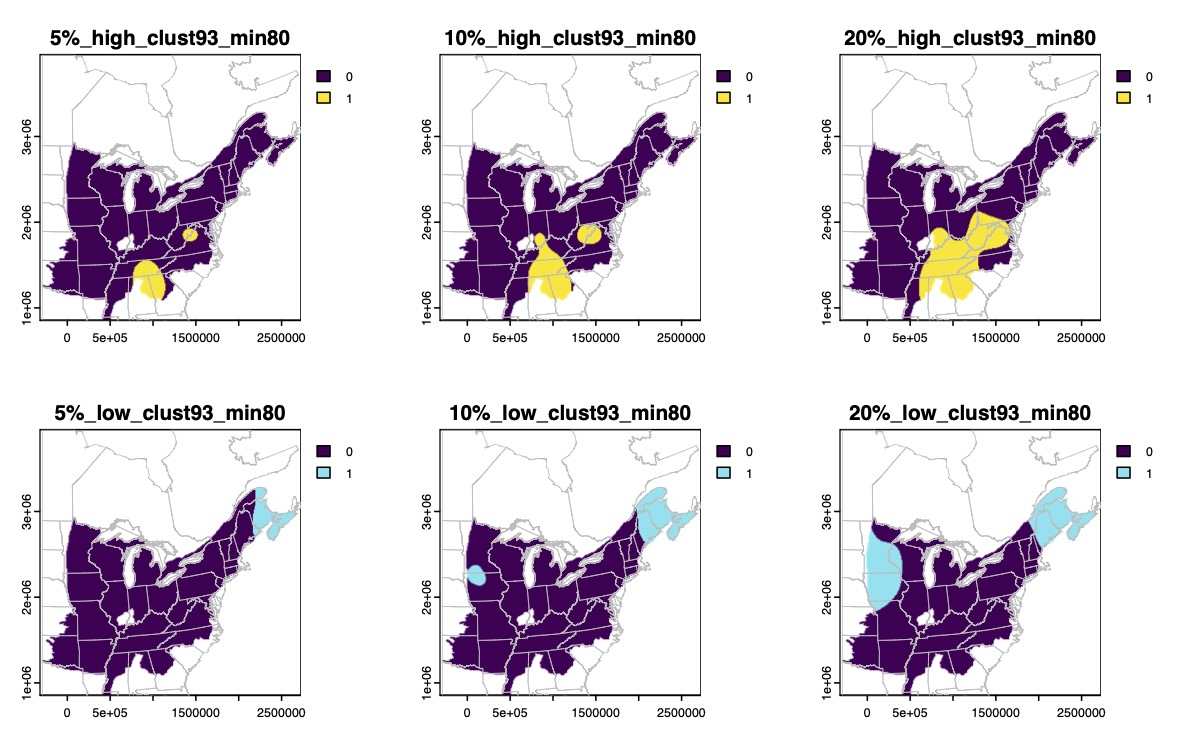

Supplementary Fig. 50.

Binary maps of the highest (yellow) or lowest (blue) genomic diversity for *Anaxyrus americanus* based on 5%, 10%, or 20% quantile thresholds.

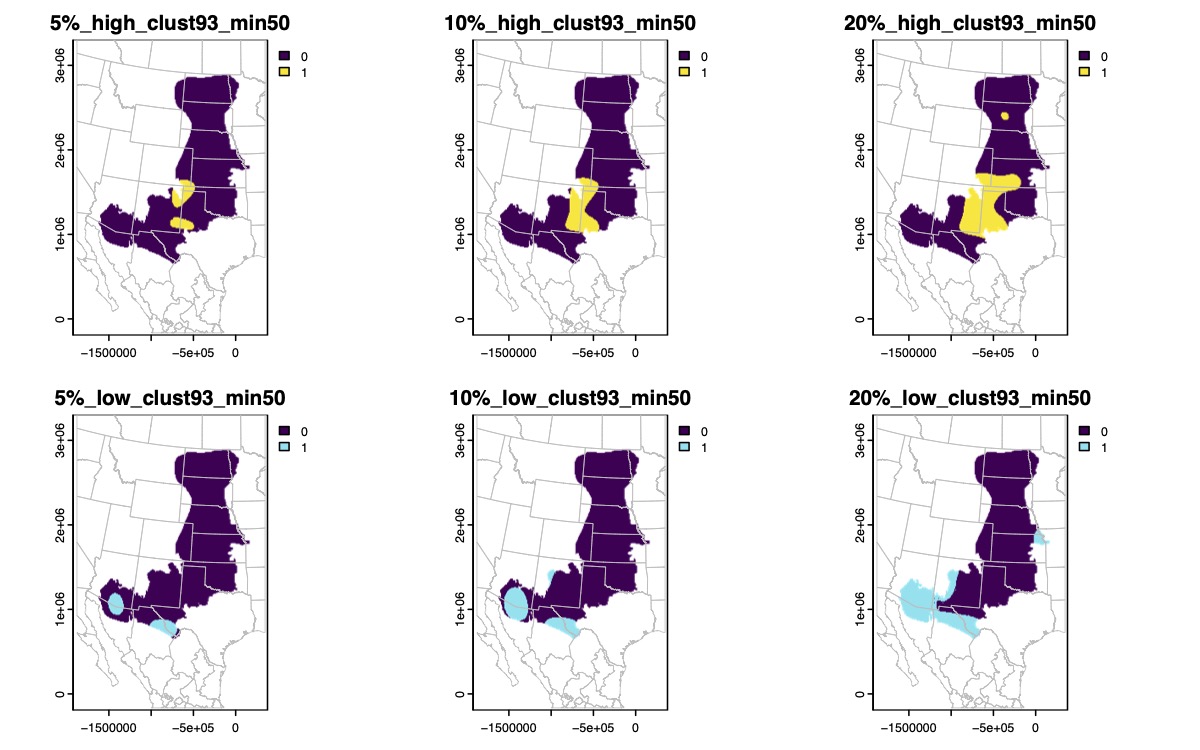

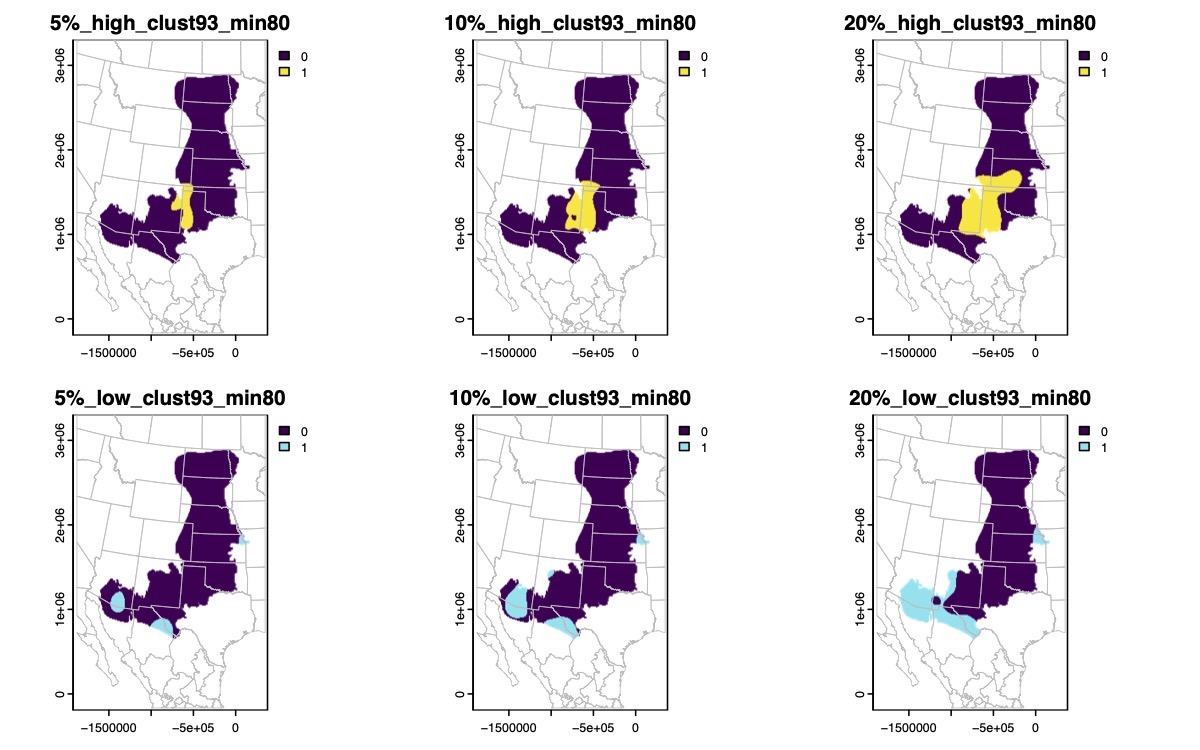

Supplementary Fig. 51.

Binary maps of the highest (yellow) or lowest (blue) genomic diversity for *Anaxyrus cognatus* based on 5%, 10%, or 20% quantile thresholds.

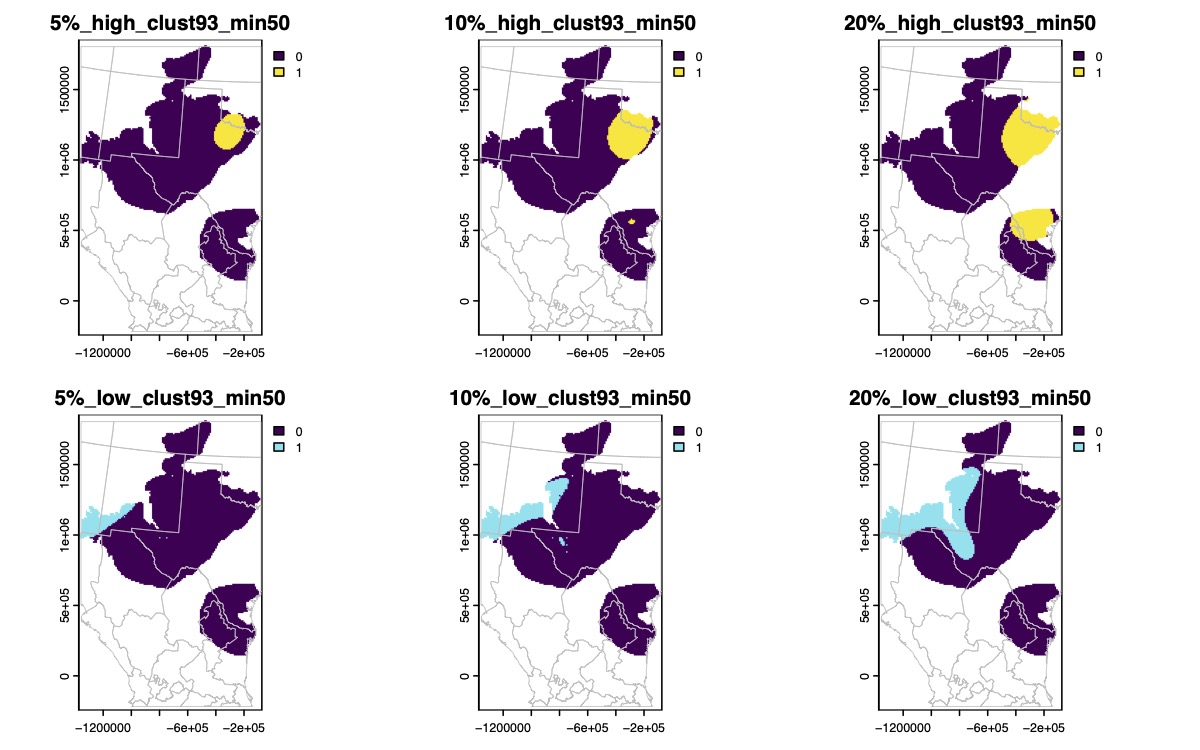

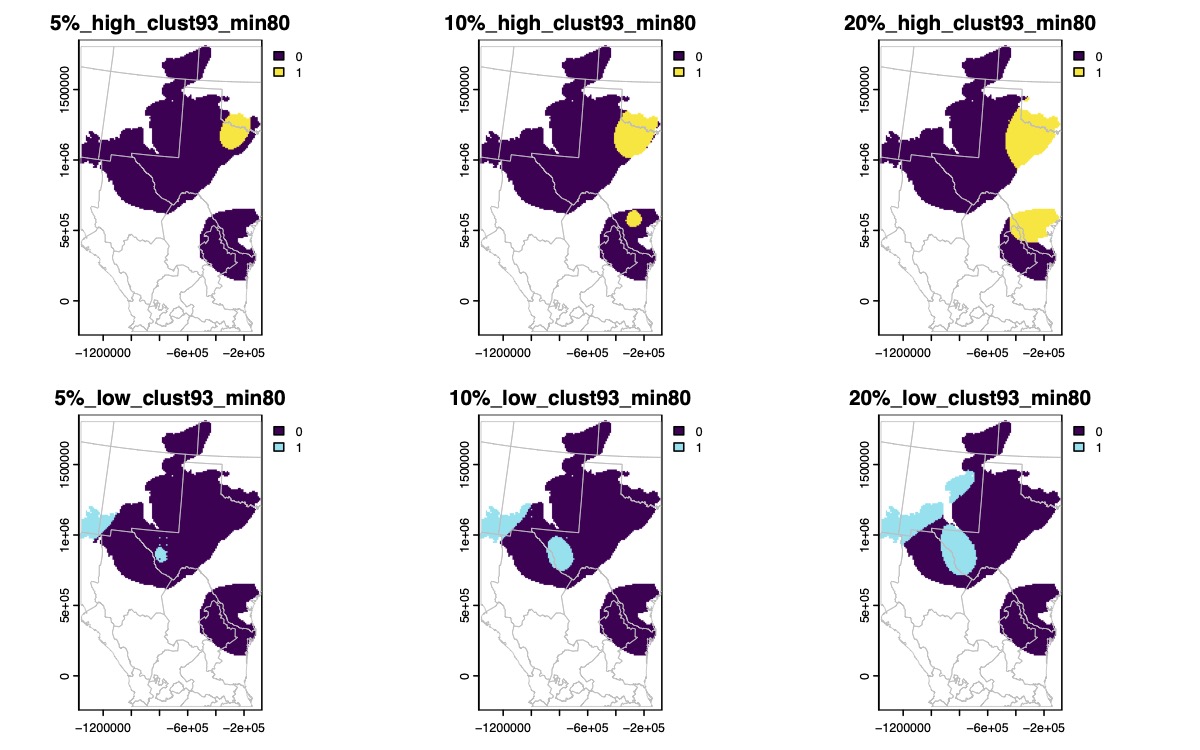

Supplementary Fig. 52.

Binary maps of the highest (yellow) or lowest (blue) genomic diversity for *Anaxyrus debilis* based on 5%, 10%, or 20% quantile thresholds.

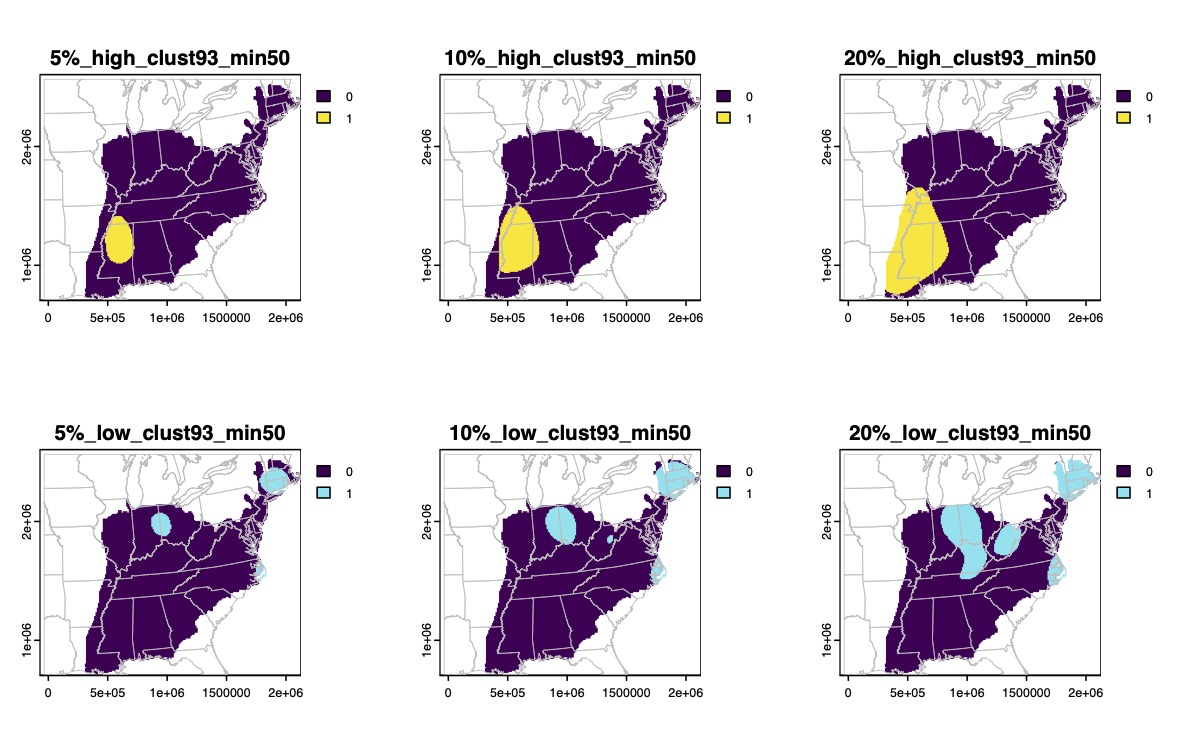

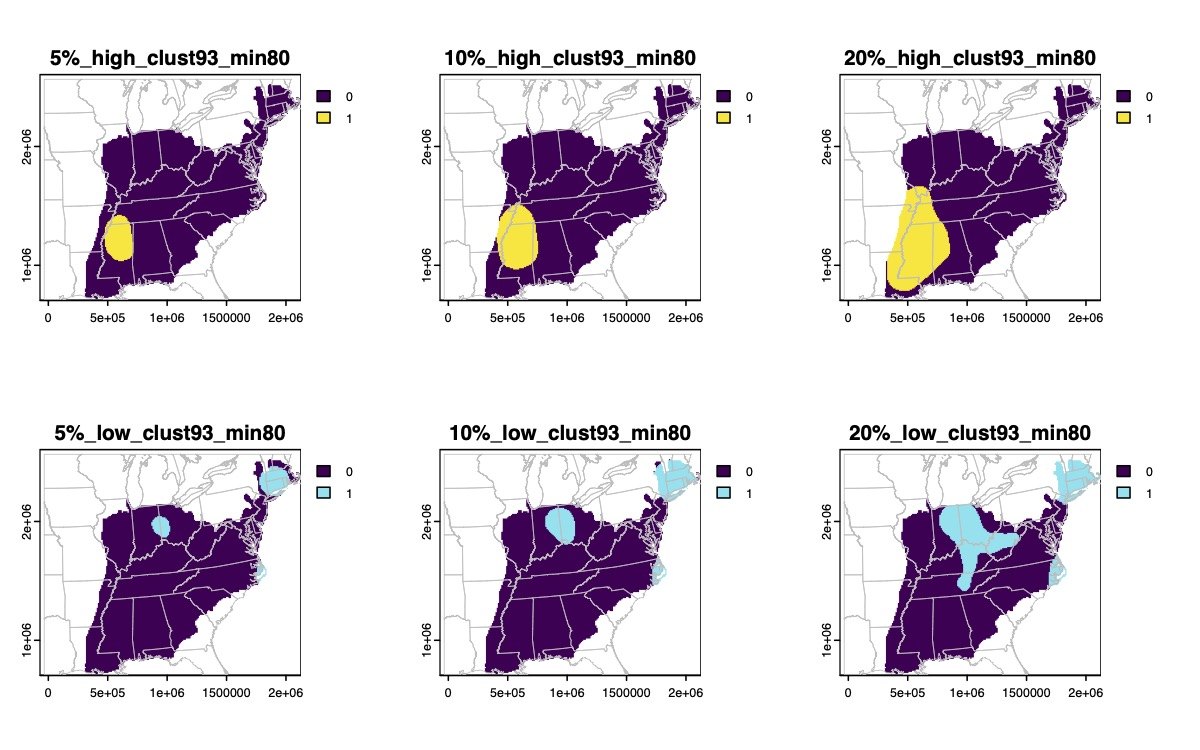

Supplementary Fig. 53.

Binary maps of the highest (yellow) or lowest (blue) genomic diversity for *Anaxyrus fowleri* based on 5%, 10%, or 20% quantile thresholds.

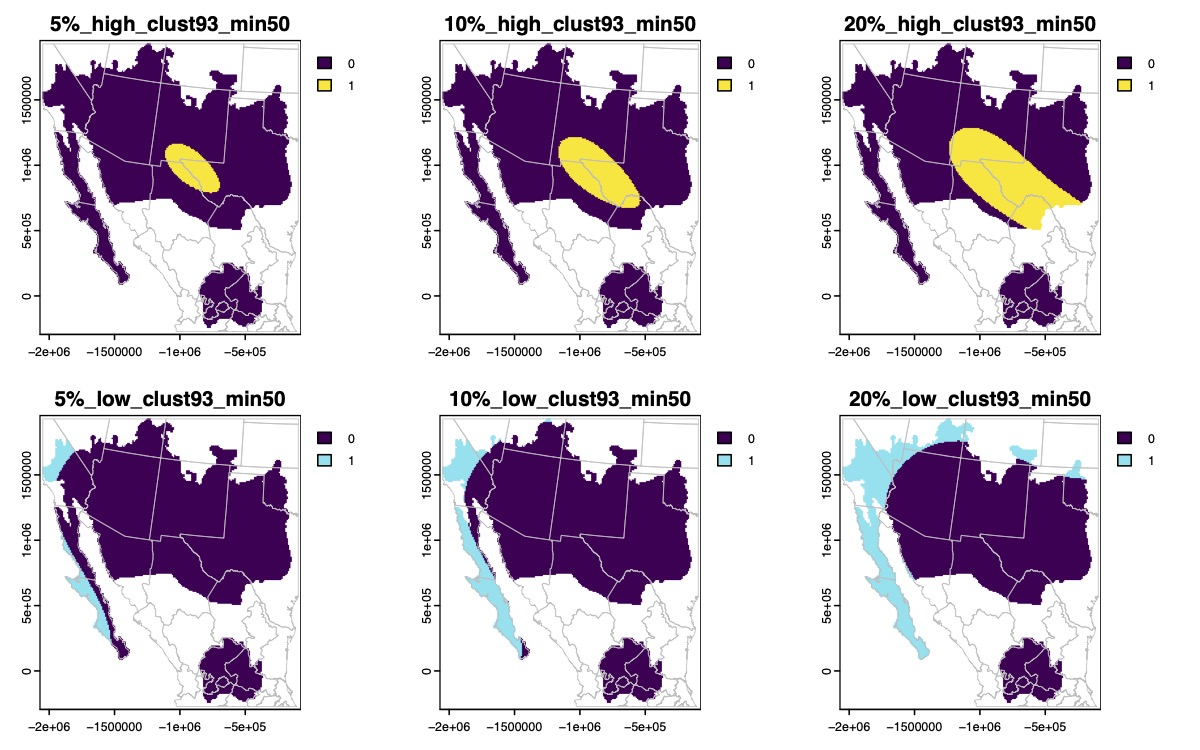

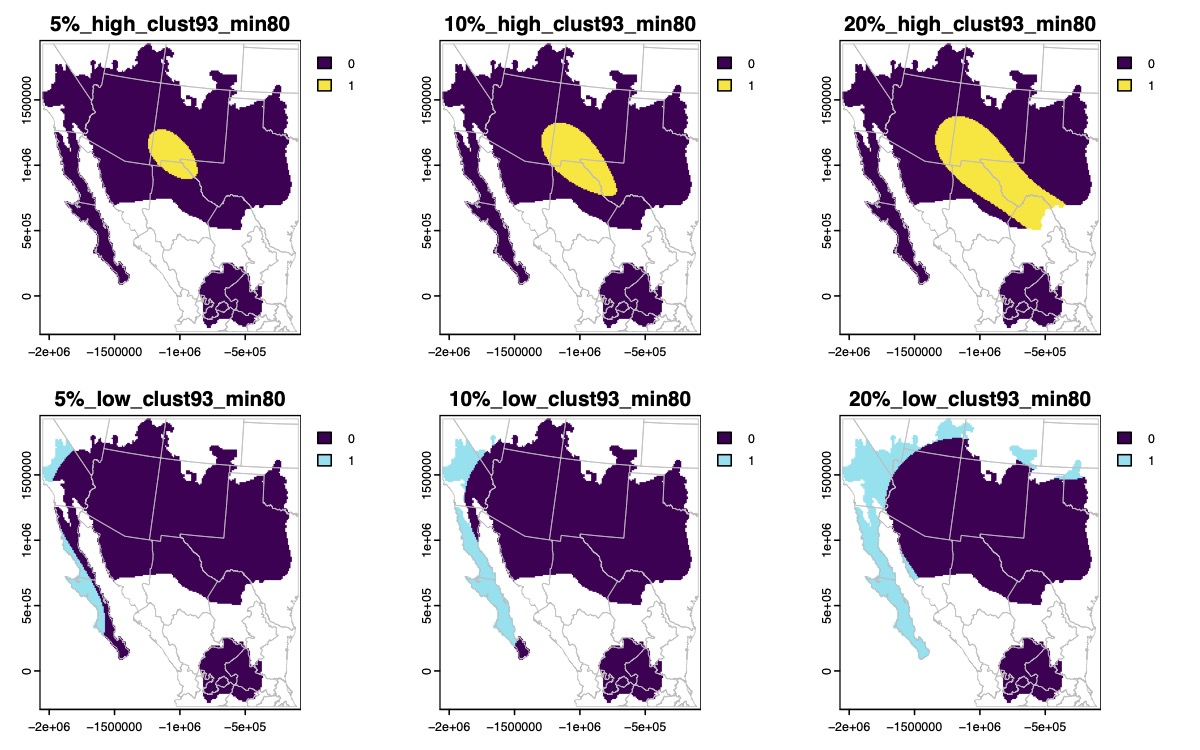

Supplementary Fig. 54.

Binary maps of the highest (yellow) or lowest (blue) genomic diversity for *Anaxyrus punctatus* based on 5%, 10%, or 20% quantile thresholds.

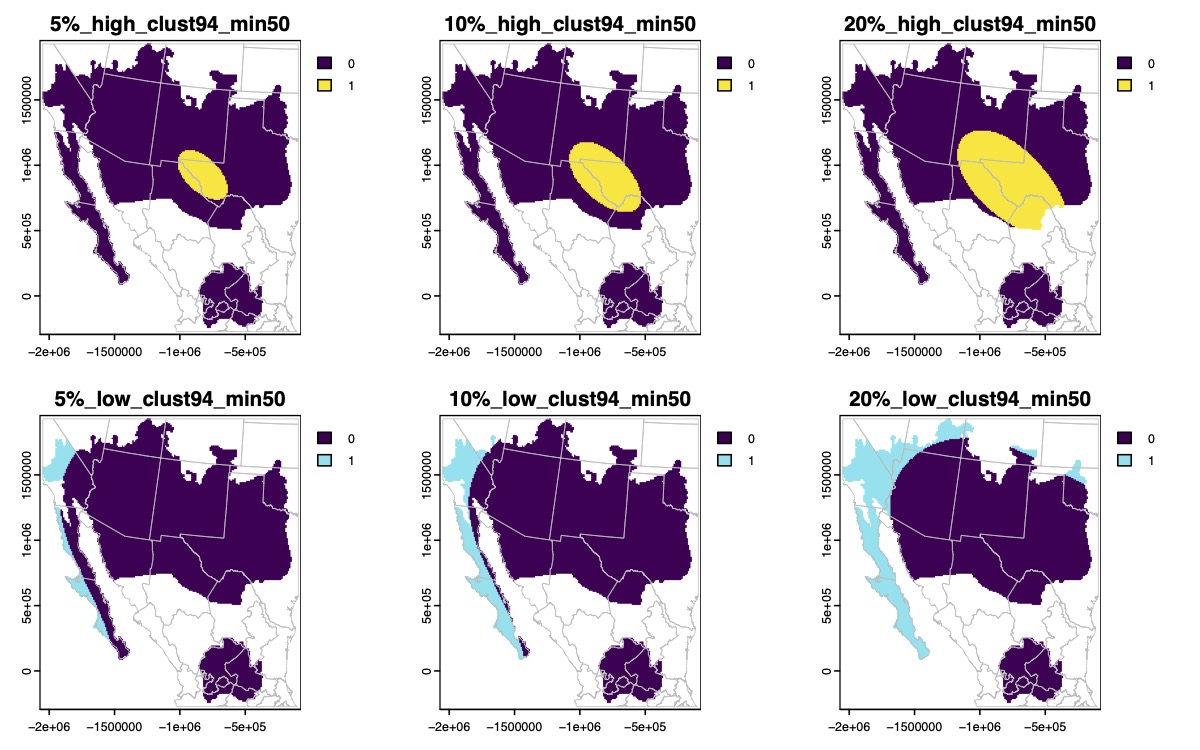

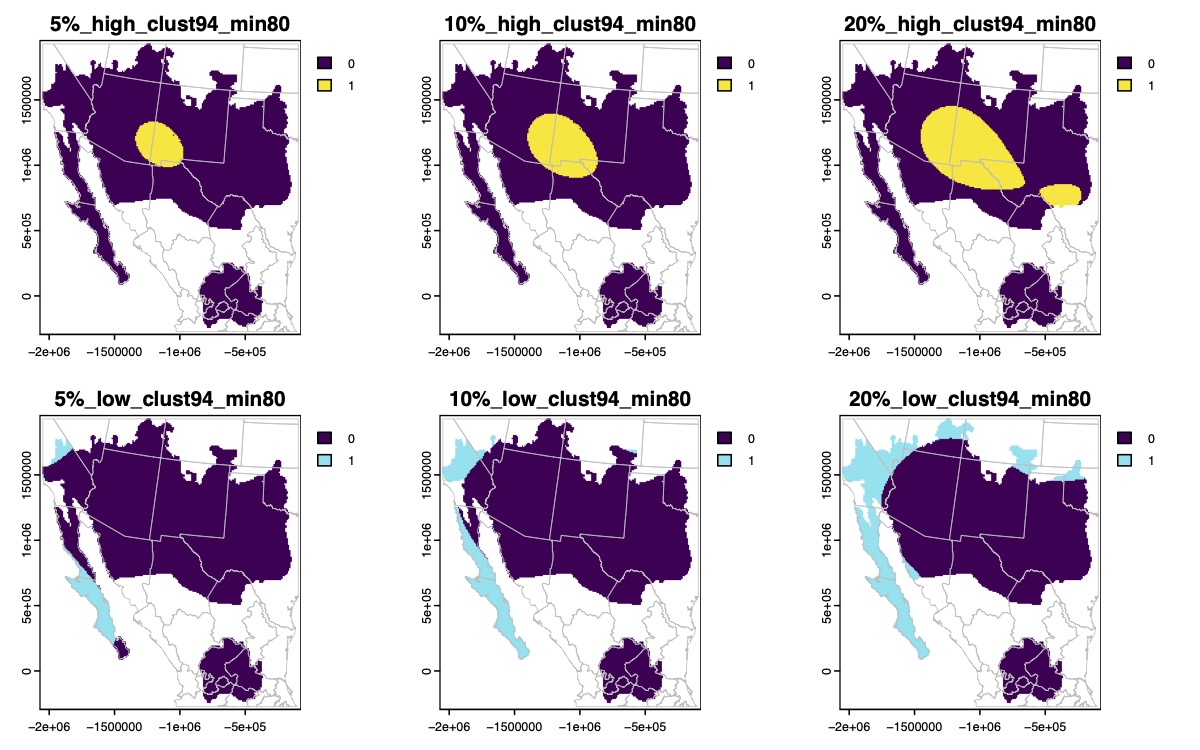

Supplementary Fig. 54. (continued)

Binary maps of the highest (yellow) or lowest (blue) genomic diversity for *Anaxyrus punctatus* based on 5%, 10%, or 20% quantile thresholds.

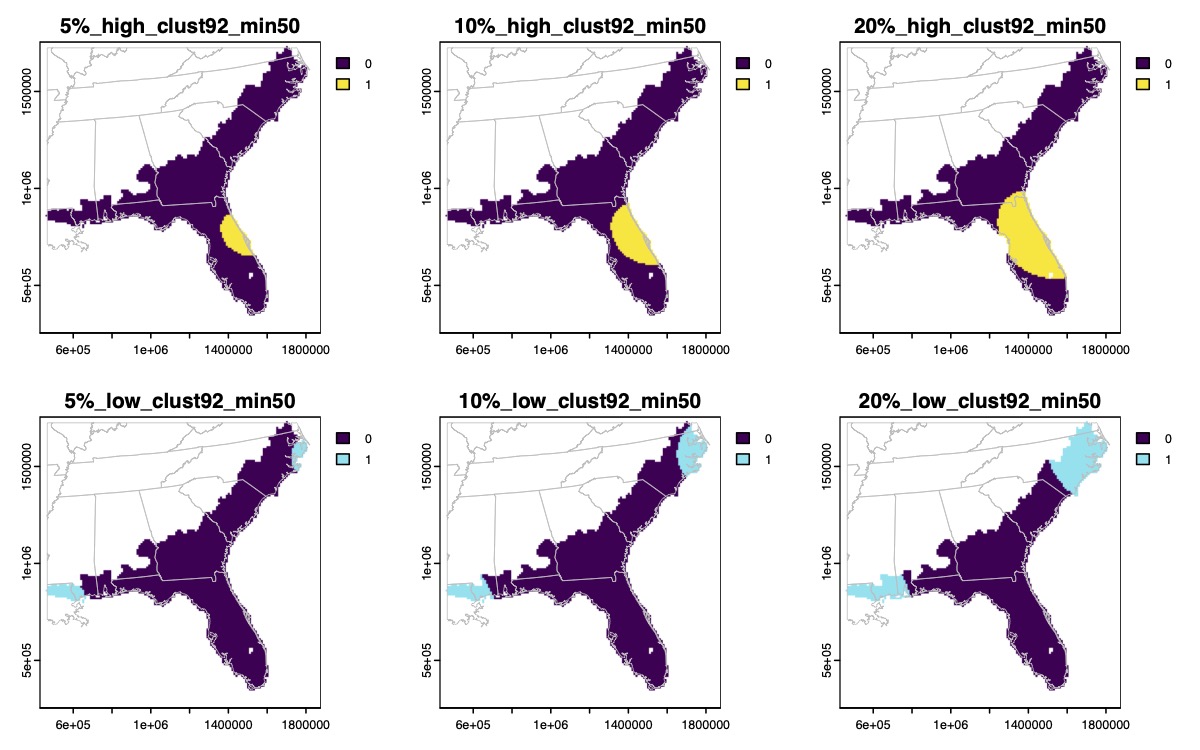

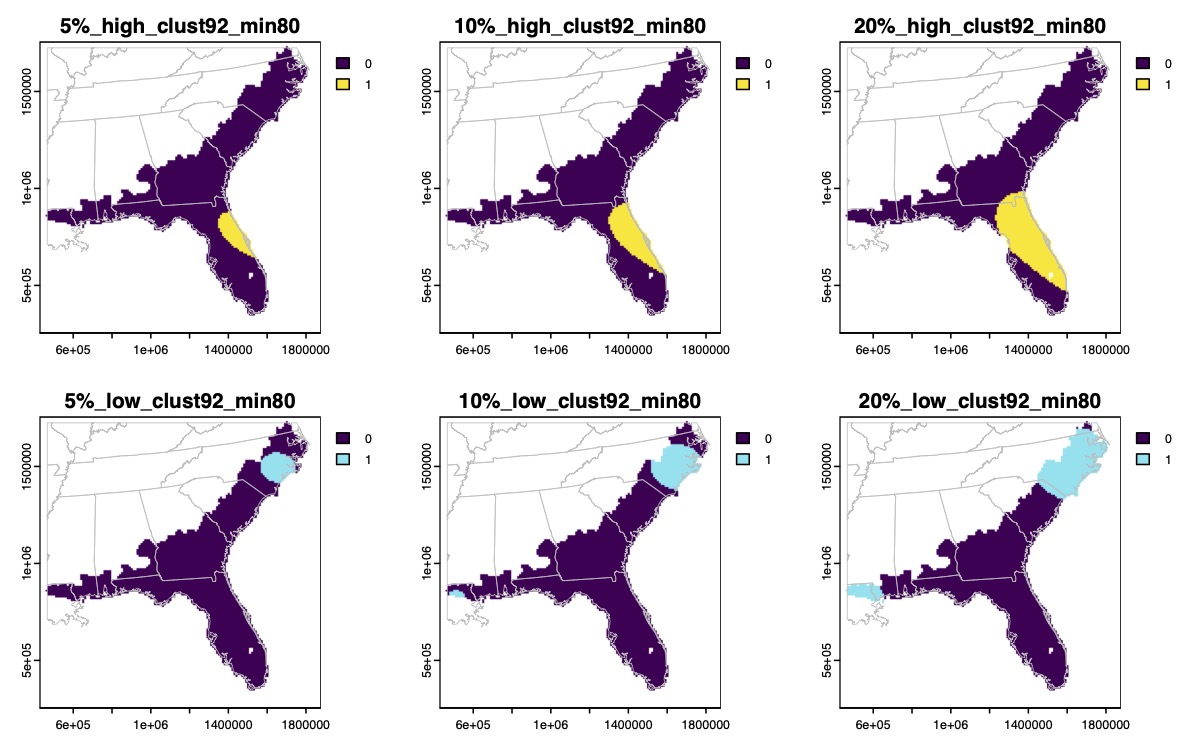

Supplementary Fig. 55.

Binary maps of the highest (yellow) or lowest (blue) genomic diversity for *Anaxyrus quercicus* based on 5%, 10%, or 20% quantile thresholds.

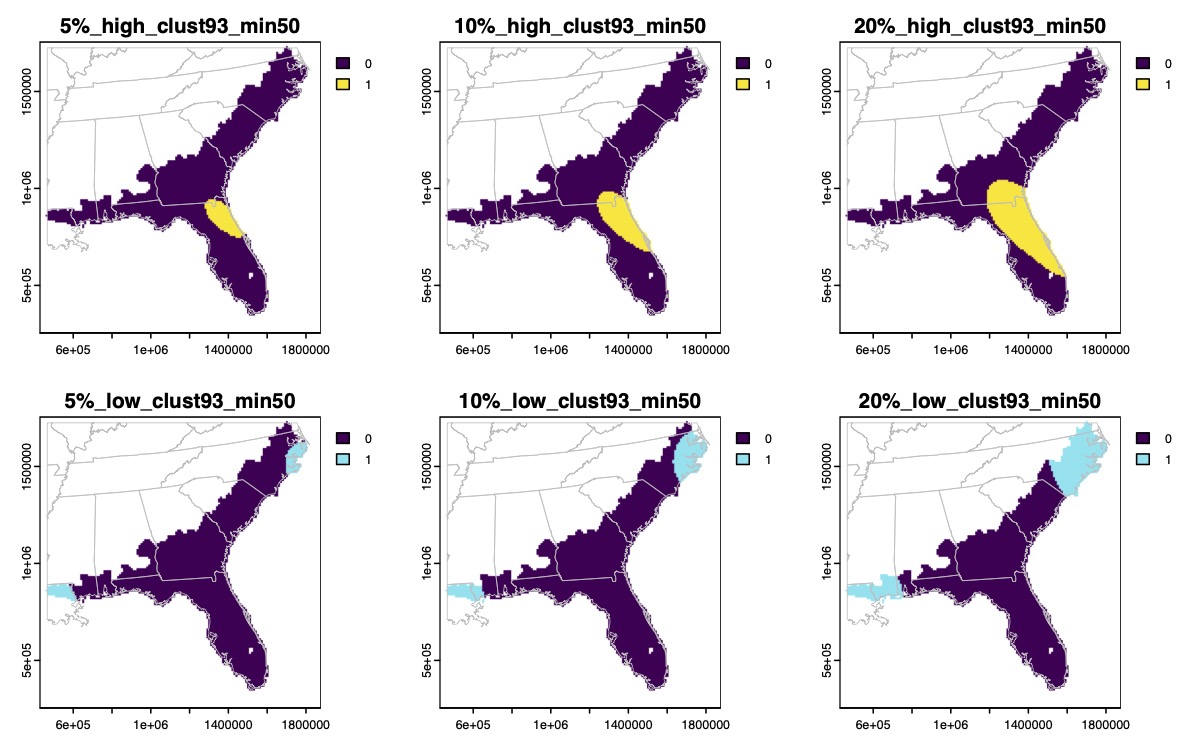

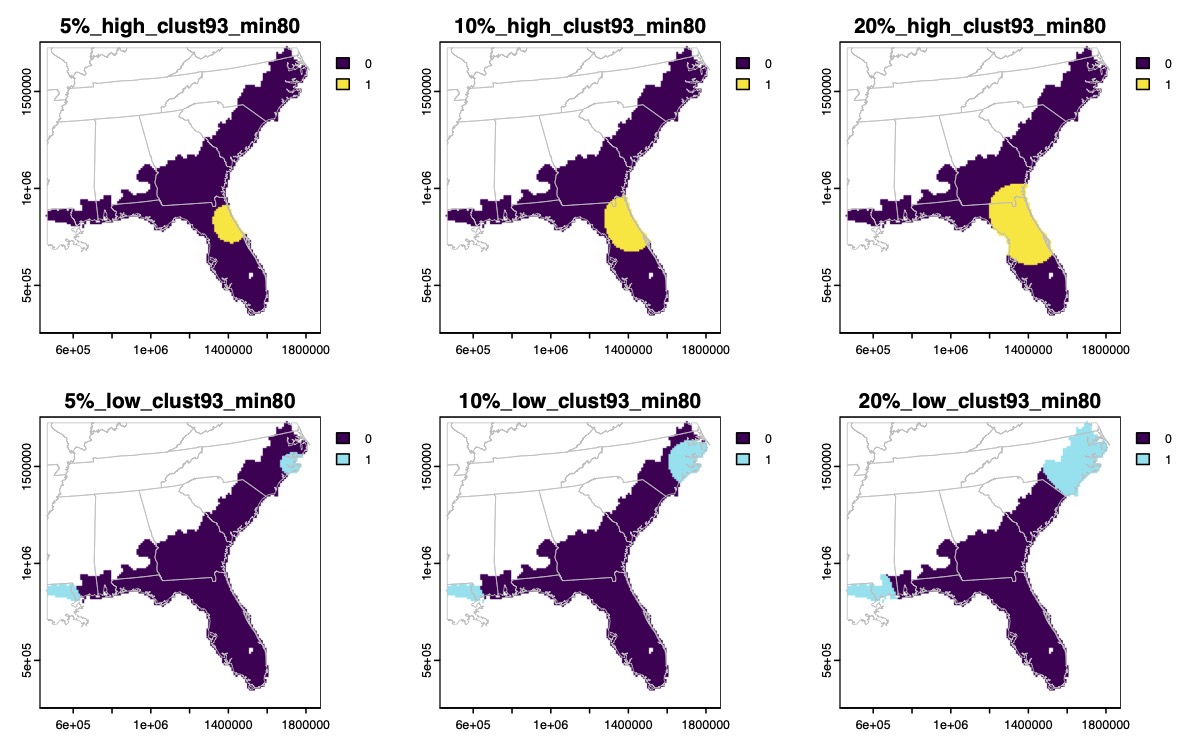

Supplementary Fig. 55. (continued)

Binary maps of the highest (yellow) or lowest (blue) genomic diversity for *Anaxyrus quercicus* based on 5%, 10%, or 20% quantile thresholds.

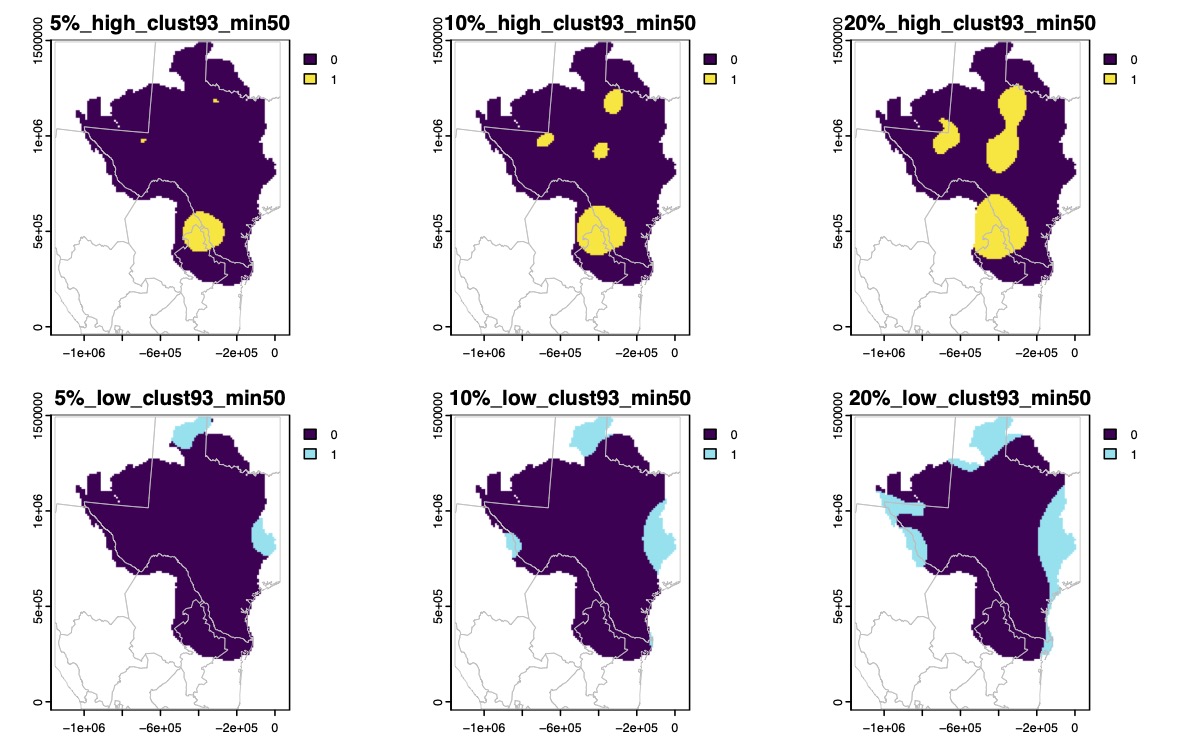

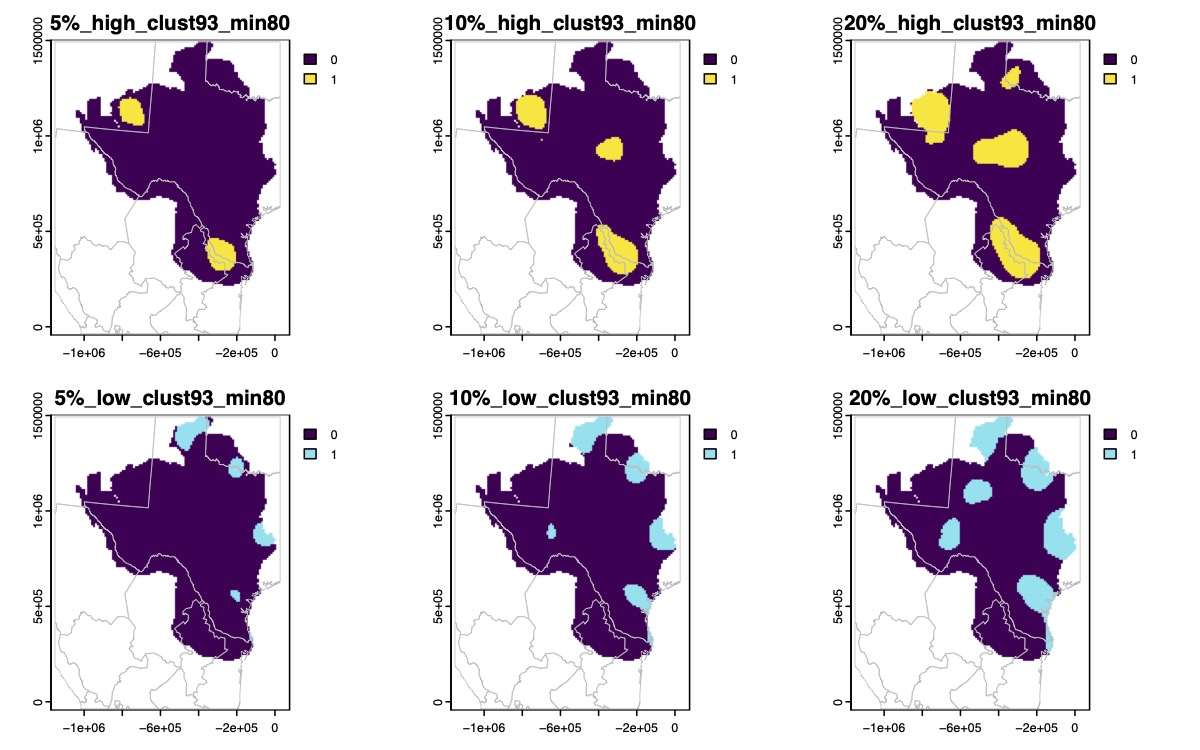

Supplementary Fig. 56.

Binary maps of the highest (yellow) or lowest (blue) genomic diversity for *Anaxyrus speciosus* based on 5%, 10%, or 20% quantile thresholds.

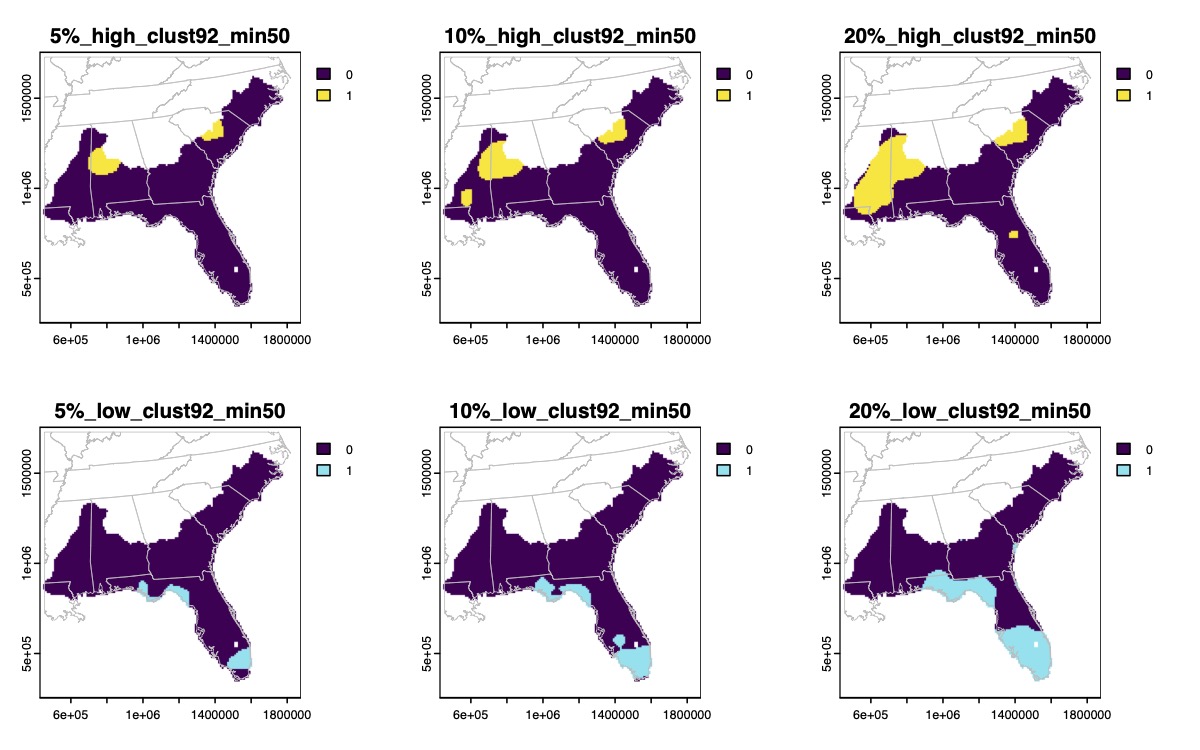

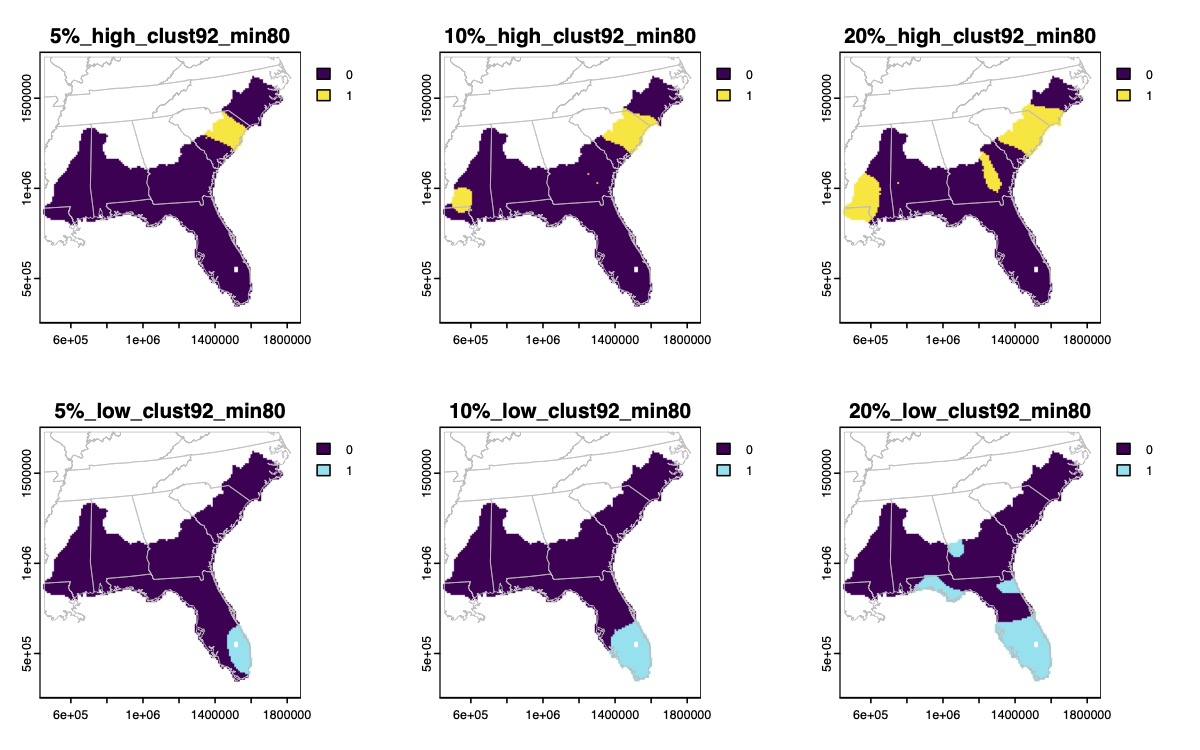

Supplementary Fig. 57.

Binary maps of the highest (yellow) or lowest (blue) genomic diversity for *Anaxyrus terrestris* based on 5%, 10%, or 20% quantile thresholds.

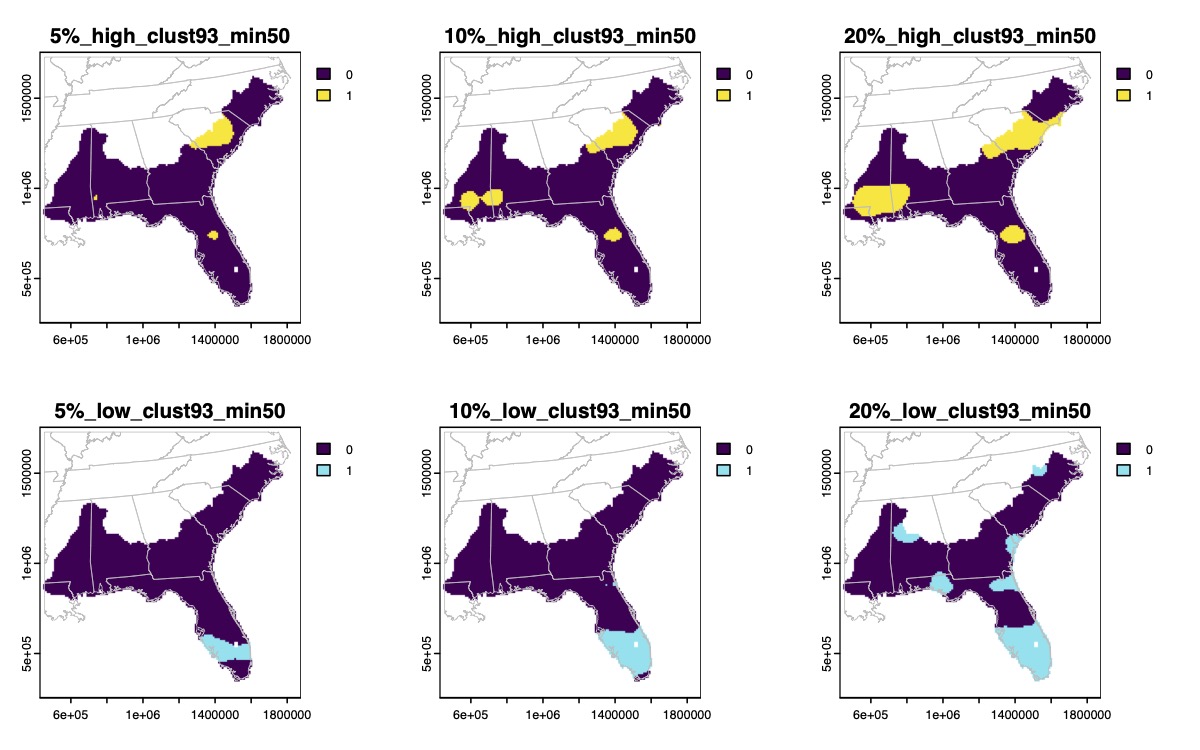

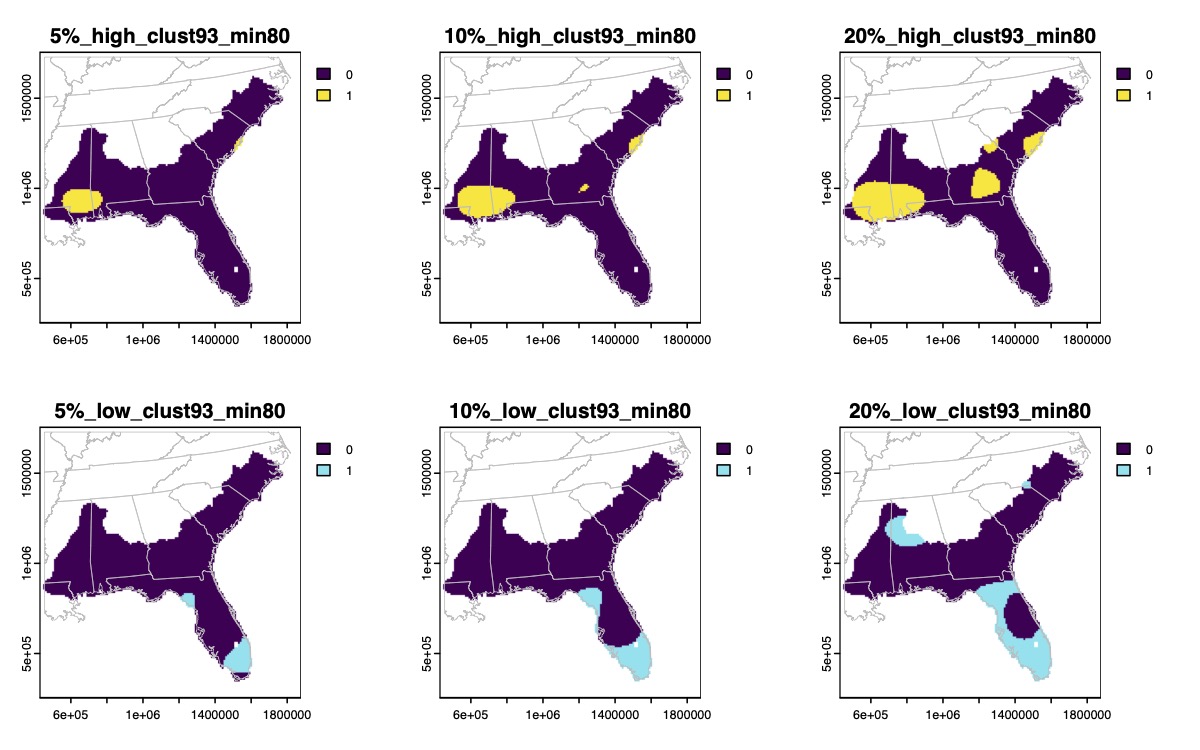

Supplementary Fig. 57. (continued)

Binary maps of the highest (yellow) or lowest (blue) genomic diversity for *Anaxyrus terrestris* based on 5%, 10%, or 20% quantile thresholds.

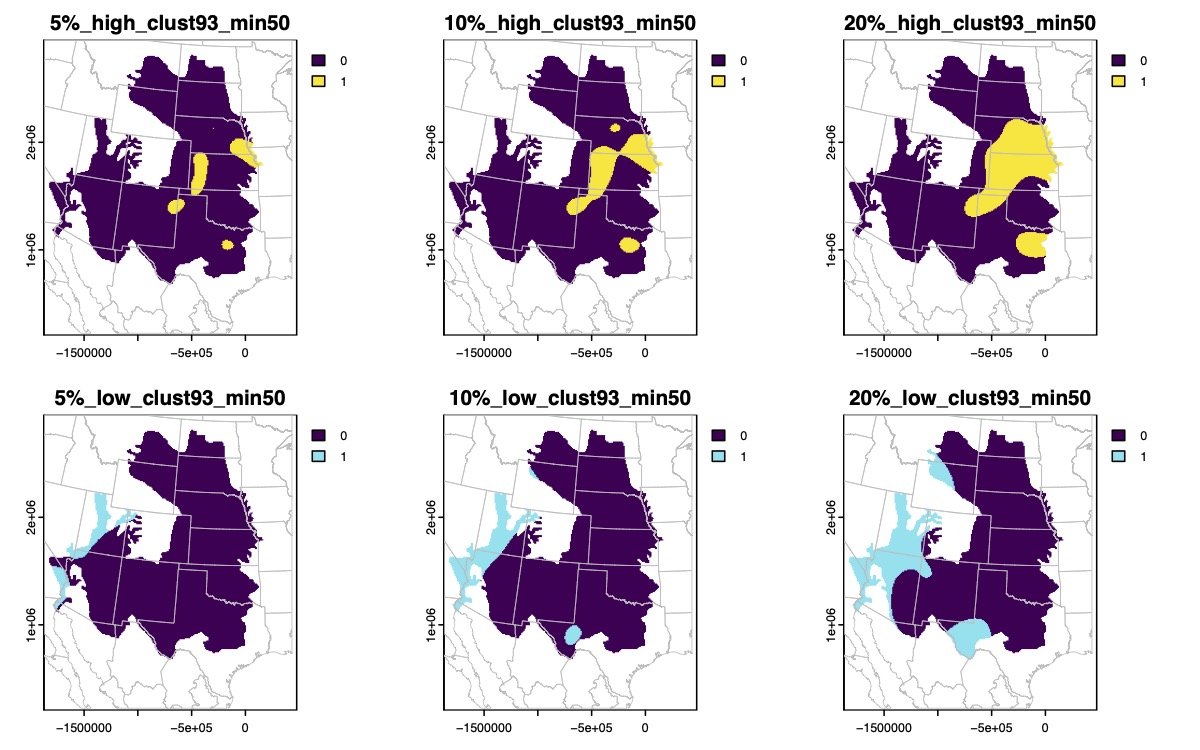

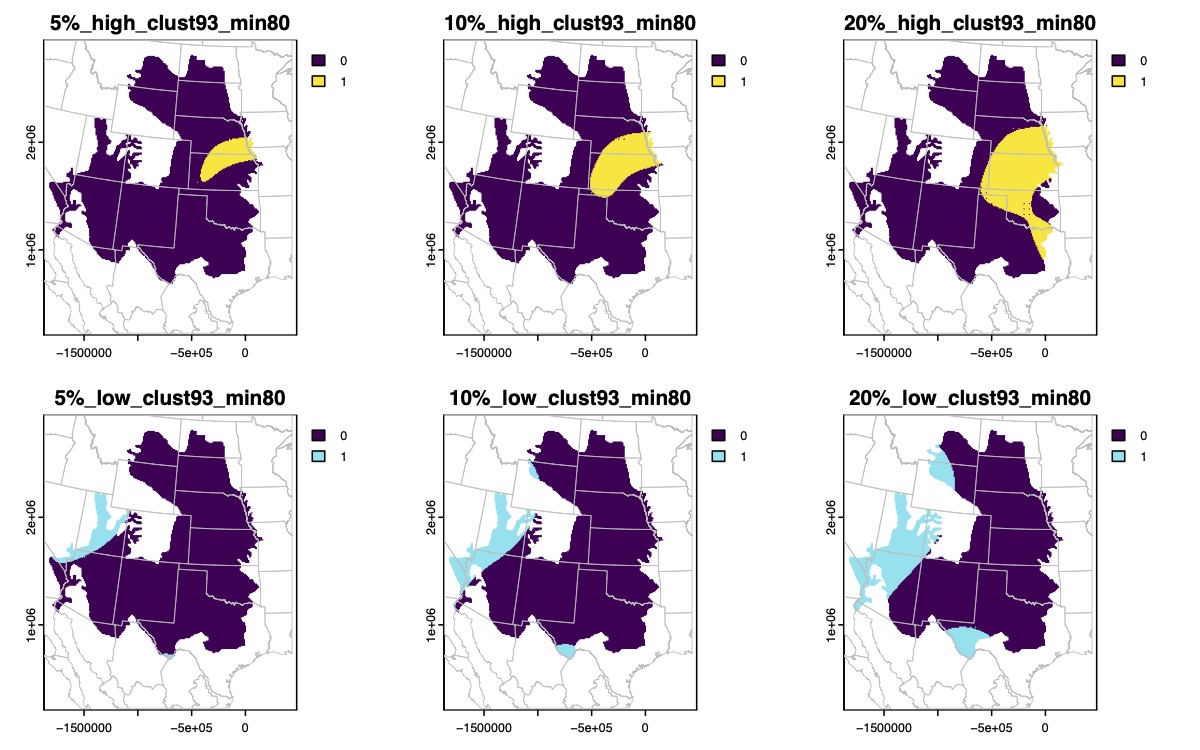

Supplementary Fig. 58.

Binary maps of the highest (yellow) or lowest (blue) genomic diversity for *Anaxyrus woodhousii* based on 5%, 10%, or 20% quantile thresholds.

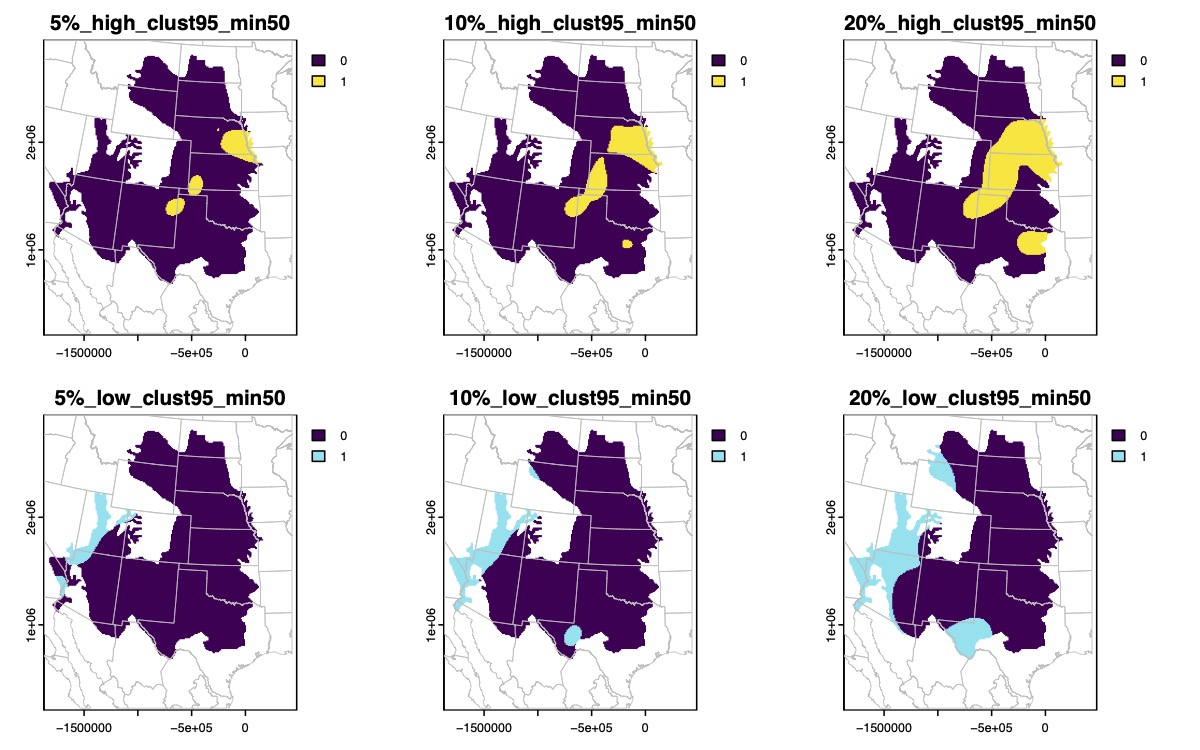

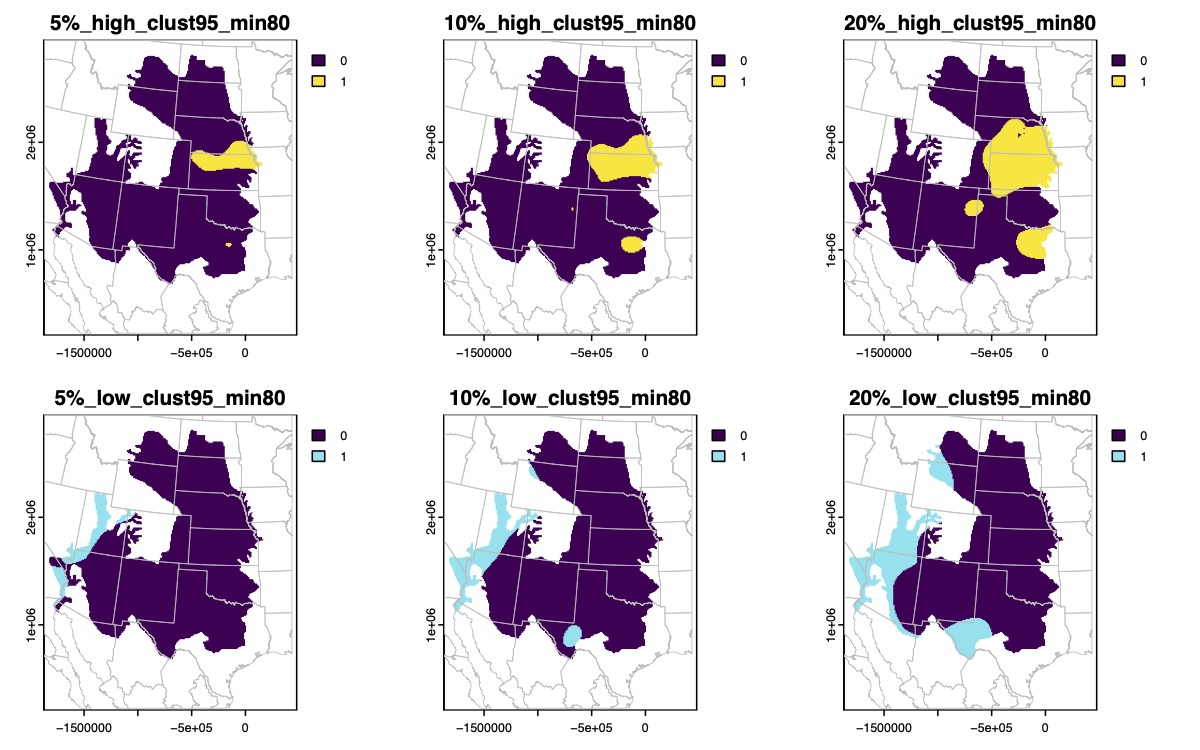

Supplementary Fig. 58. (continued)

Binary maps of the highest (yellow) or lowest (blue) genomic diversity for *Anaxyrus woodhousii* based on 5%, 10%, or 20% quantile thresholds.

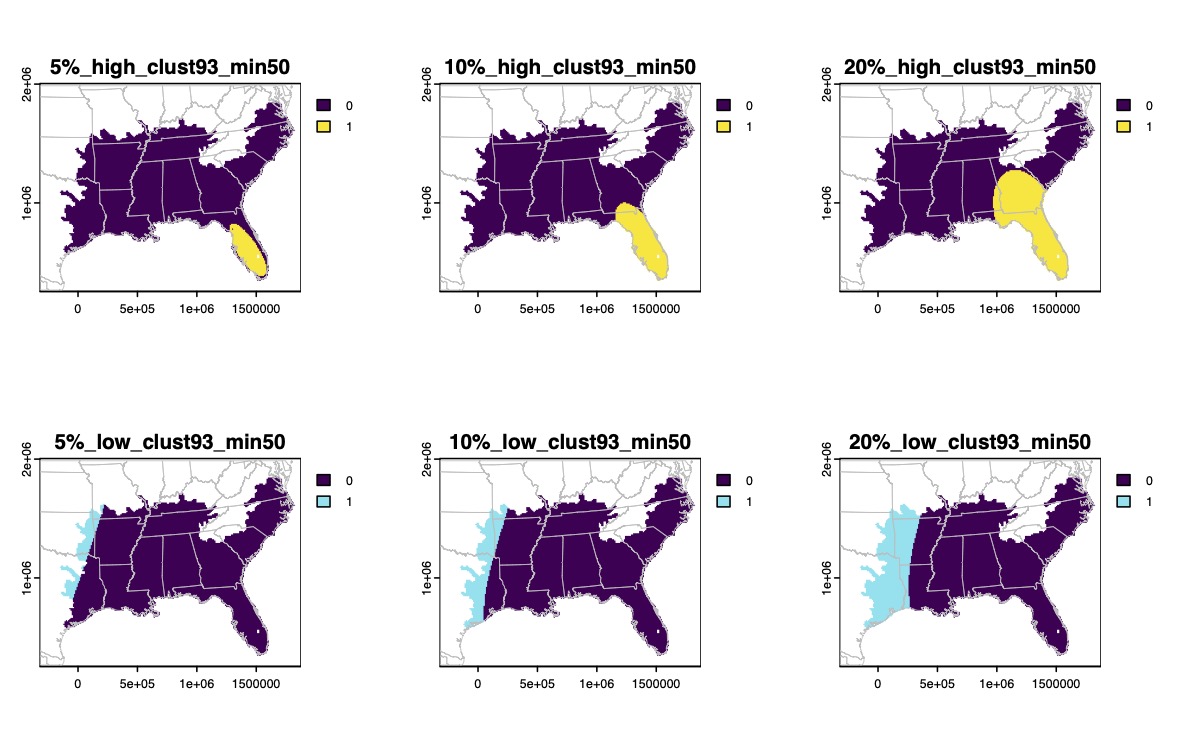

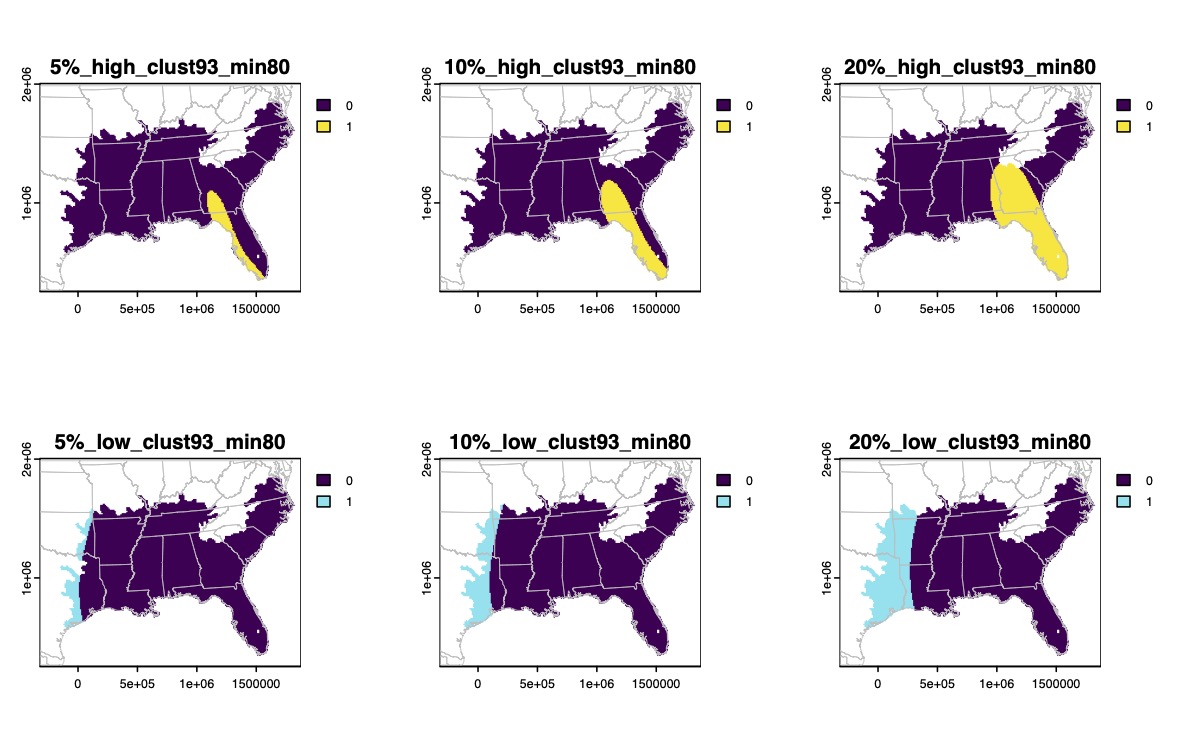

Supplementary Fig. 59.

Binary maps of the highest (yellow) or lowest (blue) genomic diversity for *Gastrophryne carolinensis* based on 5%, 10%, or 20% quantile thresholds.

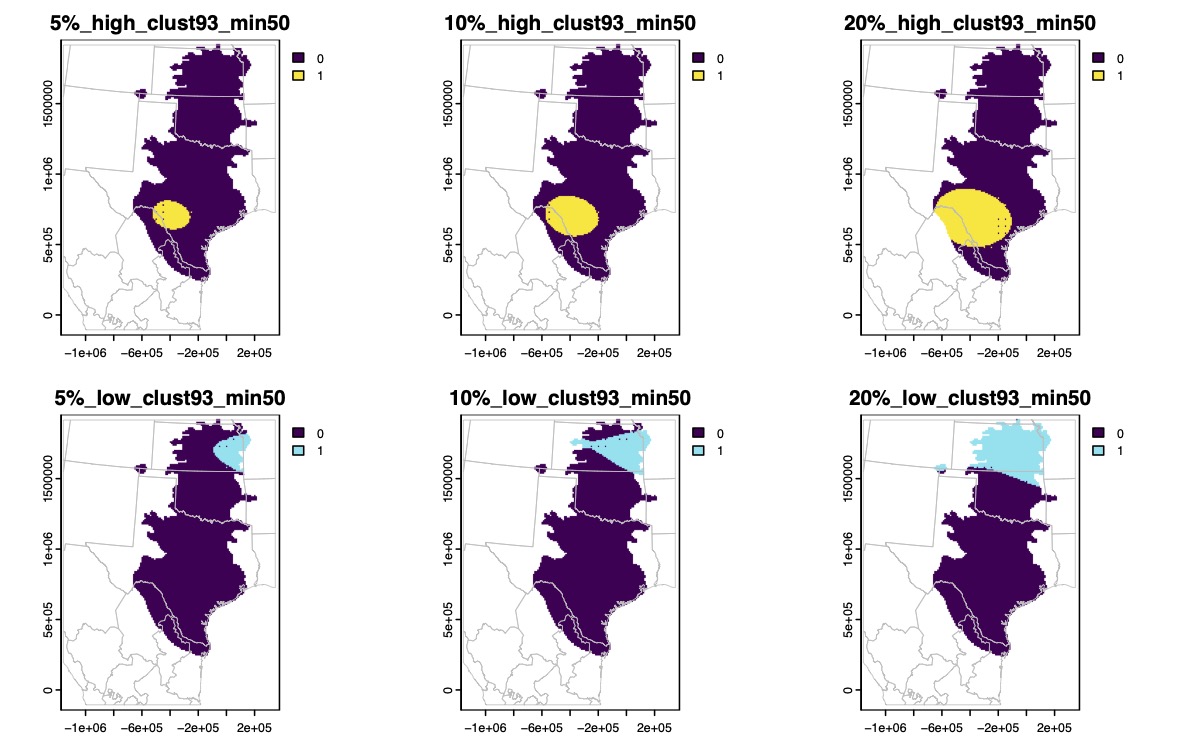

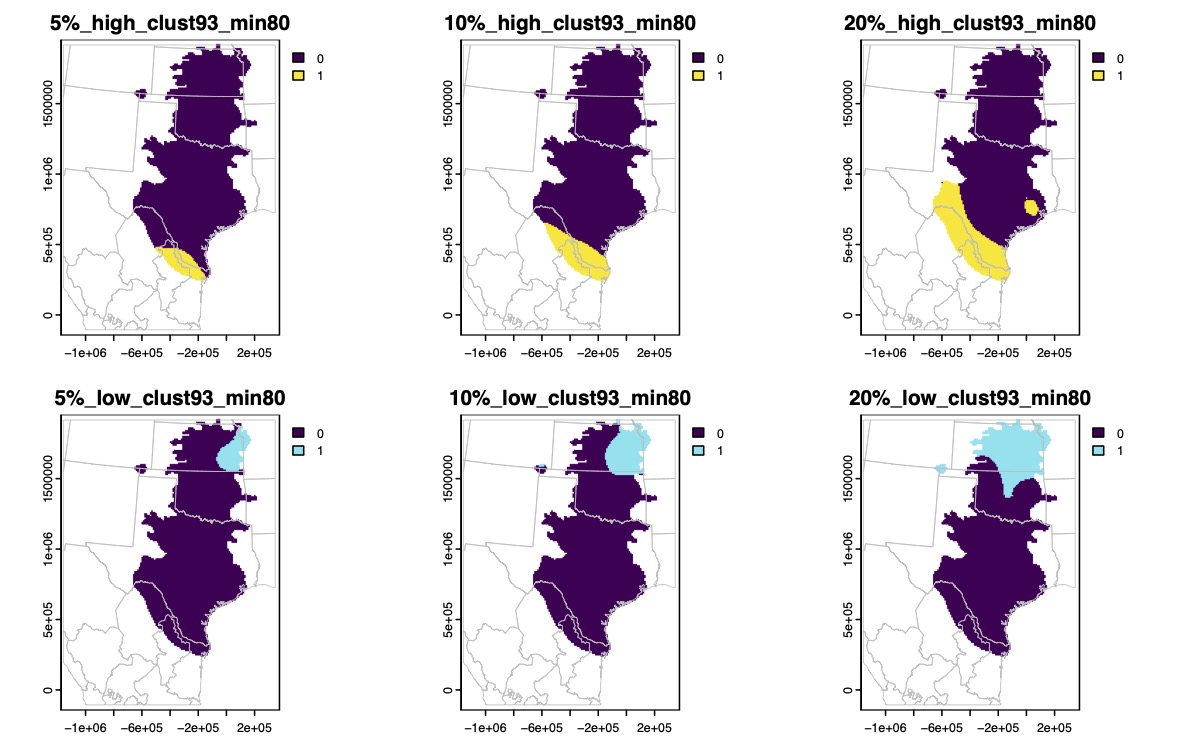

Supplementary Fig. 60.

Binary maps of the highest (yellow) or lowest (blue) genomic diversity for *Gastrophryne olivacea* based on 5%, 10%, or 20% quantile thresholds.

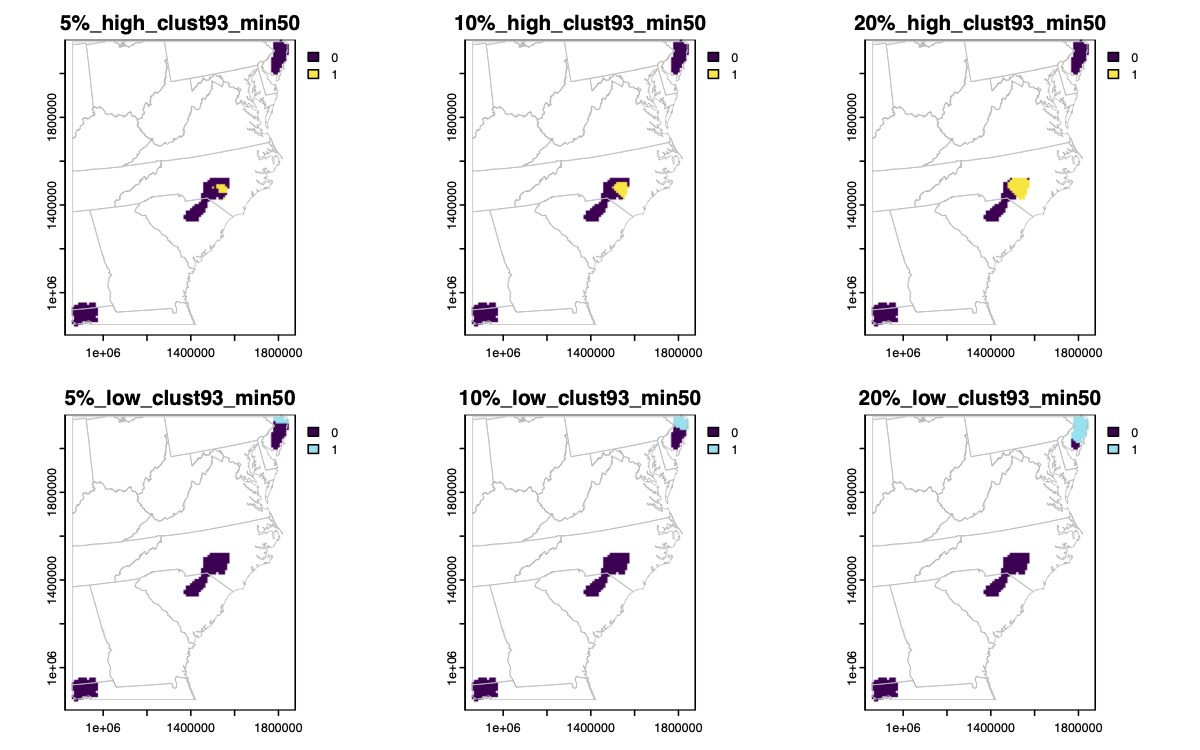

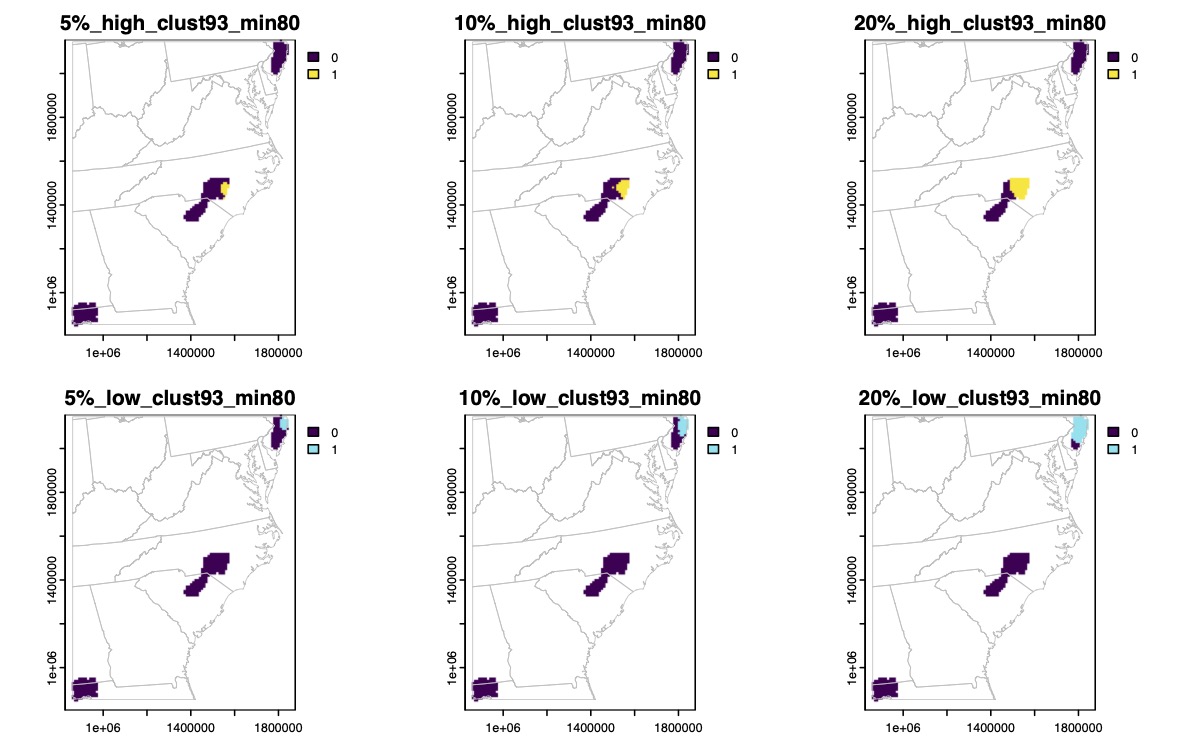

Supplementary Fig. 61.

Binary maps of the highest (yellow) or lowest (blue) genomic diversity for *Hyla andersonii* based on 5%, 10%, or 20% quantile thresholds.

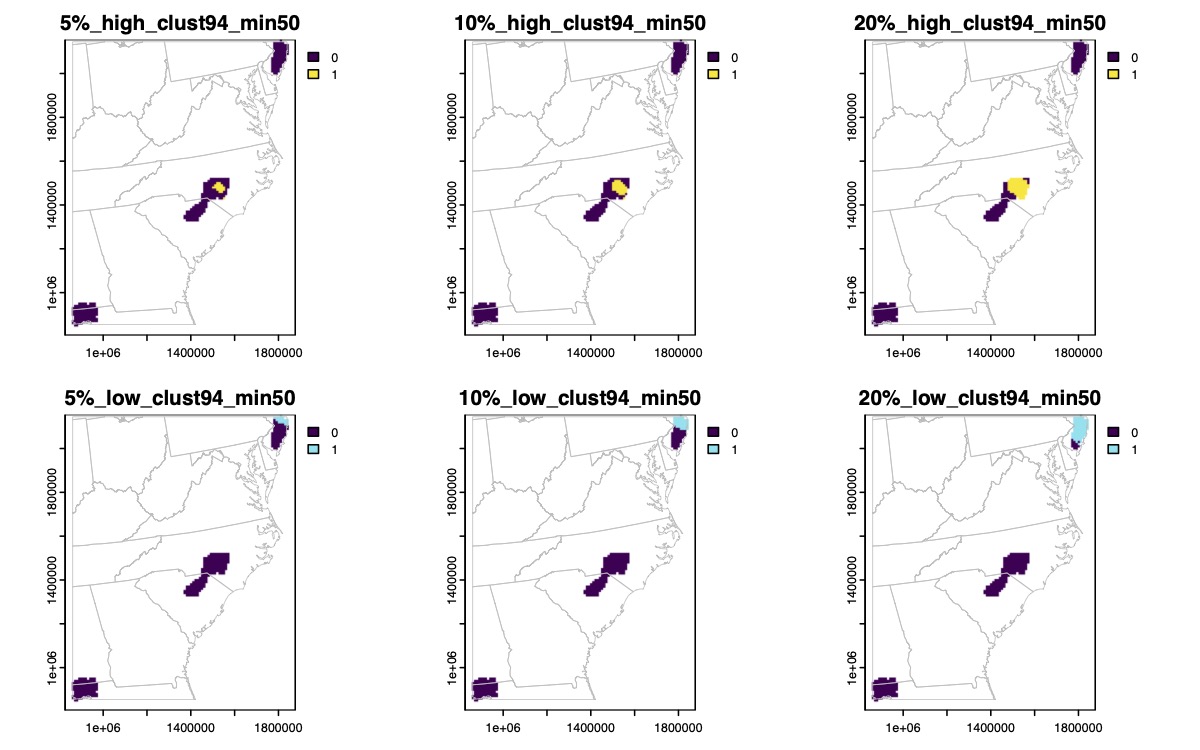

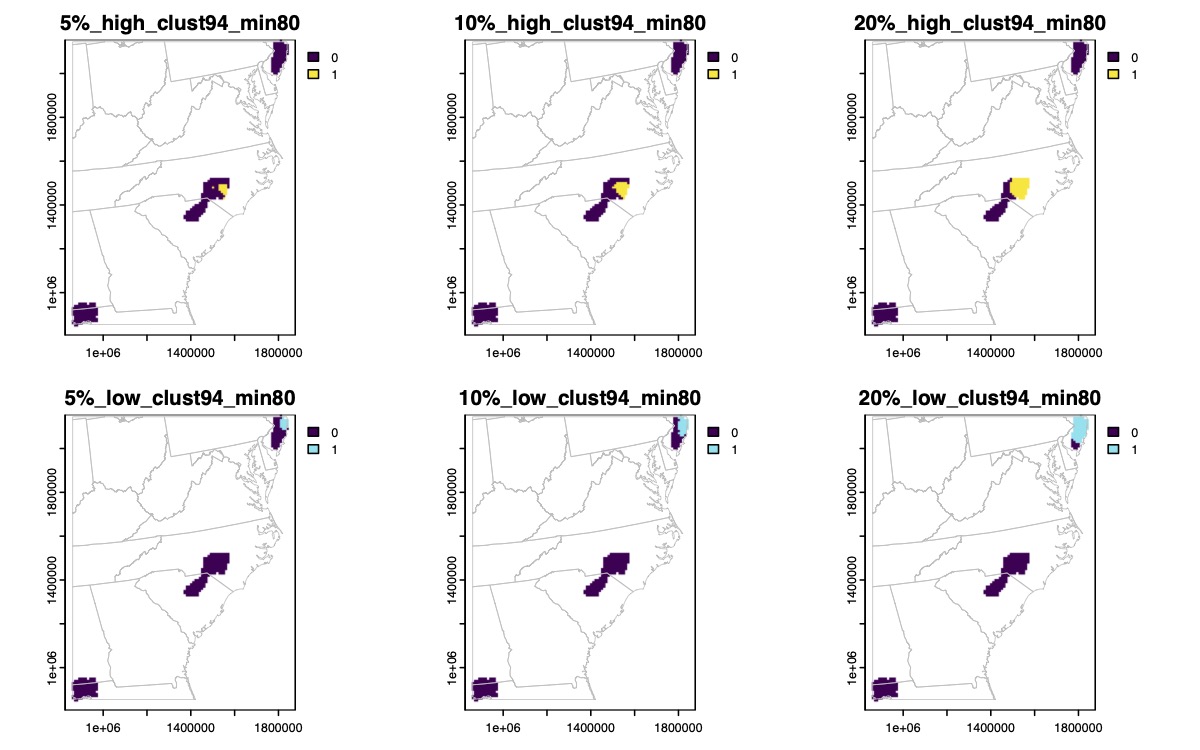

Supplementary Fig. 61. (continued)

Binary maps of the highest (yellow) or lowest (blue) genomic diversity for *Hyla andersonii* based on 5%, 10%, or 20% quantile thresholds.

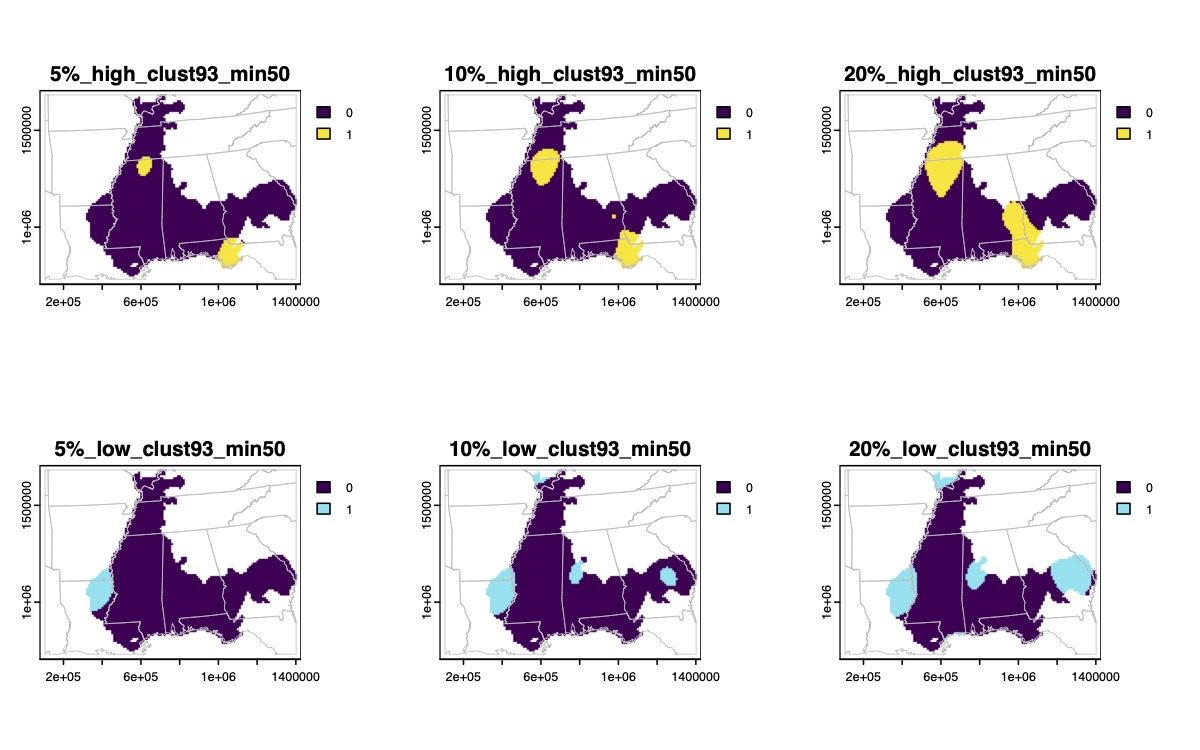

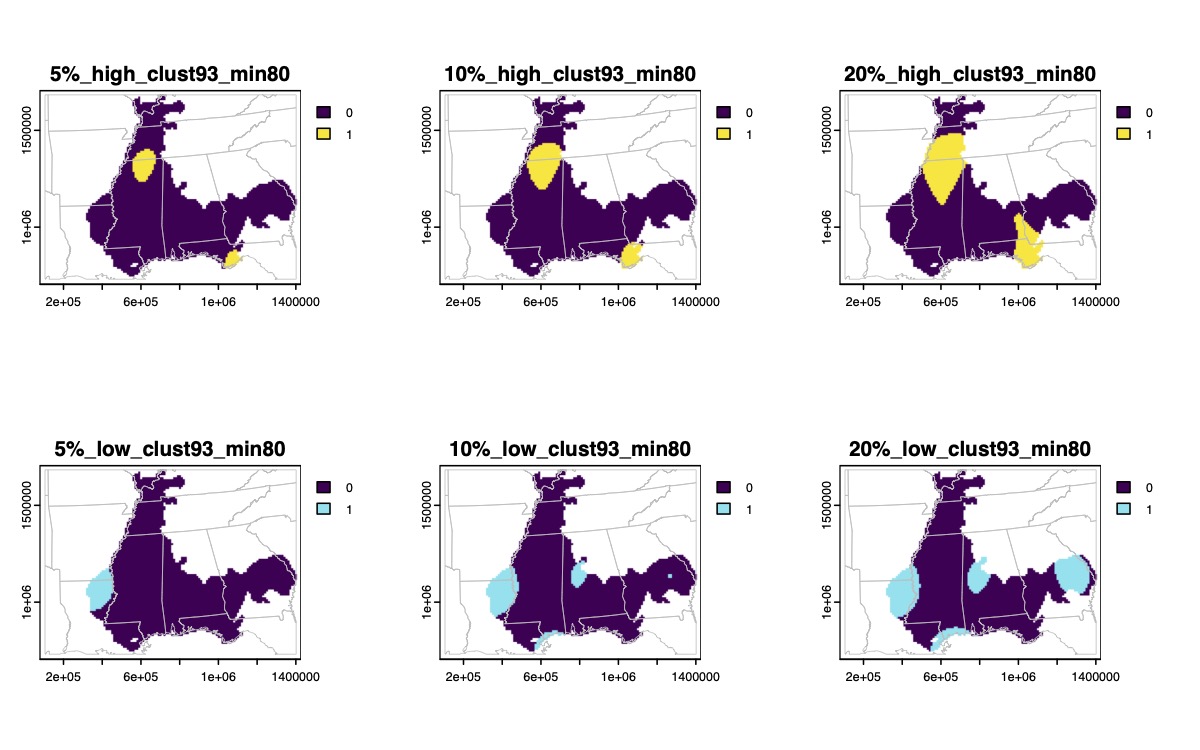

Supplementary Fig. 62.

Binary maps of the highest (yellow) or lowest (blue) genomic diversity for *Hyla avivoca* based on 5%, 10%, or 20% quantile thresholds.

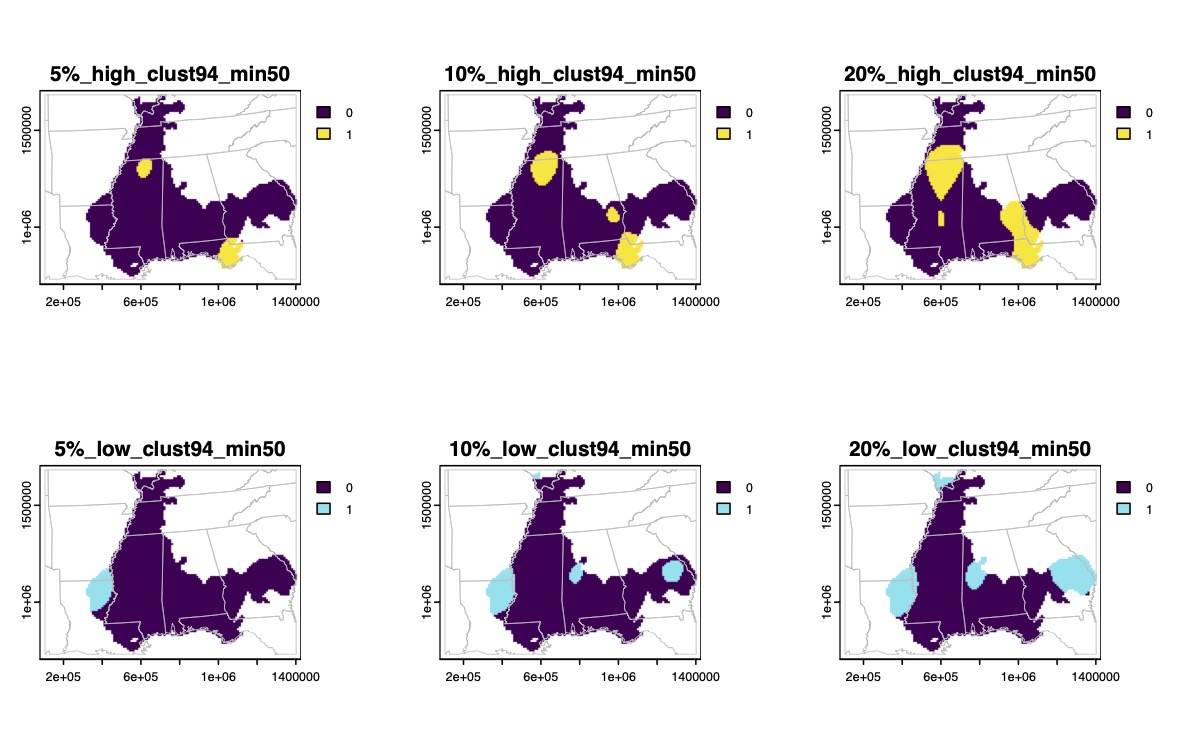

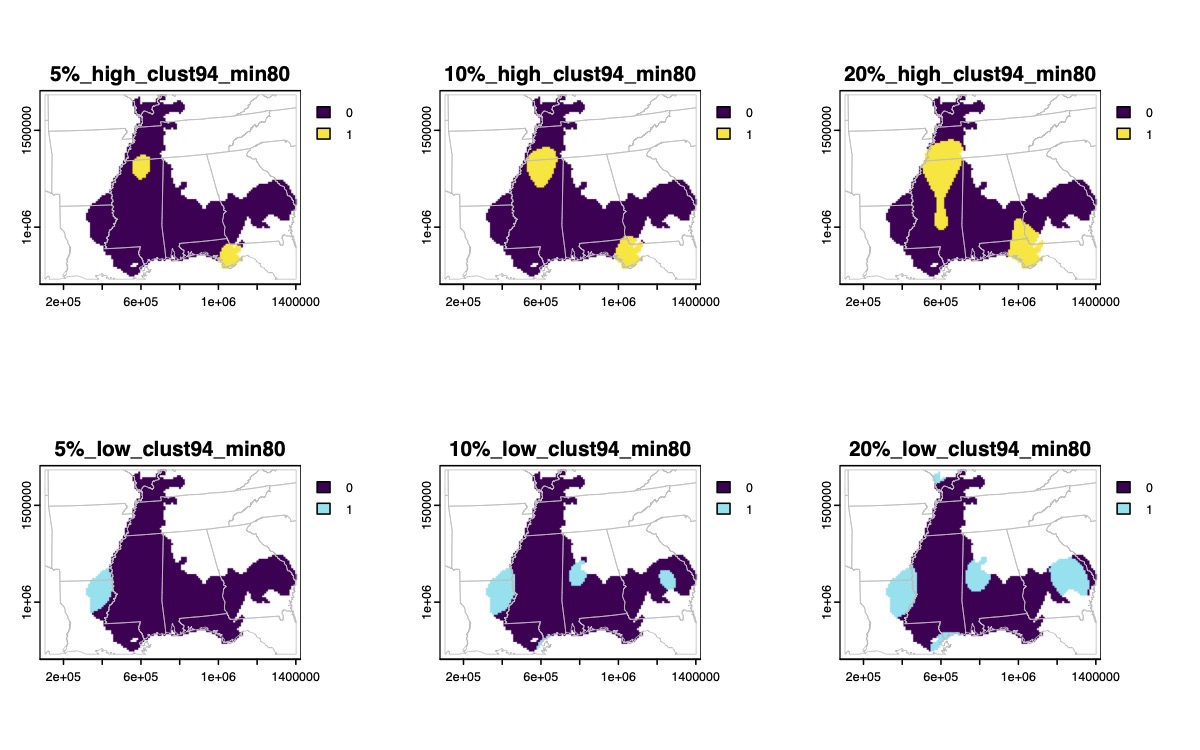

Supplementary Fig. 62. (continued)

Binary maps of the highest (yellow) or lowest (blue) genomic diversity for *Hyla avivoca* based on 5%, 10%, or 20% quantile thresholds.

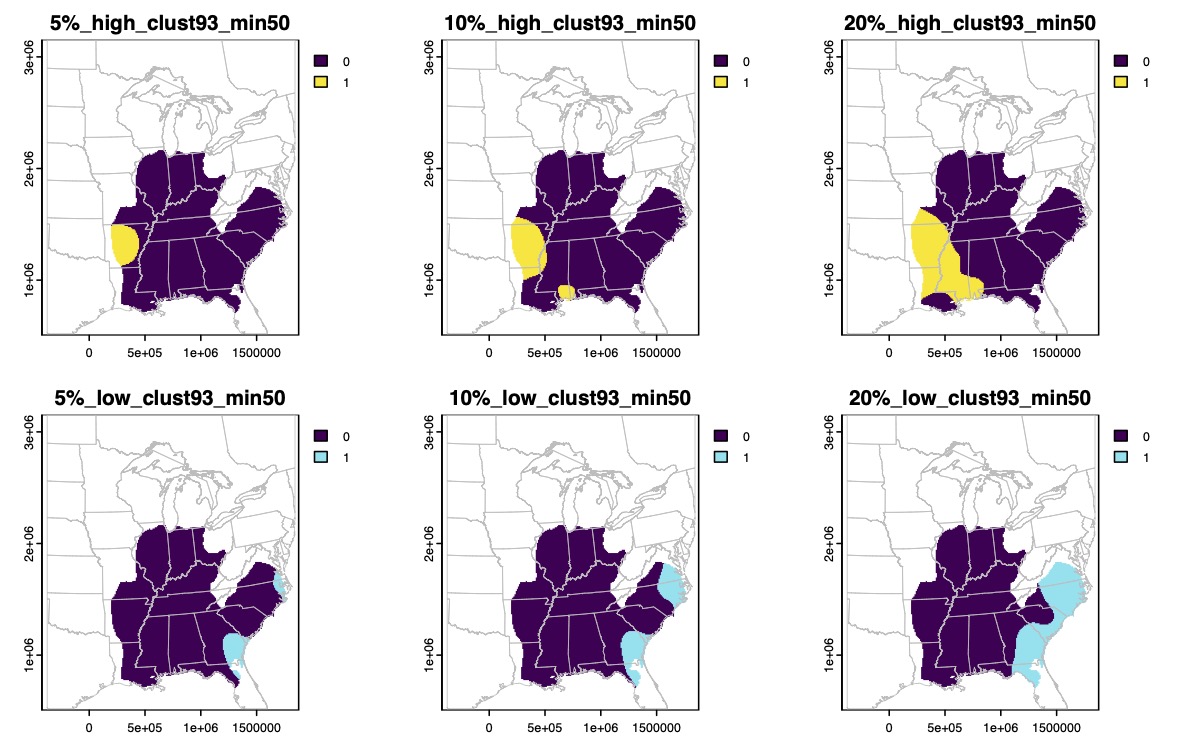

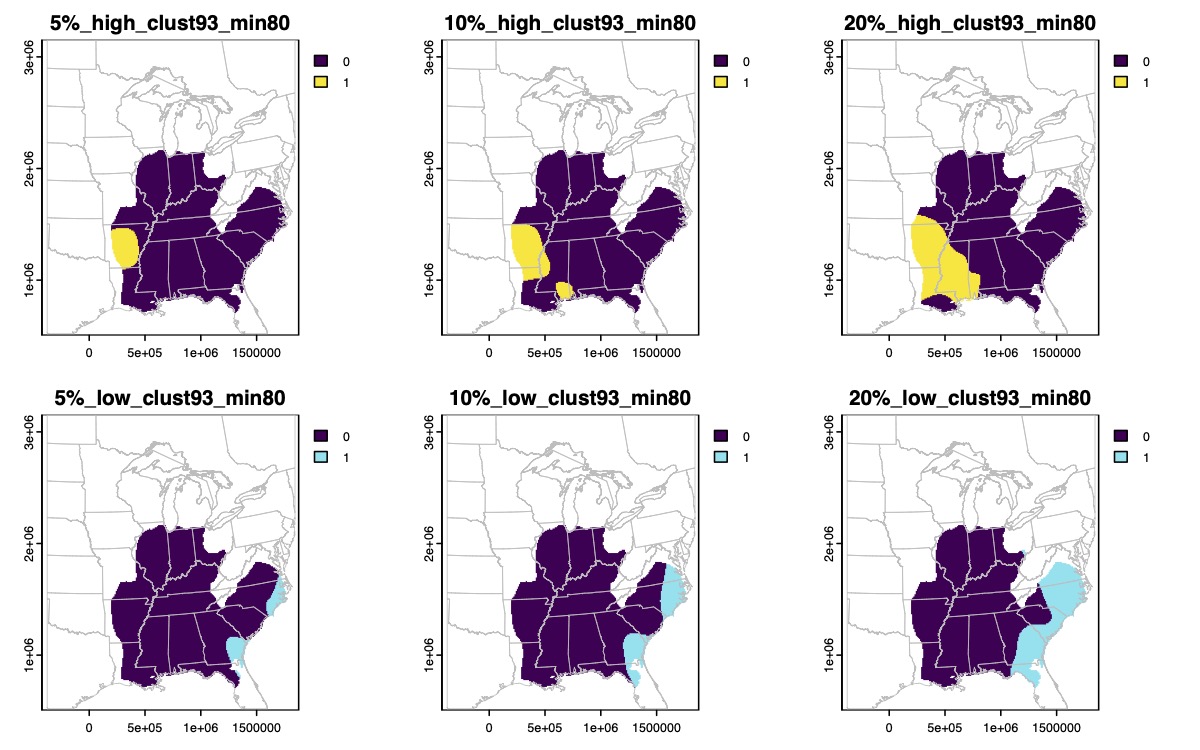

Supplementary Fig. 63.

Binary maps of the highest (yellow) or lowest (blue) genomic diversity for *Hyla chrysoscelis* based on 5%, 10%, or 20% quantile thresholds.

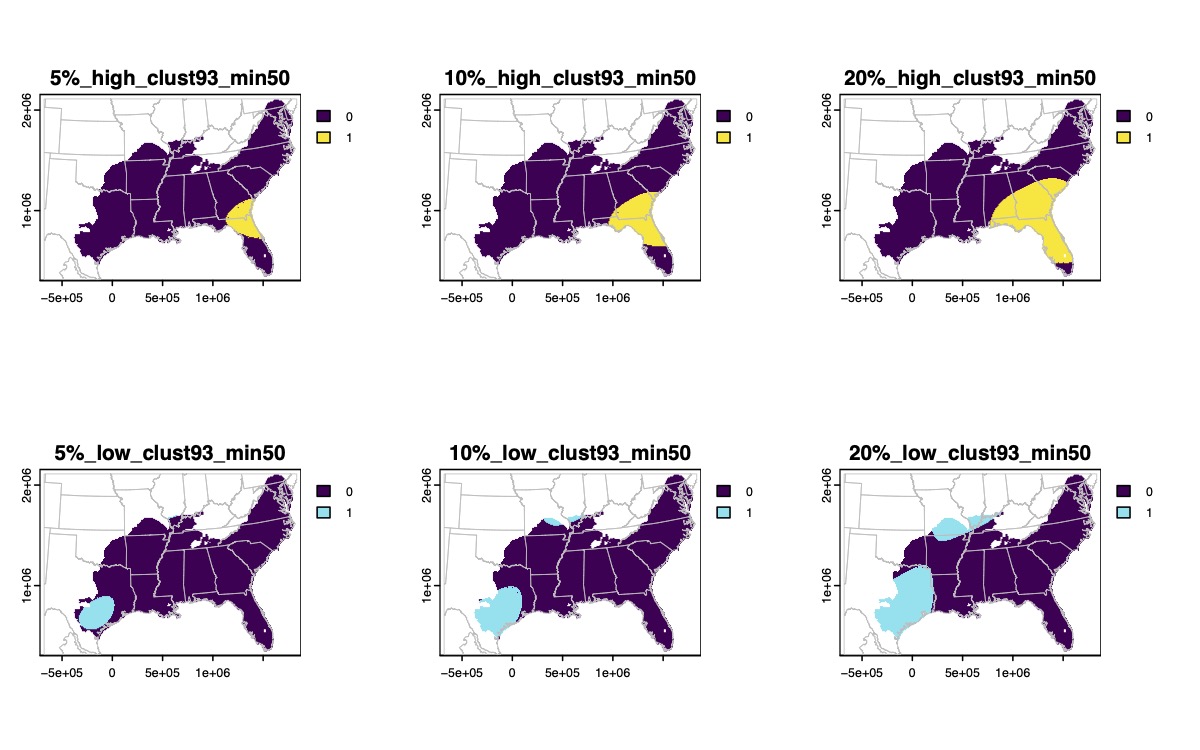

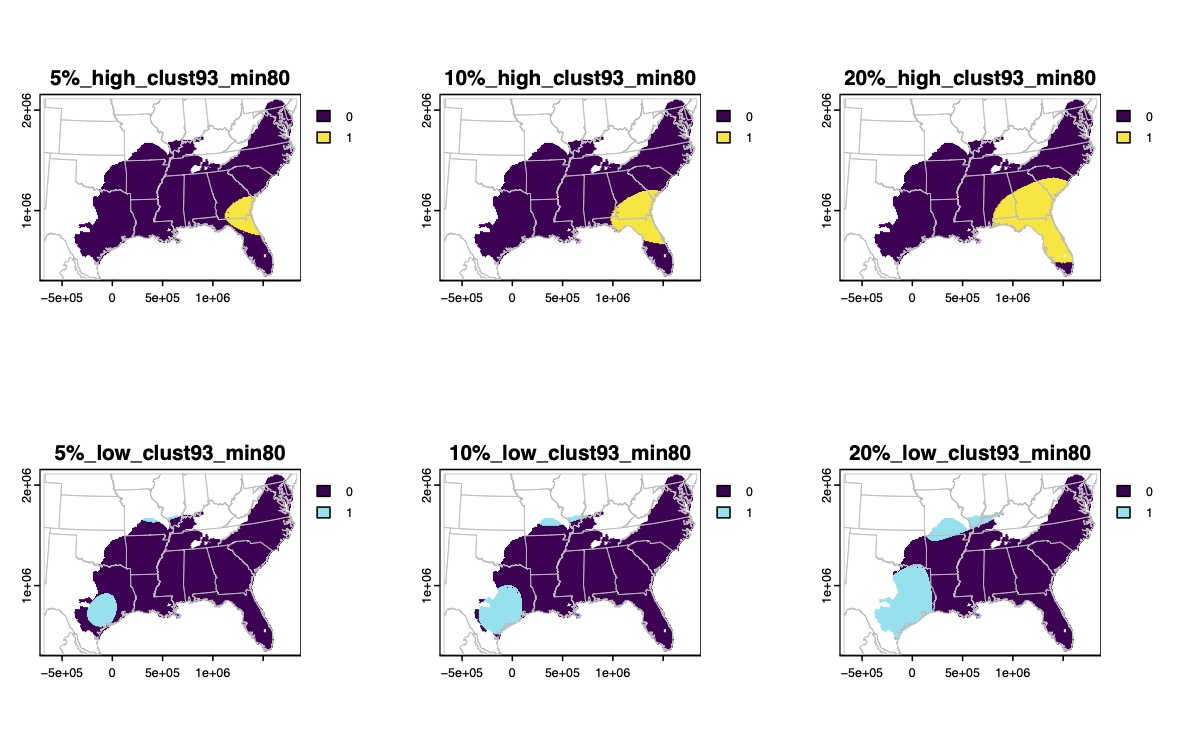

Supplementary Fig. 64.

Binary maps of the highest (yellow) or lowest (blue) genomic diversity for *Hyla cinerea* based on 5%, 10%, or 20% quantile thresholds.

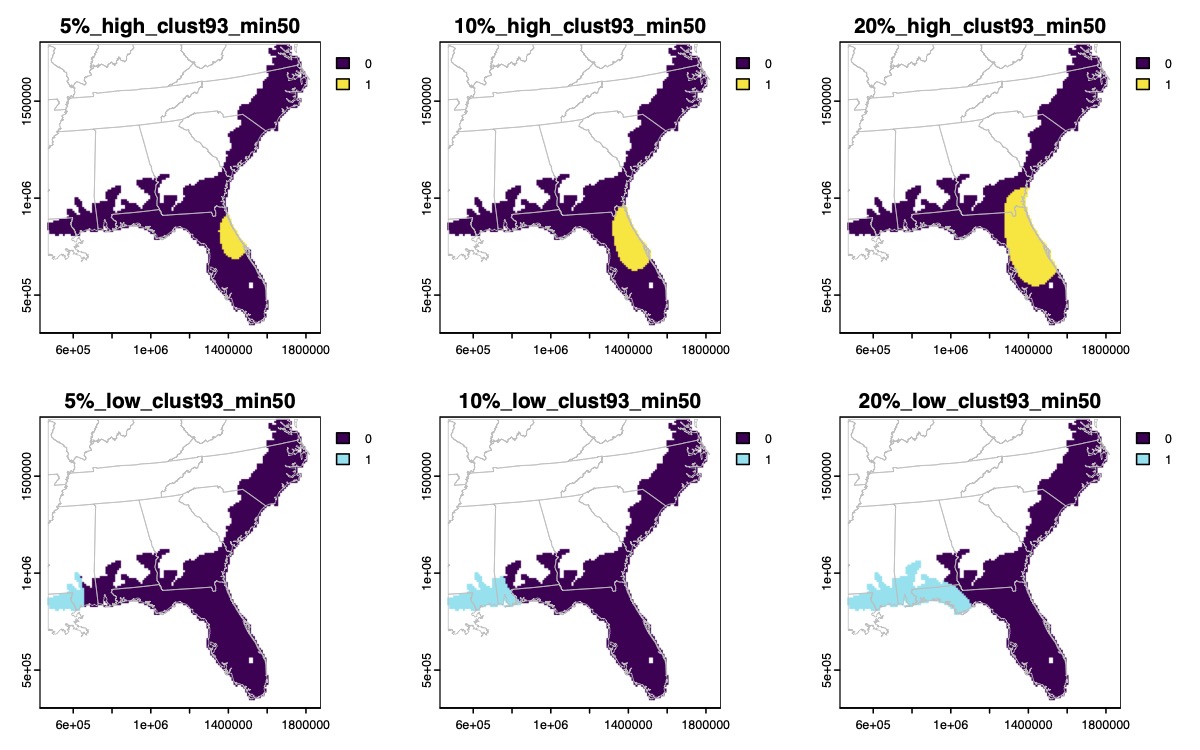

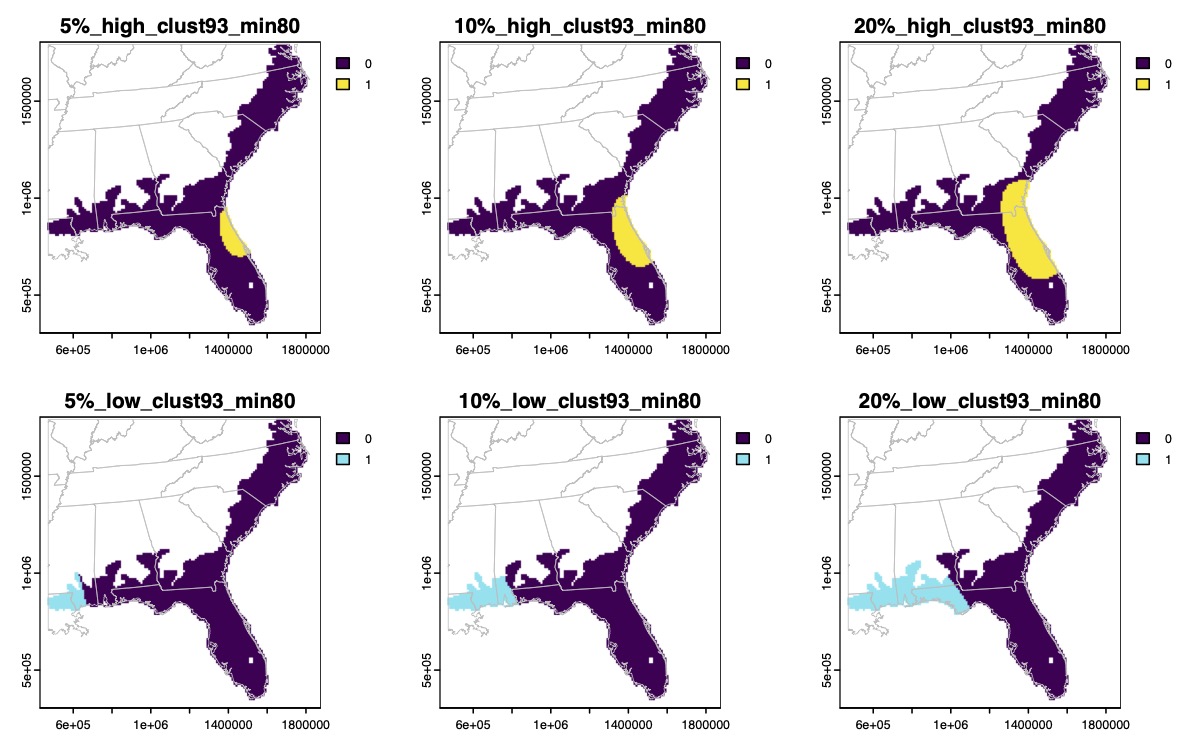

Supplementary Fig. 65.

Binary maps of the highest (yellow) or lowest (blue) genomic diversity for *Hyla femoralis* based on 5%, 10%, or 20% quantile thresholds.

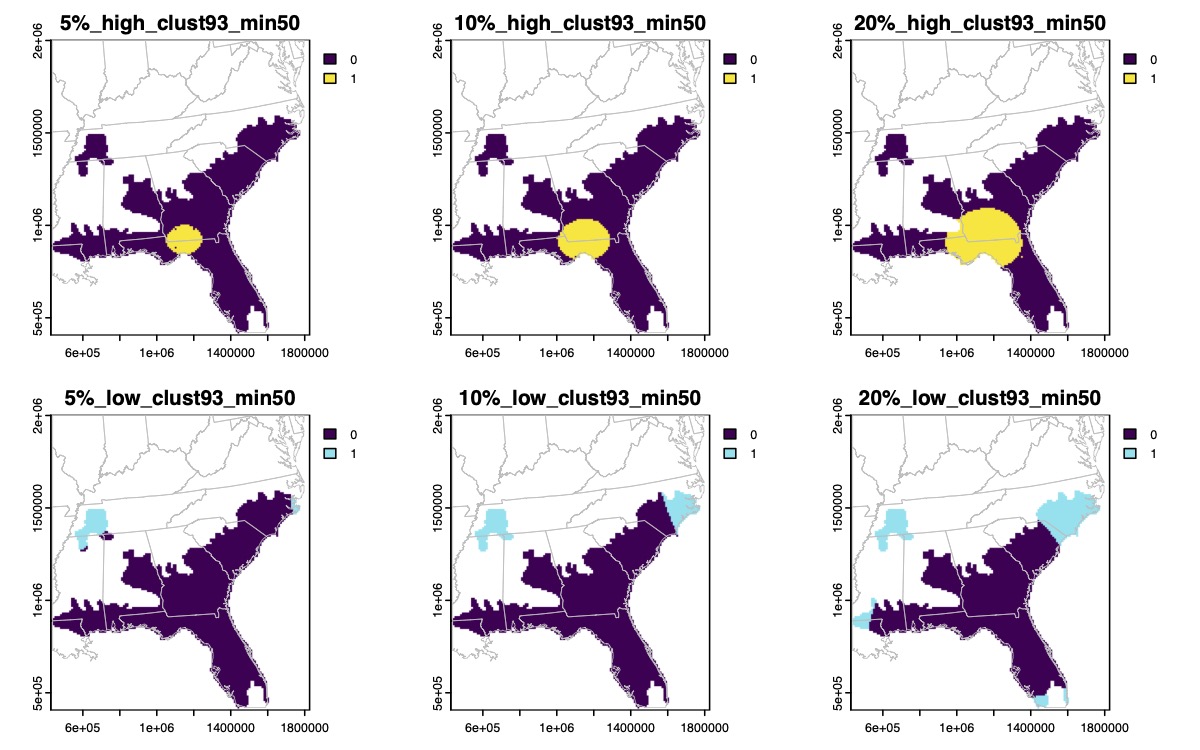

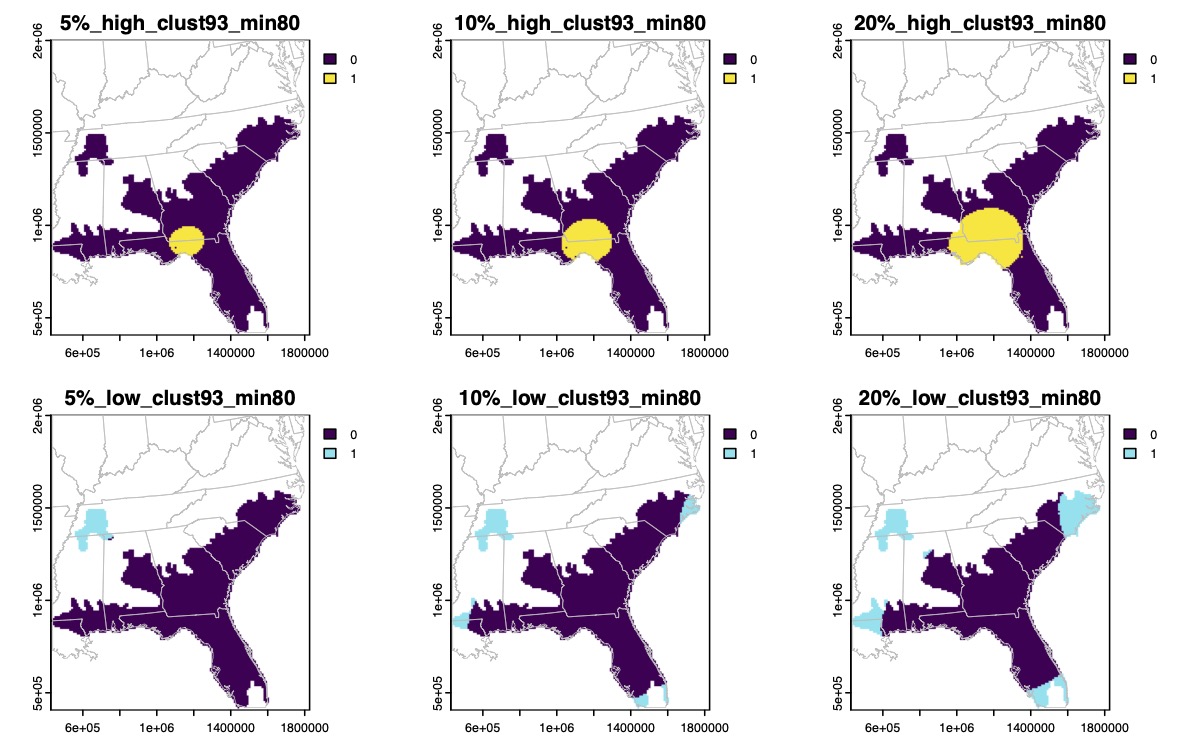

Supplementary Fig. 66.

Binary maps of the highest (yellow) or lowest (blue) genomic diversity for *Hyla gratiosa* based on 5%, 10%, or 20% quantile thresholds.

Supplementary Fig. 67.

Binary maps of the highest (yellow) or lowest (blue) genomic diversity for *Hyla squirella* based on 5%, 10%, or 20% quantile thresholds.

Supplementary Fig. 68.

Binary maps of the highest (yellow) or lowest (blue) genomic diversity for *Incilius nebulifer* based on 5%, 10%, or 20% quantile thresholds.

Supplementary Fig. 69.

Binary maps of the highest (yellow) or lowest (blue) genomic diversity for *Pseudacris clarkii* based on 5%, 10%, or 20% quantile thresholds.

Supplementary Fig. 70.

Binary maps of the highest (yellow) or lowest (blue) genomic diversity for *Pseudacris crucifer* based on 5%, 10%, or 20% quantile thresholds.

Supplementary Fig. 70. (continued)

Binary maps of the highest (yellow) or lowest (blue) genomic diversity for *Pseudacris crucifer* based on 5%, 10%, or 20% quantile thresholds.

Supplementary Fig. 71.

Binary maps of the highest (yellow) or lowest (blue) genomic diversity for *Pseudacris feriarum* based on 5%, 10%, or 20% quantile thresholds.

Supplementary Fig. 71. (continued)

Binary maps of the highest (yellow) or lowest (blue) genomic diversity for *Pseudacris feriarum* based on 5%, 10%, or 20% quantile thresholds.

Supplementary Fig. 72.

Binary maps of the highest (yellow) or lowest (blue) genomic diversity for *Pseudacris fouquettei* based on 5%, 10%, or 20% quantile thresholds.

Supplementary Fig. 72. (continued)

Binary maps of the highest (yellow) or lowest (blue) genomic diversity for *Pseudacris fouquettei* based on 5%, 10%, or 20% quantile thresholds.

Supplementary Fig. 73.

Binary maps of the highest (yellow) or lowest (blue) genomic diversity for *Pseudacris maculata* based on 5%, 10%, or 20% quantile thresholds.

Supplementary Fig. 73. (continued)

Binary maps of the highest (yellow) or lowest (blue) genomic diversity for *Pseudacris maculata* based on 5%, 10%, or 20% quantile thresholds.

Supplementary Fig. 74.

Binary maps of the highest (yellow) or lowest (blue) genomic diversity for *Pseudacris nigrita* based on 5%, 10%, or 20% quantile thresholds.

Supplementary Fig. 75.

Binary maps of the highest (yellow) or lowest (blue) genomic diversity for *Pseudacris ocularis* based on 5%, 10%, or 20% quantile thresholds.

Supplementary Fig. 75. (continued)

Binary maps of the highest (yellow) or lowest (blue) genomic diversity for *Pseudacris ocularis* based on 5%, 10%, or 20% quantile thresholds.

Supplementary Fig. 76.

Binary maps of the highest (yellow) or lowest (blue) genomic diversity for *Pseudacris ornata* based on 5%, 10%, or 20% quantile thresholds.

Supplementary Fig. 76. (continued)

Binary maps of the highest (yellow) or lowest (blue) genomic diversity for *Pseudacris ornata* based on 5%, 10%, or 20% quantile thresholds.

Supplementary Fig. 77.

Binary maps of the highest (yellow) or lowest (blue) genomic diversity for *Pseudacris streckeri* based on 5%, 10%, or 20% quantile thresholds.

Supplementary Fig. 78.

Binary maps of the highest (yellow) or lowest (blue) genomic diversity for *Pseudacris triseriata* based on 5%, 10%, or 20% quantile thresholds.

Supplementary Fig. 78. (continued)

Binary maps of the highest (yellow) or lowest (blue) genomic diversity for *Pseudacris triseriata* based on 5%, 10%, or 20% quantile thresholds.

Supplementary Fig. 79.

Binary maps of the highest (yellow) or lowest (blue) genomic diversity for *Rana berlandieri* based on 5%, 10%, or 20% quantile thresholds.

Supplementary Fig. 80.

Binary maps of the highest (yellow) or lowest (blue) genomic diversity for *Rana blairi* based on 5%, 10%, or 20% quantile thresholds.

Supplementary Fig. 80. (continued)

Binary maps of the highest (yellow) or lowest (blue) genomic diversity for *Rana blairi* based on 5%, 10%, or 20% quantile thresholds.

Supplementary Fig. 81.

Binary maps of the highest (yellow) or lowest (blue) genomic diversity for *Rana catesbeiana* based on 5%, 10%, or 20% quantile thresholds.

Supplementary Fig. 82.

Binary maps of the highest (yellow) or lowest (blue) genomic diversity for *Rana clamitans* based on 5%, 10%, or 20% quantile thresholds.

Supplementary Fig. 83.

Binary maps of the highest (yellow) or lowest (blue) genomic diversity for *Rana grylio* based on 5%, 10%, or 20% quantile thresholds.

Supplementary Fig. 83. (continued)

Binary maps of the highest (yellow) or lowest (blue) genomic diversity for *Rana grylio* based on 5%, 10%, or 20% quantile thresholds.

Supplementary Fig. 84.

Binary maps of the highest (yellow) or lowest (blue) genomic diversity for *Rana palustris* based on 5%, 10%, or 20% quantile thresholds.

Supplementary Fig. 84. (continued)

Binary maps of the highest (yellow) or lowest (blue) genomic diversity for *Rana palustris* based on 5%, 10%, or 20% quantile thresholds.

Supplementary Fig. 85.

Binary maps of the highest (yellow) or lowest (blue) genomic diversity for *Rana pipiens* based on 5%, 10%, or 20% quantile thresholds.

Supplementary Fig. 86.

Binary maps of the highest (yellow) or lowest (blue) genomic diversity for *Rana sphenocephala* based on 5%, 10%, or 20% quantile thresholds.

Supplementary Fig. 86. (continued)

Binary maps of the highest (yellow) or lowest (blue) genomic diversity for *Rana sphenocephala* based on 5%, 10%, or 20% quantile thresholds.

Supplementary Fig. 87.

Binary maps of the highest (yellow) or lowest (blue) genomic diversity for *Rana sylvatica* based on 5%, 10%, or 20% quantile thresholds.

Supplementary Fig. 87. (continued)

Binary maps of the highest (yellow) or lowest (blue) genomic diversity for *Rana sylvatica* based on 5%, 10%, or 20% quantile thresholds.

Supplementary Fig. 88.

Binary maps of the highest (yellow) or lowest (blue) genomic diversity for *Scaphiopus couchii* based on 5%, 10%, or 20% quantile thresholds.

Supplementary Fig. 88. (continued)

Binary maps of the highest (yellow) or lowest (blue) genomic diversity for *Scaphiopus couchii* based on 5%, 10%, or 20% quantile thresholds.

Supplementary Fig. 89.

Binary maps of the highest (yellow) or lowest (blue) genomic diversity for *Scaphiopus holbrookii* based on 5%, 10%, or 20% quantile thresholds.

Supplementary Fig. 89. (continued)

Binary maps of the highest (yellow) or lowest (blue) genomic diversity for *Scaphiopus holbrookii* based on 5%, 10%, or 20% quantile thresholds.

Supplementary Fig. 90.

Binary maps of the highest (yellow) or lowest (blue) genomic diversity for *Scaphiopus hurterii* based on 5%, 10%, or 20% quantile thresholds.

Supplementary Fig. 90. (continued)

Binary maps of the highest (yellow) or lowest (blue) genomic diversity for *Scaphiopus hurterii* based on 5%, 10%, or 20% quantile thresholds.

Supplementary Fig. 91.

Binary maps of the highest (yellow) or lowest (blue) genomic diversity for *Spea bombifrons* based on 5%, 10%, or 20% quantile thresholds.

Supplementary Fig. 91. (continued)

Binary maps of the highest (yellow) or lowest (blue) genomic diversity for *Spea bombifrons* based on 5%, 10%, or 20% quantile thresholds.

Supplementary Fig. 92.

Binary maps of the highest (yellow) or lowest (blue) genomic diversity for *Spea multiplicata* based on 5%, 10%, or 20% quantile thresholds.

Supplementary Fig. 92. (continued)

Binary maps of the highest (yellow) or lowest (blue) genomic diversity for *Spea multiplicata* based on 5%, 10%, or 20% quantile thresholds.

**Supplementary Fig. 93.**

Composite maps of high (“Hotspots”) and low (“Coldspots”) genomic diversity across 46 species at the 5% threshold. (**a**, **b**) The color scale indicates the number of species in each grid cell with their highest (**a**) or lowest (**b**) 5% of genomic diversity. (**c**, **d**) Areas of significant clustering where multiple species have their highest (**c**, orange) or lowest (**d**, blue) 5% of genomic diversity. (**e**, **f**) The color scale indicates the proportion of species in each grid cell with their highest (**e**) or lowest (**f**) 5% of genomic diversity. (**g**, **h**) Areas of significant clustering where multiple species have their highest (**g**, orange) or lowest (**h**, blue) 5% of genomic diversity.

**Supplementary Fig. 94.**

Composite maps of high (“Hotspots”) and low (“Coldspots”) genomic diversity across 46 species at the 10% threshold. (**a**, **b**) The color scale indicates the number of species in each grid cell with their highest (**a**) or lowest (**b**) 10% of genomic diversity. (**c**, **d**) Areas of significant clustering where multiple species have their highest (**c**, orange) or lowest (**d**, blue) 10% of genomic diversity. (**e**, **f**) The color scale indicates the proportion of species in each grid cell with their highest (**e**) or lowest (**f**) 10% of genomic diversity. (**g**, **h**) Areas of significant clustering where multiple species have their highest (**g**, orange) or lowest (**h**, blue) 10% of genomic diversity.

**Supplementary Fig. 95.**

Model results from *MCMCglmm* testing potential predictors of whether species show a latitudinal gradient in genomic diversity based on Spearman correlations (binary response – yes/no). For ecoregions, the effects are relative to Eastern Temperate Forests. Posterior means and 95% credible intervals are combined across four model runs. The 95% credible intervals all overlap with 0, indicating no predictors are significant. Models include 44 species.

**b**

**a**

**Supplementary Fig. 96.**

Model results from *MCMCglmm* testing potential predictors of the correlation strength between latitude and genomic diversity based on Spearman correlations (continuous response – Spearman’s rho). For ecoregions, the effects are relative to Eastern Temperate Forests. Posterior means and 95% credible intervals are combined across four model runs. The 95% credible intervals all overlap with 0, indicating no predictors are significant. (**a**) Reduced models with 25 species that have thermal tolerance information, thus all six predictors are included. (**b**) Full models with 44 species and five predictors.

Supplementary Table 1 (separate file). Detailed sample information including catalogue numbers, localities, and sequence data.

Supplementary Table 2 (separate file). Species-level information and results from correlations and spatial regressions.
